# Spatial Transcriptomics Reveals Region-Specific Host Responses in Amoebic Gill Disease of Atlantic Salmon

**DOI:** 10.64898/2026.10.09.756816

**Authors:** Yee Wan Liu, Bachar Cheaib, Antonella Bacigalupo, Brendan Robertson, Matthew C. Sinton, Juan F. Quintana, Gemma Thomson, Vivienne Morrison, Colin Nixon, Elzbieta Krol, Samuel Martin, Hamish Rodger, Alexander West, Ralph Bickerdike, Fintan Egan, Neil Ruane, Michael Barrett, Martin Llewellyn

## Abstract

Amoebic gill disease (AGD), caused by the parasite *Neoparamoeba perurans,* is a major welfare and economic burden for the Atlantic salmon aquaculture industry. However, the spatial organisation of immune functions within the gill, the site of infection, remains poorly resolved. Here, we present the first spatial transcriptomic atlas of the Atlantic salmon gill affected by AGD, generated using 10x Genomics Visium on four AGD-positive fish, confirmed by gill scoring and quantitative PCR for parasite load. As no uninfected gills were included, this study maps the anatomical and immunological organisation of AGD-affected gills rather than defining AGD-specific pathology. Unsupervised clustering of spatial gene expression resolved distinct transcriptional domains that grouped into four principal anatomical compartments of the gill, including (i) epithelial and lamellar regions defined by keratin and osmoregulatory gene expression, (ii) gill arch regions enriched for connective tissue and smooth muscle markers, (iii) an erythroid-associated vascular cluster, and (iv) a skeletal muscle cluster with signatures of glycolysis and oxidative phosphorylation. Across these domains, we identified a spatially organised and diffused immune landscape, with pattern recognition receptors, antimicrobial effectors and chemokines concentrated at the filament surface, and immunoglobulin light chain transcripts consistent with B lymphocyte activity enriched toward the gill arch. Integration of the spatial data with a published single-nucleus reference atlas using reference-based deconvolution resolved the cellular identity underlying these domains and revealed frequent co-localisation of immune and structural cell types, suggesting regulated cell to cell communication within the gill-associated lymphoid tissue. Overall, our results show that the Atlantic salmon gill is not a transcriptionally uniform surface but a regionally specialised organ, with immune functions distributed across, and specialised within, distinct anatomical compartments

## 1 Introduction

Amoebic gill disease (AGD), caused by *Neoparamoeba perurans* (Young *et al*., 2007), is marked by severe lesions in the branchial epithelium of salmon, which compromise gas exchange and lead to respiratory distress, reduced feeding, impaired growth and elevated mortality (Marcos-López and Rodger, 2020; Rozas-Serri, 2019). These clinical outcomes not only raise major fish welfare concerns but also result in substantial economic losses due to decreased productivity and the need for repeated, costly treatments (Kube *et al*., 2012; Power *et al*., 2025). To effectively tackle the economic impact and welfare issues caused by AGD, it is necessary to move beyond conventional diagnostic approaches such as macroscopic gill scoring and histology (Adams *et al*., 2004; Taylor *et al*., 2009).

High-resolution gene expression analyses, including transcriptomics, offer a detailed view of the underlying molecular mechanisms, enabling the discovery of critical biomarkers and therapeutic targets that traditional methods cannot identify (Botwright *et al*., 2021; Gjessing *et al*., 2020). These approaches provide comprehensive molecular and mechanistic insights into host-pathogen interactions and cellular responses, thereby facilitating improved disease management and the development of targeted interventions (Botwright *et al*., 2021).

The conventional approach to gene expression profiling, bulk RNA sequencing (RNA seq), provides an averaged transcriptomic profile from a heterogeneous tissue or cell population (Lowe *et al*., 2017). This approach allows for identifying differentially expressed genes and reveals general molecular functions across samples. For instance, Botwright et al. (2020), using a dual RNA-seq approach, simultaneously profiling the transcriptomes of both the Atlantic salmon host and the parasite *N. perurans* from gill tissue from both lesioned and non-lesioned areas, head kidney, and spleen from AGD-affected and naïve salmon. Their analysis successfully identified key molecular interactions, including the alteration of host gene networks by specific transcription factors and potential parasite virulence genes (Botwright *et al*., 2021). While such an approach offers valuable molecular insights, it averages the transcriptomic signals across all cells, thereby obscuring the distinct contributions of individual cell types and leaving key questions unanswered, such as which cells specifically orchestrate the immune response within the host.

To address the limitation of the loss of cellular heterogeneity, single-cell RNA sequencing (scRNA-seq) emerged, enabling transcriptomic profiling at single-cell resolution (Kashima *et al*., 2020). In this method, individual cells are isolated from heterogeneous samples, through the means of microfluidic droplet encapsulation, along with barcoded primers and reagents (Kashima *et al*., 2020). Cell lysis and reverse transcription occur within each droplet, during which a unique DNA sequence, known as a barcode, is incorporated into the cDNA (Kashima *et al*., 2020). This allows sequencing reads to be traced back to their cell of origin, enabling the precise reconstruction of the transcriptome of each cell. For instance, Gervais et al. (2023) used single-cell RNA sequencing on Atlantic salmon SHK-1 cells infected with ISAV to capture cell-specific transcriptional changes during infection. They revealed distinct antiviral states and paracrine signalling in bystander cells, which could otherwise be masked by traditional bulk RNA sequencing (Gervais *et al*., 2023). Such ability to dissect cellular diversity and dynamic states makes single-cell transcriptomics a “gold standard” tool (Andresen *et al*., 2024), especially in complex biological systems such as immune tissues where traditional methods often obscure critical subpopulations or transient cellular activities (Efremova *et al*., 2020).

While scRNA-seq has dramatically enhanced the resolution of transcriptomic analysis, its dependence on fresh, intact cells presents limitations, particularly with challenging or archived samples, where dissociation, such as those caused by enzymatic digestion, can compromise cell integrity or introduce artifacts (Denisenko *et al*., 2020). In such scenarios, single nuclei transcriptomics (snRNA-seq) emerges as a powerful alternative, while providing high-resolution gene expression data from single nuclei by circumventing the drawbacks of tissue dissociation, making it more suitable for fragile, complex, or preserved specimens (Denisenko *et al*., 2020; Grindberg *et al*., 2013; McLaughlin *et al*., 2022). Such an advantage is highlighted by studies such as West et al. (2021), who used snRNA-seq on Atlantic salmon gills and revealed a dramatic reduction in various immune-related cell types during smoltification. Furthermore, Andresen et al. (2024) directly compared both methods, snRNA-seq and scRNA-seq, on Atlantic salmon head kidney, revealing cellular heterogeneity and developmental trajectories within B cells, T cells, and endothelial cells, and compiling a comprehensive list of marker genes for immune-relevant cell populations. Their study also highlighted that snRNA-seq was more robust for identifying certain cell types and subpopulations, which appeared to be lost during the preparation of scRNA-seq suspensions (Andresen *et al*., 2024). While snRNA-seq is adept at capturing nuclear-specific transcripts and transcription factors, which are important for cell identity, the full spectrum of mRNA, including those involved in immediate cellular functions and rapid changes during differentiation, is better represented in whole-cell transcriptomes using scRNA (Andresen *et al*., 2024).

Despite the high cellular resolution presented by both scRNA-seq and snRNA-seq, they lack spatial context (Rao *et al*., 2021). Such spatial context is important, as cell-cell interactions within the complex tissue microenvironment are crucial for development (Xia *et al*., 2019) and disease progression (Pham *et al*., 2023). To overcome this challenge, spatial transcriptomics (ST) technologies have emerged, combining gene expression profiling with preserved tissue architecture (Moses and Pachter, 2022; Williams *et al*., 2022). To date, only two ST have been performed on *Salmo salar*: one on muscle (Bjørgen *et al*., 2024) and another on its skin (Sveen *et al*., 2023). Bjørgen *et al*. (2024) investigated red and melanized muscle lesions in farmed Atlantic salmon, identifying genes involved in hypoxia and immune responses as likely responsible for muscle discoloration and inflammation. Sveen et al. (2023) identified tissue-specific gene markers, such as abundant collagen type 1 in the skin and keratins in epithelial tissue, highlighting the diverse cell types and gene expression patterns across various skin layers. As highlighted in these initial *S. salar* studies, spatial transcriptomics retains the anatomical context of gene expression, thereby bridging the gap between high-resolution molecular data and histological structure.

Indeed, sequencing-based ST methods, such as 10x Genomics Visium, are powerful tools for investigating how cellular states and interactions are shaped by their spatial environment; however, they lack single-cell or nuclear resolution (Sun *et al*., 2018). To address this caveat, computational methods, such as the Robust Cell Type Decomposition (RCTD), are designed to integrate single-cell RNA-seq data with spatial transcriptomics data (Cable *et al*., 2022; Sun *et al*., 2018). RCTD can effectively decompose the mixed gene expression signals from spatial transcriptomics spots, each of which may contain multiple cells, into proportions of the different cell types (Cable *et al*., 2022; Li, Zhang, *et al*., 2022). This can be achieved by leveraging cell type profiles learned from scRNA-seq data and correcting for differences between sequencing technologies (Cable *et al*., 2022).

Our study aims to characterise the spatial transcriptomic architecture of the Atlantic salmon gill during amoebic gill disease, resolving how epithelial, vascular, muscular and immune compartments are organised *in situ* in response to *N. perurans* infection. To achieve this, we: (i) applied 10x Genomics Visium spatial transcriptomics to gill tissue using four AGD-confirmed specimens; (ii) defined tissue domains through unsupervised clustering; and (iii) resolved the cellular composition underlying these domains to map immunologically relevant cell populations across the gill-associated lymphoid tissue and characterise their spatial co-localisation patterns.

## 2 Materials and methods

### 2.1 Fish Origin and Ethical Approval

A mixed-sex Irish strain of Atlantic salmon (*Salmo salar*) was obtained as eyed eggs from a commercial supplier (Mowi Ireland). Fish were reared under standard husbandry conditions at the Marine Institute’s Newport Research Facility (County Mayo, Ireland) using a flow-through freshwater system until smoltification. Post-smolts were transferred to the Marine Institute’s Lehanagh Pool Marine Research Site in Bertraghboy Bay, Connemara, Ireland, for the marine trial phase. The site operates under an aquaculture licence and adheres to organic farming standards, with no use of prescription medicines or antifoulants. All experimental procedures involving animals were conducted in accordance with the European Union Directive 2010/63/EU and its Irish transposition (S.I. No. 543 of 2012, as amended). Ethical approval was granted by the Republic of Ireland (ROI) Health Products Regulatory Authority (HPRA), authorisation number AE19121/I035, in collaboration with the Marine Institute ROI at the Lehanagh Pool Marine Research Site.

### 2.2 Amoebic Gill Disease Gill Scoring and Molecular Confirmation

To validate the AGD status on *S. salar*, fish were macroscopically assessed for AGD severity using the gill scoring system described by Taylor et al. (2016). Gill swabs were also taken from these fish for subsequent DNA extraction using DNA QIAamp Mini Kit (Cat No 51306, QIAGEN) and qPCR to quantify the amoeba load using the method established by Downes et al. (2015).

### 2.3 Tissue Embedding, Cryopreservation and Cryosection for spatial transcriptomics

Following gill excision, tissues were gently blotted to remove excess moisture and embedded in Optimal Cutting Temperature (OCT) compound (KMA-0100-00A, CellPath), a water-soluble medium that preserves tissue structure during sectioning. Samples were carefully oriented in cryomolds (E6032, Peel-A-Way™ embedding molds, Sigma Aldrich), then rapidly frozen by immersion in isopentane pre-cooled in liquid nitrogen to approximately –150 °C. This rapid freezing minimised ice crystal formation and preserved both morphology and RNA quality.

Four fish gill samples with the highest morphological integrity and RNA Integrity Number scores were selected (**Supplementary Note 1** and **Supplementary Note 2**). Each sample was oriented and mounted onto a cryostat chuck using OCT compound. Tissue sections were cut at 10 μm thickness using a Leica CM1950 cryostat, with sections placed onto one 10x Genomics Visium permeability slide and two 10x Genomics Visium gene expression slides as technical replicates for downstream spatial transcriptomic analysis. The cryostat chamber was maintained at -18 °C during sectioning. After cutting, the sections designated for spatial analysis were initially stored at -20 °C, then transferred to -80 °C for long-term storage.

### 2.4 Tissue Fixation, Hematoxylin and Eosin Staining and Imaging

Hematoxylin and Eosin staining for spatial transcriptomics was performed as described in the 10x Visium protocol CG000160, Rev D. Briefly, slides were fixed using pre-chilled Methanol (Millipore Sigma, for HPLC, ≥ 99.9%, product number 34860). Following fixation, slides underwent sequential staining with Hematoxylin, Mayer’s (Agilent, S330930-2), bluing with Dako Bluing Buffer (Agilent, CS70230-2), and counterstaining with Dako Eosin (Agilent, CS70130-2). This was followed by a dehydration step using Isopropanol (2-Propanol ≥ 99.5%, Millipore Sigma, l9516-25ML) to remove water prior to the subsequent staining steps and final air drying, thereby preparing the tissue for imaging and the downstream 10X Genomics Visium Spatial Gene Expression workflow. Stained slides were imaged using an Olympus BX63 Single Slide-Scanner, employing stitching for complete tissue section capture.

### 2.5 Library preparation and Sequencing

Once the permeabilisation time was determined using the protocol detailed in 10x Genomics Document Number CG000238, Rev E, the library was constructed following the protocol described in 10x Genomics Document Number CG000239, Rev D. Briefly, each capture area underwent 6 minutes of tissue permeabilisation, followed by a series of enzymatic reactions that generated cDNA from RNA, and ultimately led to the construction of dual-index libraries (**Supplementary Note 3**). Dual-indexed 10x Visium Spatial Gene Expression libraries were sequenced on an Illumina NovaSeq X Plus platform with paired-end 150 bp reads, yielding 72 Gb of raw data per library, followed by demultiplexing.

### 2.6 Image Processing

Brightfield images from each capture area were processed using the 10x Genomics Loupe Browser (v6.0.0). The analysis included two 10x Genomics Gene Expression slides (slide number V13U19294 and V13U19334), each with four designated capture areas labelled A, B, C, and D. Spatial barcodes corresponding to spots with underlying tissue were manually selected and exported. These barcodes were then used as a filtering step to retain only tissue-covered spots for downstream spatial gene expression analysis.

### 2.7 Genome Index Preparation for Spatial Transcriptomics

A compatible reference genome was generated using the mkref function from cellranger (v 7.0.0.), which requires both a genome assembly and a corresponding annotation file in GTF format. The Atlantic salmon genome assembly (Ssal_v3.1, GCF_905237065.1, Ensemble release 112), containing over 65,000 genes and pseudogenes, including nearly 43,000 protein-coding genes, was used along with its annotation file.

### 2.8 Alignment and Gene Quantification

Reads were aligned to the reference genome previously generated using the spaceranger (v 3.0.1) software. Spaceranger is a pipeline that performs read demultiplexing, alignment, and gene counting specifically designed for 10x Genomics spatial transcriptomics data, enabling the assignment of reads to spatial locations on the tissue section.

### 2.9 Data Filtration, Transformation, Dimensionality reduction and Clustering in Seurat

The resulting filtered gene expression matrices from Space Ranger were loaded into R (v 4.5.1) as a Seurat object using the Load10X_Spatial function from the Seurat R package. This step imports the gene expression data along with spatial coordinates and associated image data, providing the foundation for downstream spatial analysis. Empty spots, which lacked tissue coverage, were filtered out manually with loupe browser to reduce noise in the downstream analysis. The filtered data were then subjected to SCTransform, a variance-stabilising transformation implemented in the Seurat R package, which normalises the data whilst accounting for the relationship between gene expression mean and variance, leading to improved identification of variable features. Principal component analysis (PCA) was subsequently performed, which reduces the dimensionality of the data by identifying principal components that capture the main sources of variation in gene expression. The number of dimensions to retain was determined by identifying the elbow point in the elbow plot (**Supplementary Note 4**). Subsequently, Harmony, an algorithm designed to integrate data from multiple samples whilst minimising batch effects, was run to account for any remaining technical variability between the technical replicates. UMAP spatial spots dimensionality reduction was then performed using the RunUMAP function from the Seurat package. Following dimensionality reduction with PCA, a k-nearest neighbours (k-NN) graph was computed using the FindNeighbors function in Seurat. From this, a Shared Nearest Neighbour (SNN) graph was constructed, which improves clustering robustness by accounting for the number of shared neighbours between spots. Clustering was then performed with the FindClusters function of Seurat, grouping spots with similar gene expression profiles at a resolution of 0.3. A resolution of 0.3 was selected as optimal on the basis that the spatial UMAP and slide visualisations were consistent with expected biological organisation.

### 2.10 Statistics After Clustering

To identify genes that distinguish distinct spatial regions transcriptionally, several analyses were carried out. Differentially expressed genes (DEGs) were identified using the FindMarkers function in Seurat (v5.3.0.), which applies the Wilcoxon Rank-Sum test to compare the expression of each gene within a given cluster against all other clusters. This method detects DEGs without considering spatial context. Marker genes representative of each cluster were further identified using the FindAllMarkers function of Seurat to characterise transcriptional profiles across spatial domains. To capture genes exhibiting spatial heterogeneity, the FindSpatiallyVariableFeatures function of Seurat was applied independently to each sample. This method highlights genes whose expression patterns vary across spatial coordinates, enabling the identification of spatially variable genes (SVGs). Genes identified as both DEGs and SVGs were considered of particular interest, as they reflect both transcriptional and spatial specificity. Finally, only genes that were identified by both FindAllMarkers and FindSpatiallyVariableFeatures were included in downstream correlation analyses. Average log fold-change values were calculated for these genes across clusters and used to generate a Pearson correlation matrix, which was hierarchically clustered to assess inter-cluster relationships.

### 2.11 Integration with snRNA-seq data from West et al. (2021)

To allow integration between spatial transcriptomics data aligned to the Ssal_v3.1 Atlantic salmon genome and a previously published snRNA-seq dataset from West et al. (2021), which was aligned to the ICSASG_v2 genome, a genome coordinate conversion was performed with BCFtools/liftover (Genovese et al., 2024). This conversion tool mapped gene identifiers from ICSASG_v2 to their corresponding locations in the Ssal_v3.1 reference, allowing for consistent comparison across datasets. BCFtools/liftover also enabled the conversion of ENTREZ gene IDs used in the original snRNA-seq data to ENSEMBL gene IDs used in the spatial transcriptomics data aligned with Ssal_v3.1. Robust Cell Type Decomposition (RCTD) (Cable *et al*., 2022) was then applied to the spatial transcriptomics data to estimate cell-type composition at each spatial spot. The West *et al*. (2021) snRNA-seq dataset, annotated with immune cell populations, served as the reference. This allowed spatial spots to be scored based on the transcriptional similarity to known immune cell types, facilitating the identification of spatial patterns of immune cell infiltration within the gill tissue.

## 3 Results

### 3.1 Validation of AGD clinical severity in samples prior to spatial transcriptomic analysis

Gill samples from Atlantic salmon showed variable Amoebic Gill Disease (AGD) severity. Gross gill scoring (**Figure 1**) and qPCR (**Figure 2**) consistently identified Samples A and B as severe, while Samples C and D were classified as mild. Although detailed histological interpretation of AGD was limited by cryopreservation artefacts (**Supplementary Note 5**), the concordance between macroscopic gill lesions and parasite load supported the classification of these four gill samples as AGD-positive. These four samples were processed using 10X Visium spatial transcriptomics, yielding high-quality, deeply sequenced datasets with strong QC metrics suitable for reliable spatial gene expression mapping (**Supplementary Note 6**). Because all four gills were AGD-positive and no uninfected gills were included, the following sections do not characterise AGD pathology or identify disease markers. Instead, they describe the spatial gene expression underlying the gill’s anatomical organisation and the distribution of immune functions across these anatomical regions within AGD-affected gills.

**Figure 1.**
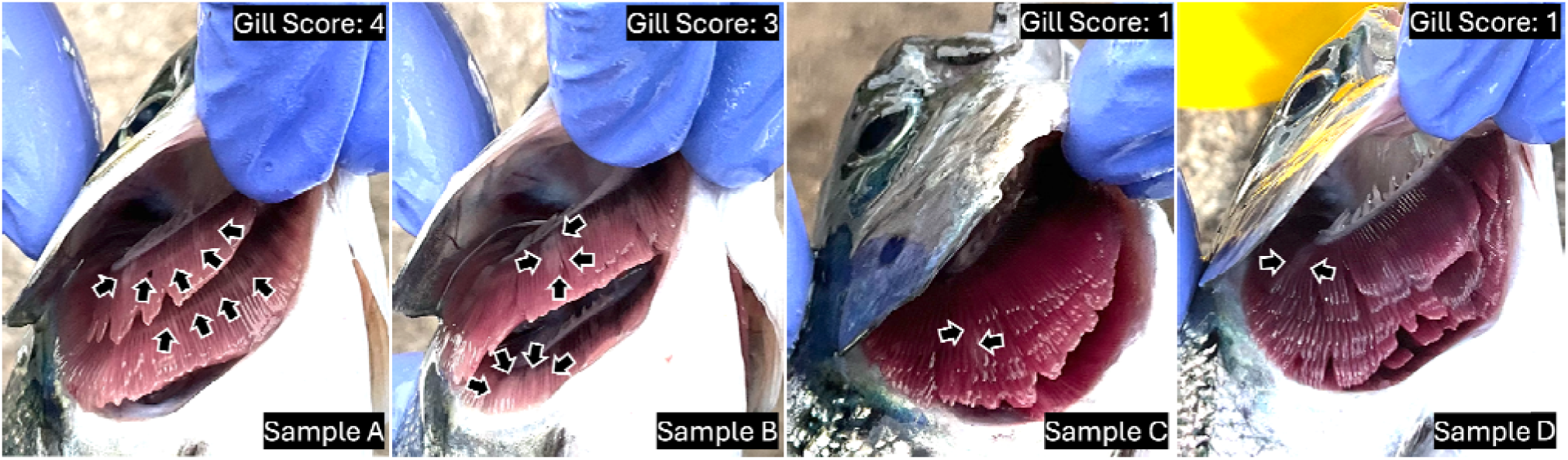
Macroscopic examination of gill lesions in four Atlantic salmon (*Salmo salar*) specimens (Samples A, B, C, and D) selected for cryopreservation and subsequent spatial transcriptomics analysis. White patches indicative of amoebic gill disease (AGD) are visible on the gill surface and are marked with arrows. Lesions were scored using a standard AGD gill scoring system, which ranges from 0 to 5 based on the extent and severity of visible lesions. Score 0: No visible lesions; Scores 1 to 2: Mild to moderate focal white patches on the gill filaments; Scores 3 to 4: Moderate to extensive white patches, often accompanied by increased mucus production and lamellar thickening; Score 5: Severe and widespread lesions with significant gill damage. Samples A and B received higher gill scores (3–4), consistent with severe AGD, while Samples C and D were scored lower (score 1), indicating milder clinical presentation.

**Figure 2.**
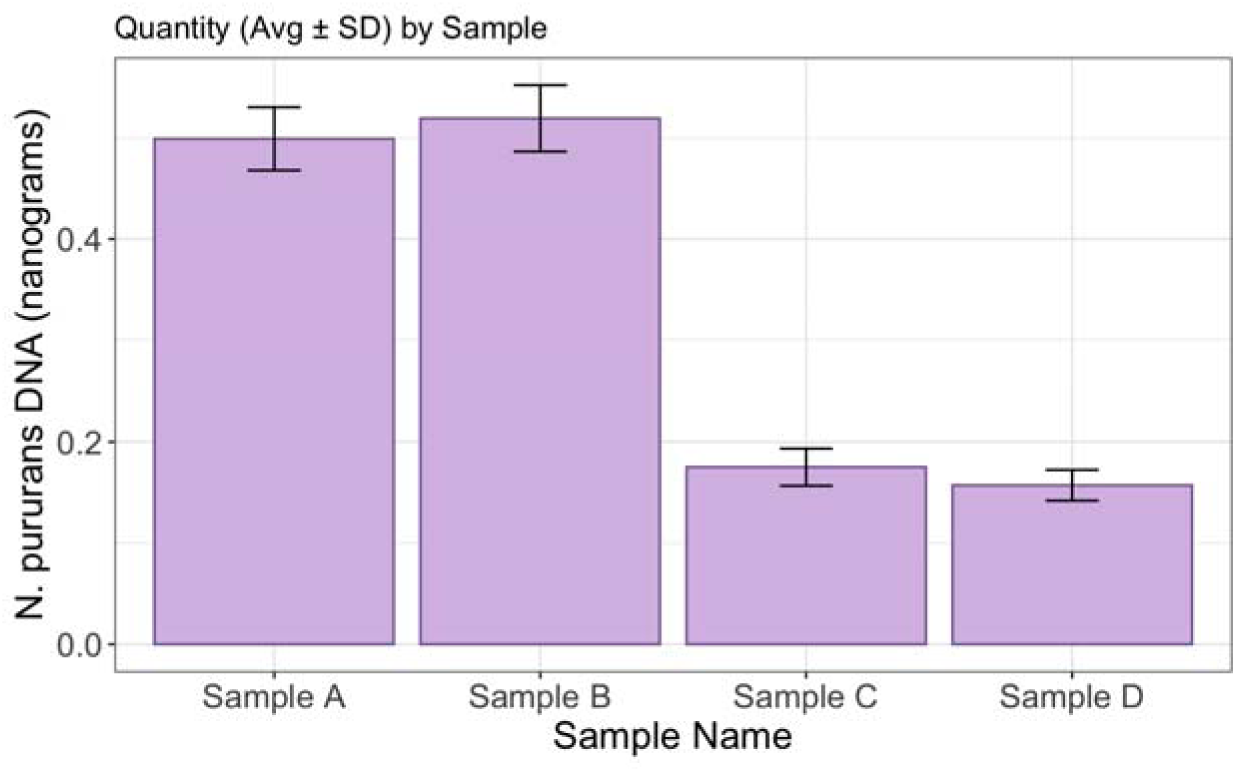
Quantification of *Neoparamoeba perurans* DNA in gill samples from four Atlantic salmon (*Salmo salar*) using quantitative PCR (qPCR), following the protocol described by Downes *et al*. (2015). The x-axis shows the four fish samples analysed (A to D), while the y-axis indicates the average estimated parasite load per sample, expresse as nanograms of *N. perurans* DNA. Error bars denote the standard deviation of three technical replicates per sample.

### 3.2 Anatomical overview of the fourteen spatial gene expression clusters identified across samples

Unsupervised clustering performed, using a shared nearest neighbour (SNN) modularity optimisation algorithm across all spots from the eight samples, had grouped spatial spots with similar transcriptomes. This resulted in 14 distinct spatial clusters (**Figure 3**), with the number of spatial spots for each cluster detailed in **Table 1**.

**Figure 3:**
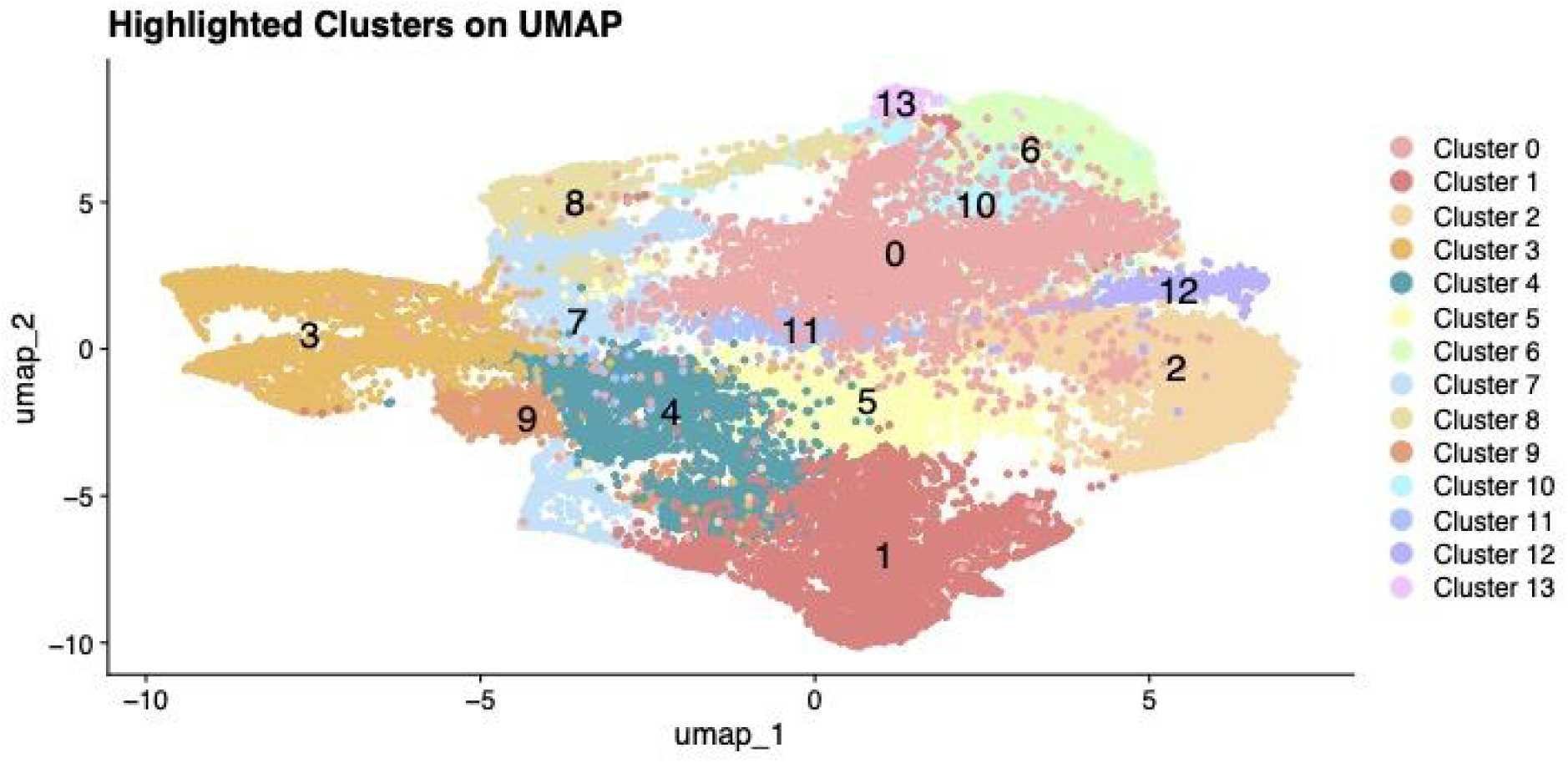
UMAP projection of spatial transcriptomic spots from the gill of Salmo salar, revealing 14 distinct clusters: Cluster 0 (•), 1 (•), 2 (•), 3 (•), 4 (•), 5 (•), 6 (•), 7 (•), 8(•), 9(•), 10(•), 11(•), 12(•), and 13(•). These clusters were identified using the Louvain algorithm at a resolution of 0.3. Clustering was performed following PCA-based dimensionality reduction, construction of a k-nearest neighbours (k-NN) graph, and generation of a Shared Nearest Neighbour (SNN) graph (Waltman and Van Eck, 2013).

**Table 1.** The number of spatial spots across 8 capture areas (4 samples with technical duplicates) for each cluster.

| Cluster | 0 | 1 | 2 | 3 | 4 | 5 | 6 | 7 | 8 | 9 | 10 | 11 | 12 | 13 |
| --- | --- | --- | --- | --- | --- | --- | --- | --- | --- | --- | --- | --- | --- | --- |
| No. of spatial spots | 4,257 | 3,767 | 3,474 | 2,836 | 2,597 | 2,405 | 2,076 | 1,600 | 1,478 | 863 | 778 | 555 | 458 | 212 |

Following the unsupervised clustering, Wilcoxon Rank-Sum tests were performed to compare each cluster against all others, identifying 4,276 differentially expressed genes (DEGs; adjusted p < 0.05, |log FC| > 1). In parallel, spatial analysis using Moran’s I and mark variogram methods identified an average of 678 top spatially variable genes (SVGs) across all 8 captured areas on the 10X Visium slides. Of these, a total of 649 genes were identified as both SVGs and DEGs (**Figure 4**), suggesting these genes hold both spatial and potential functional importance within the gill tissue. These overlapping DEGs and SVGs were subsequently included in the Pearson correlation analysis using average log fold change values (**Figure 5**; tables with the top ten genes per cluster are detailed in the **Supplementary Note 7**). Subsequent hierarchical clustering revealed distinct tissue specific groupings, comprising the filament/lamellae clusters (cluster 0, 2, 5, 6, 10, 11, 12, and 13) and gill arch clusters (cluster 4, 7, 8, and 9), alongside two clusters with distinct transcriptomic profiles: the hemoglobin cluster (cluster 1) and the skeletal muscle cluster (cluster 3), as visualised in the heatmap (**Figure 5**).

**Figure 4.**
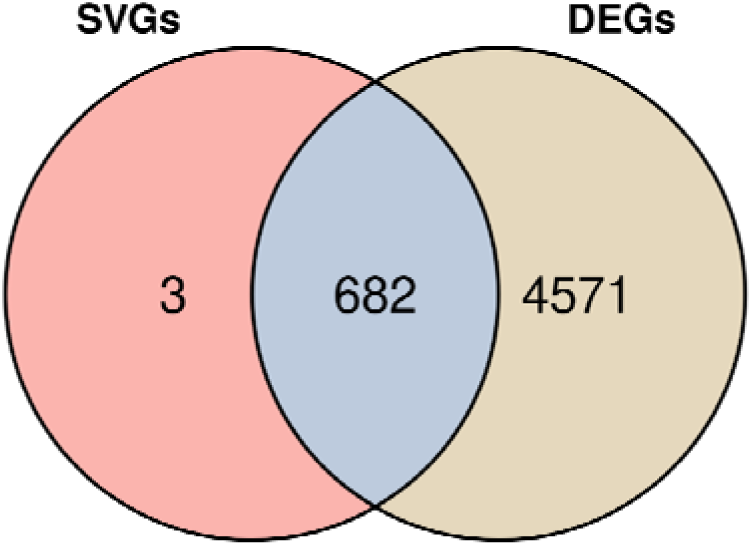
A Venn Diagram illustrates the overlap between genes identified as the top 685 spatially variable genes (SVGs) found across samples and those differentially expressed genes (DEGs) defining specific cluster expression states (with an adjusted p-value < 0.05 and a log2 fold change greater than 1 or less than -1).

**Figure 5:**
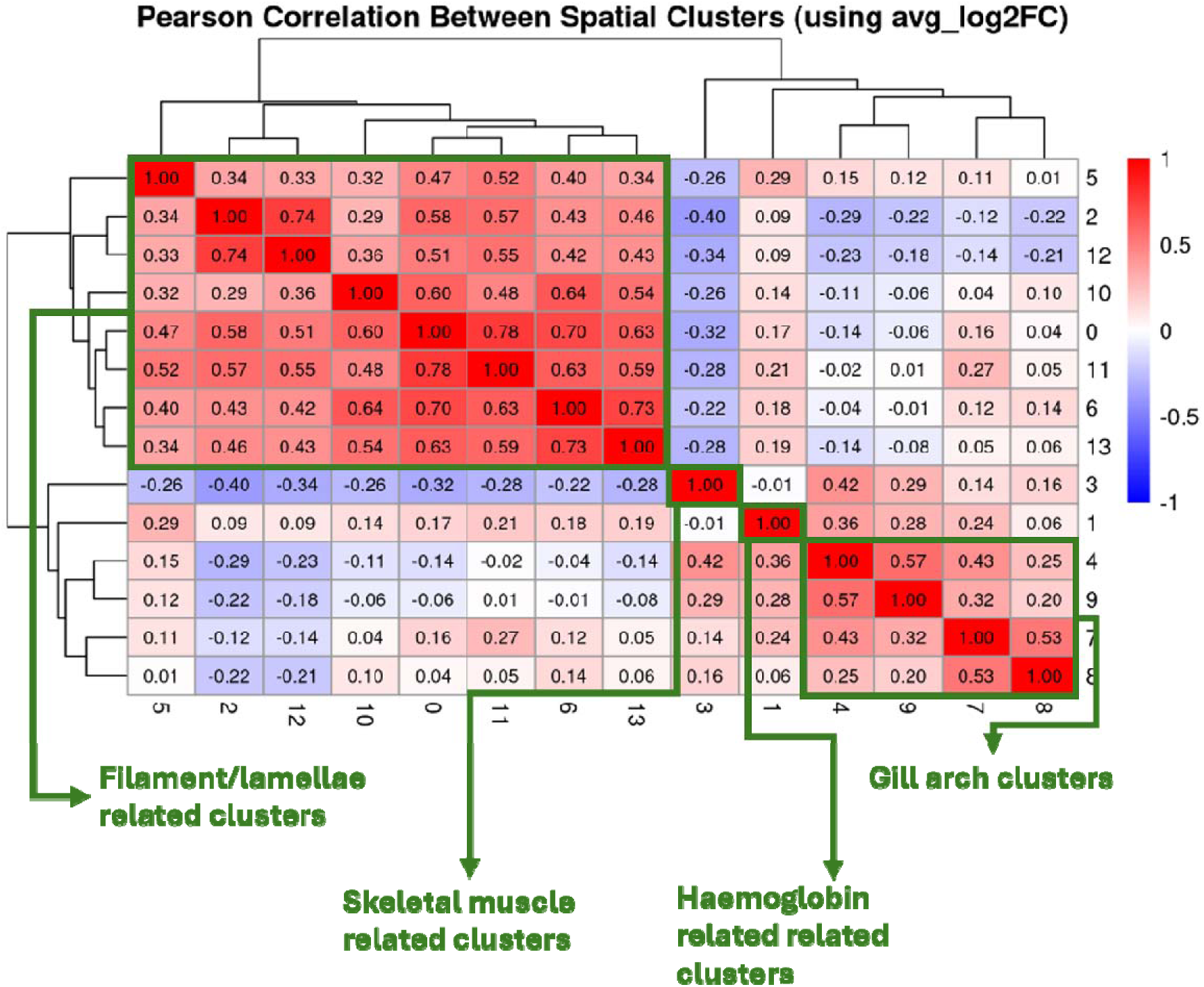
Inter-cluster relationships revealed by Pearson correlation of Marker Gene Expression. A heatmap illustrating the Pearson correlation coefficients between the average log2 fold change (avg_log2FC) of genes that were identified as top markers (adjusted p-value < 0.05 and log2 fold change > 1 from the FindMarkers analysis) and were also among the top 685 spatially variable genes. The avg_log2FC for each gene in a given cluster was calculated relative to all other clusters. Clusters are organised by hierarchical clustering (Euclidean distance and average linkage) applied to both rows and columns, visually grouping clusters with similar marker gene expression profiles. The colour intensity ranges from dark blue (Pearson correlation = -1, strong negative correlation) to dark red (Pearson correlation = +1, strong positive correlation), with white representing a correlation near zero. Numerical values within each cell indicate the exact Pearson correlation coefficient. This visualisation facilitates the identification of highly correlated clusters that may represent similar biological domains or cell types as outlined in green.

#### 3.2.1 Filament and lamellae clusters

Eight clusters (0, 2, 5, 6, 10, 11, 12, and 13) consistently positioned on the gill filaments and lamellae across all samples (**figure 6).** Their transcriptional signatures support core physiological functions, specifically epithelial integrity, active extracellular matrix (ECM) remodeling, and osmoregulation. Epithelial integrity and ECM remodeling are maintained by a structural framework of intermediate filaments and dynamic protein synthesis. For instance, elevated expression of keratins (e.g., *krt15* (**Figure 7a**), and ENSSSAG00000048372 (**Figure 7b**) provides mechanical resilience to exposed surfaces. Meanwhile, ECM remodeling, the process of degrading and reorganising the cellular surroundings, is driven by specialised genes, such as *col10a1b* (**Figure 7c**), a specialised collagen involved in structural scaffold formation, and ***spp1*** (**Figure 7d**), a matricellular protein mediating cell-matrix interactions and tissue remodelling signals. Osmoregulation, the maintenance of internal salt and water balance, is localised primarily within Clusters 12 and 2. This function is facilitated by ATP-dependent ion transporters and ion channels within chloride cells. Key examples include the ENSSSAG00000055927 (**Figure 7e**), which regulates acid-base balance, and *clic1* (**Figure 7f**), which facilitates chloride transport. Additional markers such as *trpv6* (**Figure 7g**) contribute essential calcium flux and osmosensory functions. While these markers define the core functions of these regions, additional related genes identified in these filament and lamellae clusters are detailed in the **Supplementary Note 8**.

**Figure 6:**
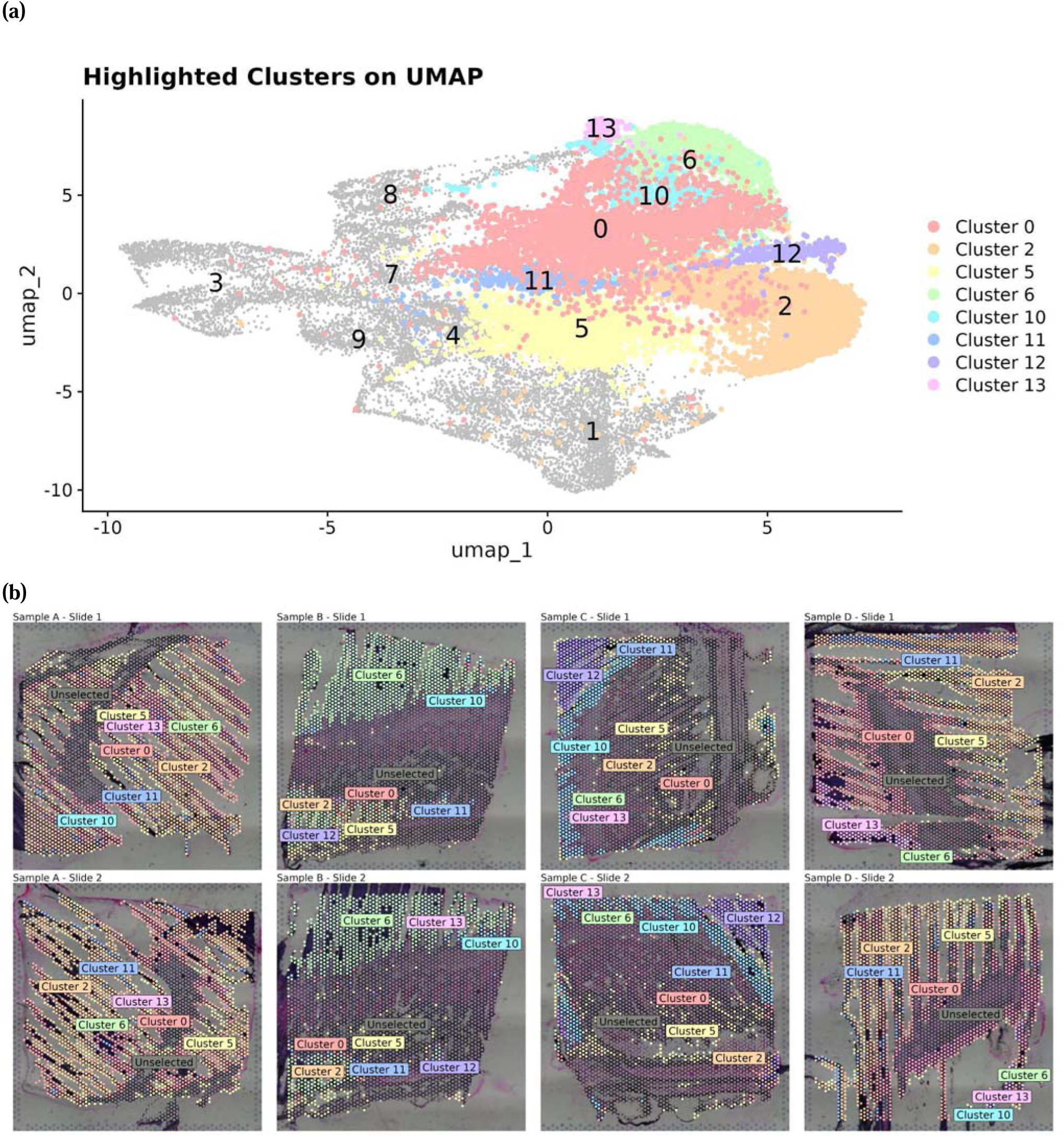
(a) UMAP projection subset highlighting eight transcriptionally similar clusters, 0 (•), 2(•), 5(•), 6(•), 10(•), 11(•), 12(•), and 13(•), that collectively define as the filament and lamellae clusters. These clusters were identified using the Louvain algorithm at a resolution of 0.3, following PCA based dimensionality reduction and construction of a Shared Nearest Neighbour (SNN) graph. Corresponding spatial mappings are shown in 8 panels in (b) with the corresponding sample name above each panel.

**Figure 7.**
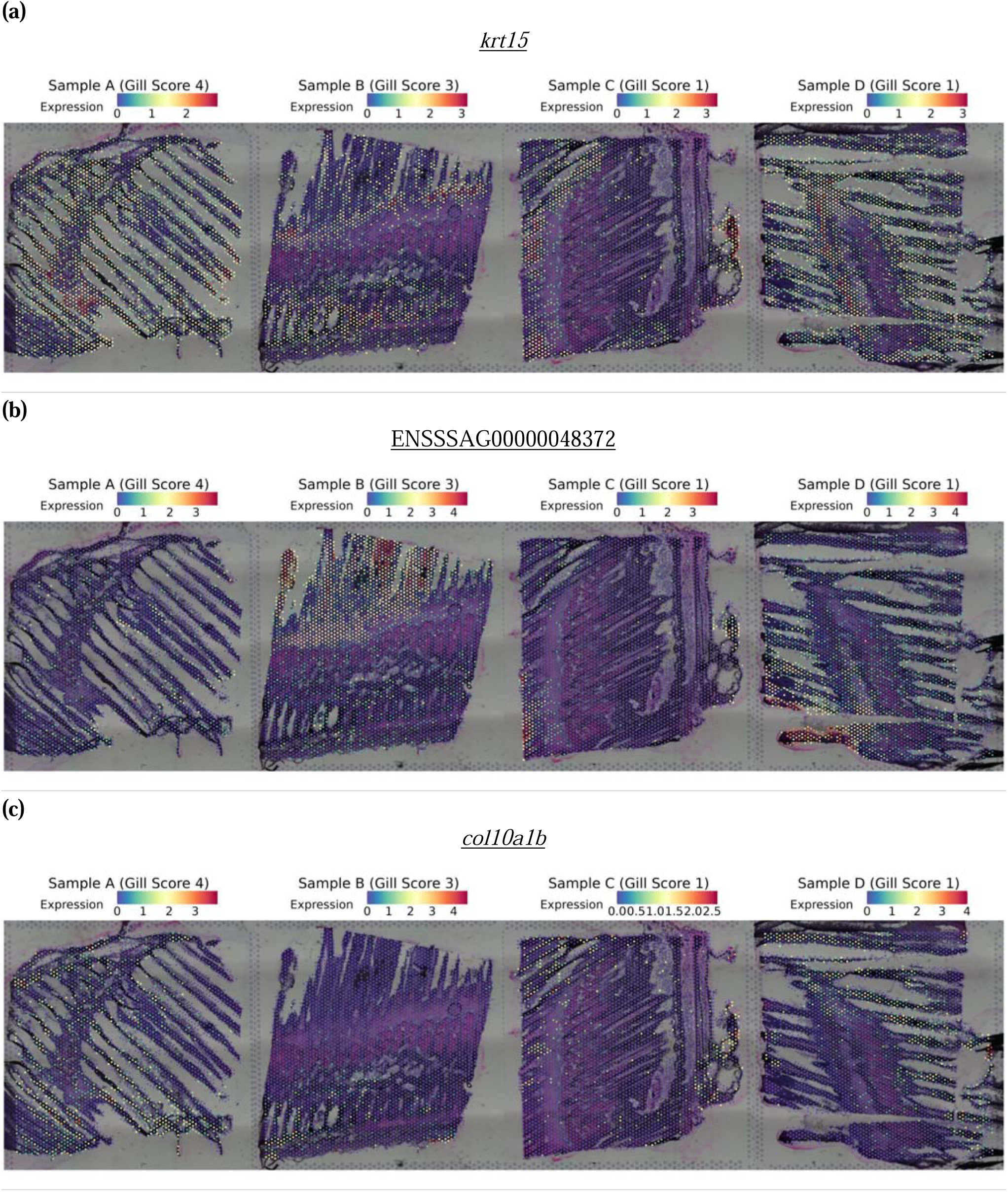
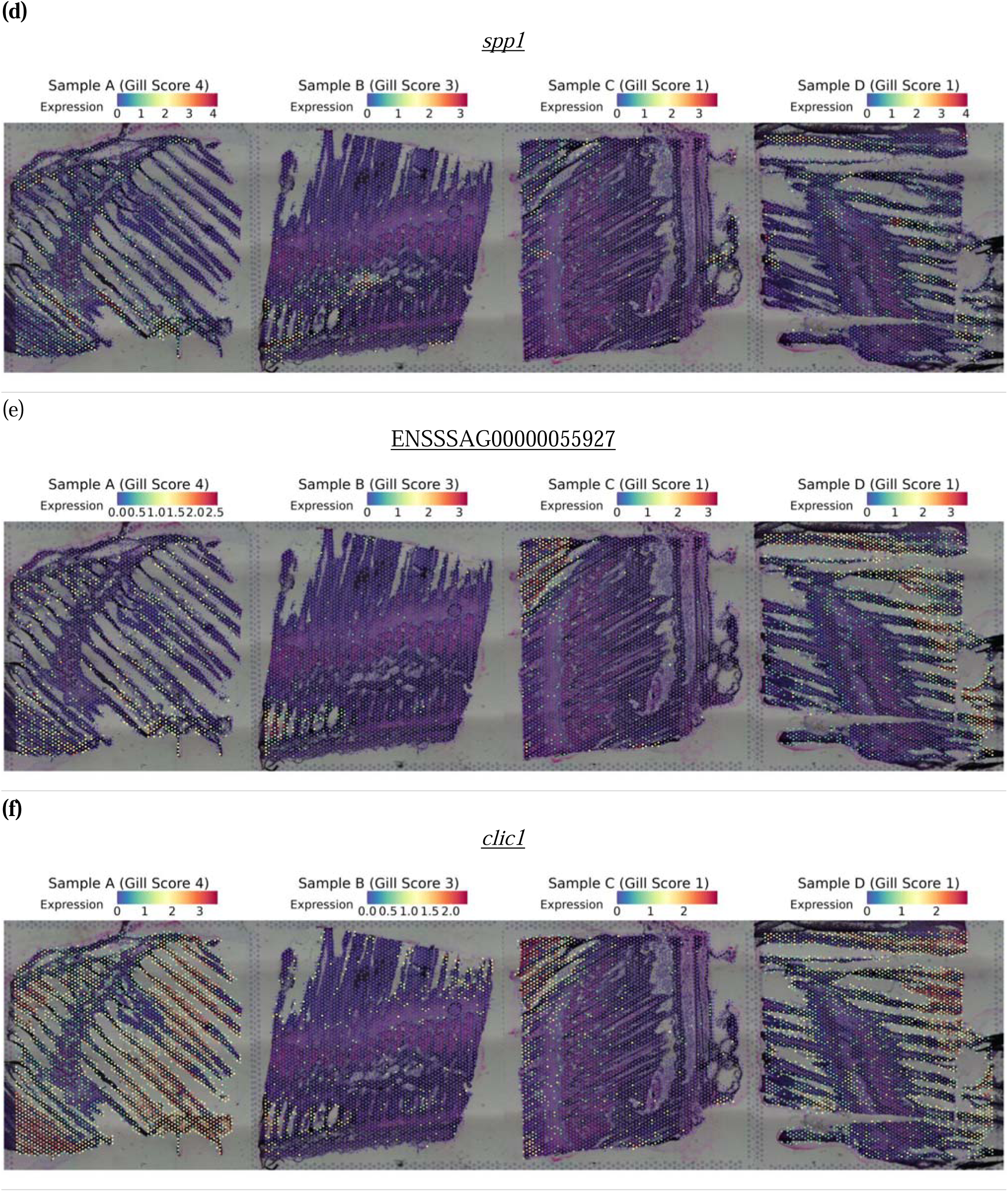
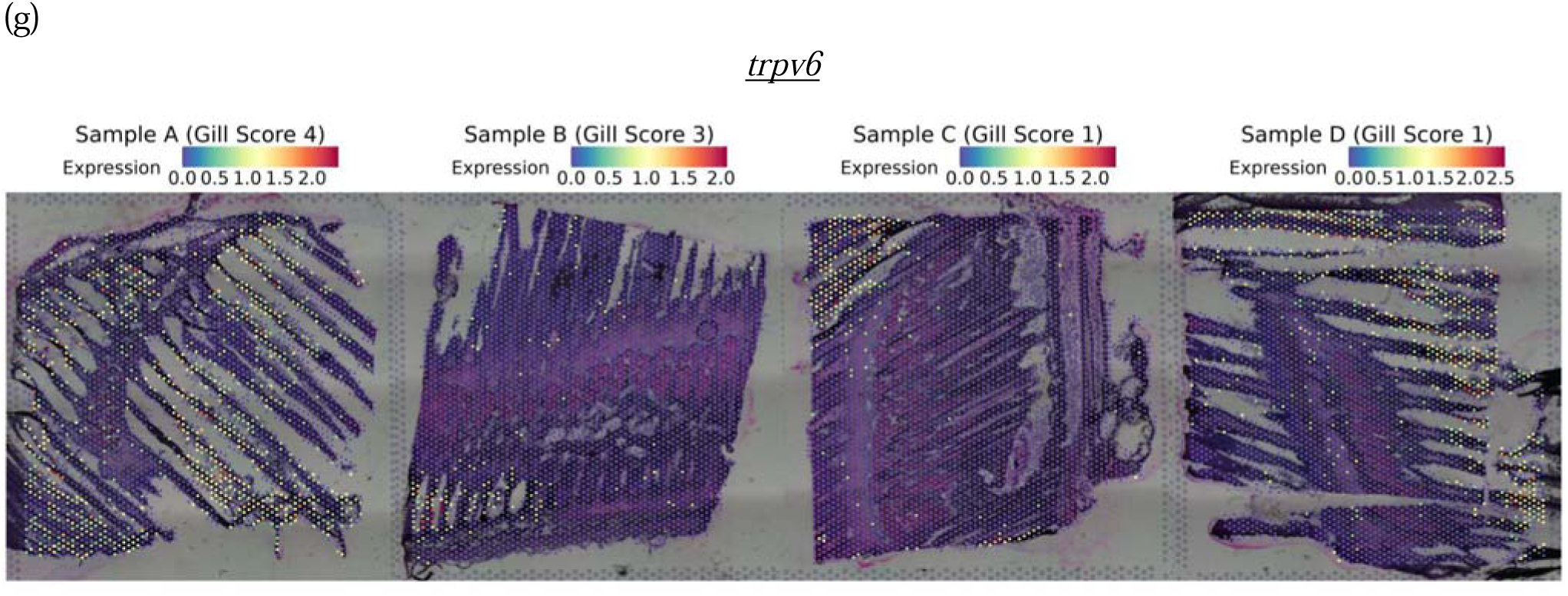
Spatial expression of representative marker genes for the filament and lamellae clusters, grouped by function. Epithelial integrity: (a) *krt15* (keratin 15; Ensembl ID: ENSSSAG00000045452; Cluster 10) and (b) ENSSSAG00000048372 (keratin, type I cytoskeletal 13; Entrez Gene ID: LOC106587847; UniProt: A0A1S3PWC1; Cluster 13). Extracellular matrix remodelling: (c) col10a1b (collagen, type X, alpha 1b; Ensembl ID: ENSSSAG00000114656; Cluster 11) and (d) spp1 (osteopontin; Ensembl ID: ENSSSAG00000100947; Cluster 11). Osmoregulation: (e) ENSSSAG00000055927 (sodium/hydrogen exchanger 3; Entrez Gene ID: LOC106569766; UniProt: A0A1S3M1V3; Cluster 12), (f) clic1 (chloride intracellular channel protein 1-like; Ensembl ID: ENSSSAG00000112534; Entrez Gene ID: LOC106588213; UniProt: A0A1S3PZ28; Cluster 2) and (g) trpv6 (transient receptor potential cation channel, subfamily V, member 6; Ensembl ID: ENSSSAG00000059186; Entrez Gene ID: LOC106592665; Cluster 12). Each panel corresponds to one tissue section from Samples A to D. The colour intensity reflects normalised expression, using SCTransform, where higher values indicate greater relative expression following sequencing depth correction.

#### 3.2.2 Gill arch clusters

Four clusters (4, 7, 8, and 9) located at the gill arch and adjacent smooth muscle regions across all samples (**Figure 8**). Their transcriptional profiles are consistent with functions expected in the gill arch, including connective tissue structure and extracellular matrix (ECM) remodelling, smooth muscle contractility, and vascular and lymphatic vessel activity. Connective tissue organisation and ECM remodelling are supported by structural and regulatory ECM components, such as *lum* (**Figure 9a**), *thbs4a* (**Figure 9b**), and *timp3* (**Figure 9c**). Smooth muscle contractility is reflected by high expression of cytoskeletal and contractile markers such as *cald1b* (**Figure 9d**), *acta2* (**Figure 9e**), *lmod1b* (**Figure 9f**), *smtna* (**Figure 9g**), *cnn1b* (**Figure 9h**), and *actn1* (**Figure 9i**). Vascular and lymphatic vessel activity is indicated by expression of *xlkd1* (**Figure 9j**), and *gp182* (**Figure 9k**). While these markers define the core functions of the gill arch and its adjacent smooth muscle regions, additional related genes identified in these clusters are detailed in the **Supplementary Note 9**.

**Figure 8.**
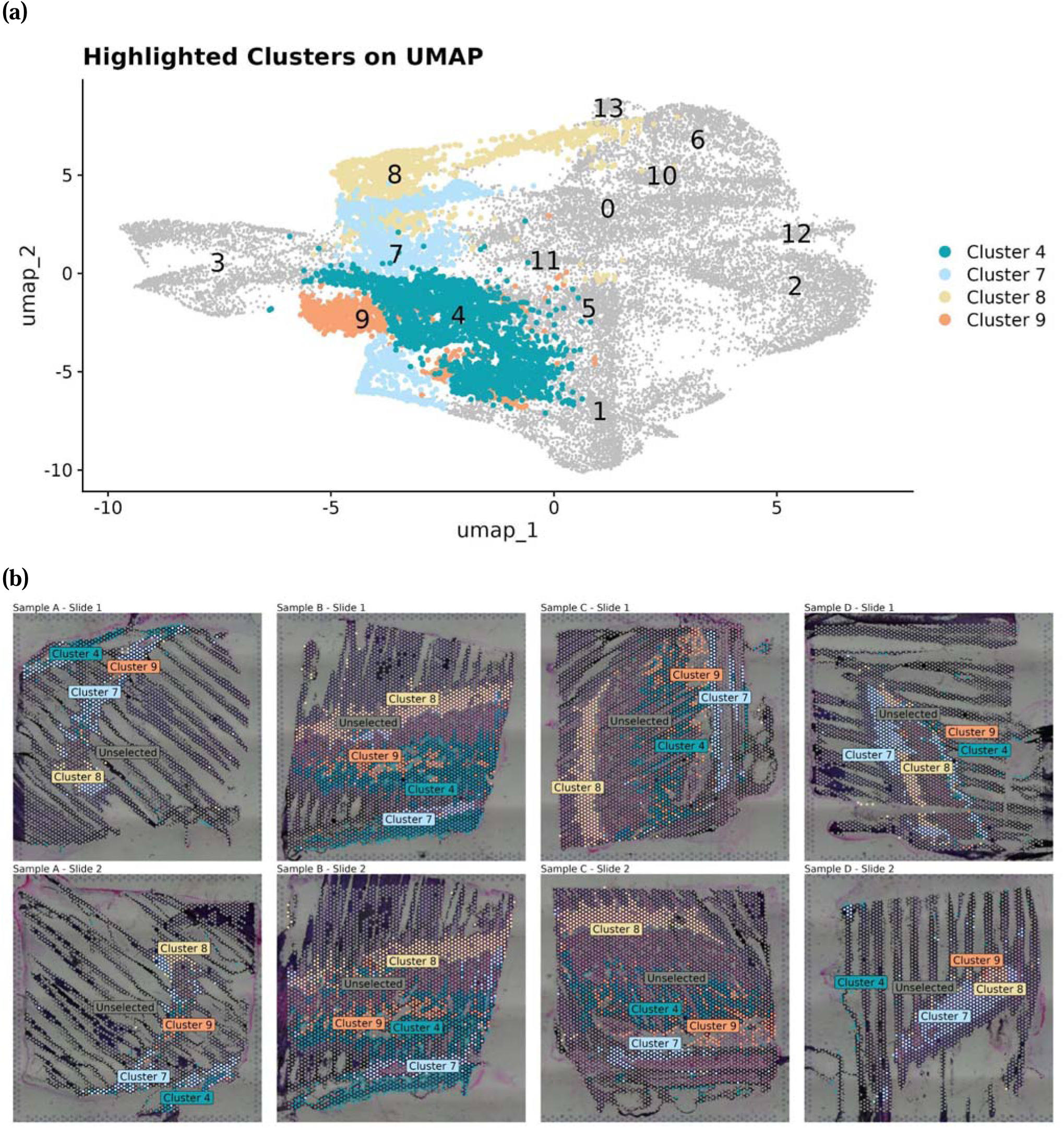
(a) UMAP projection subset highlighting four transcriptionally similar clusters, 4 (•), 7(•), 8(•), and 9(•), that collectively define gill arch clusters. Those spatial spots that are not part of the gill arch 2 are coloured as •. These clusters were identified using the Louvain algorithm at a resolution of 0.3, following PCA-based dimensionality reduction and construction of a Shared Nearest Neighbour (SNN) graph. Corresponding spatial mappings are shown in 8 panels in (b) with the corresponding sample name above each panel.

**Figure 9.**
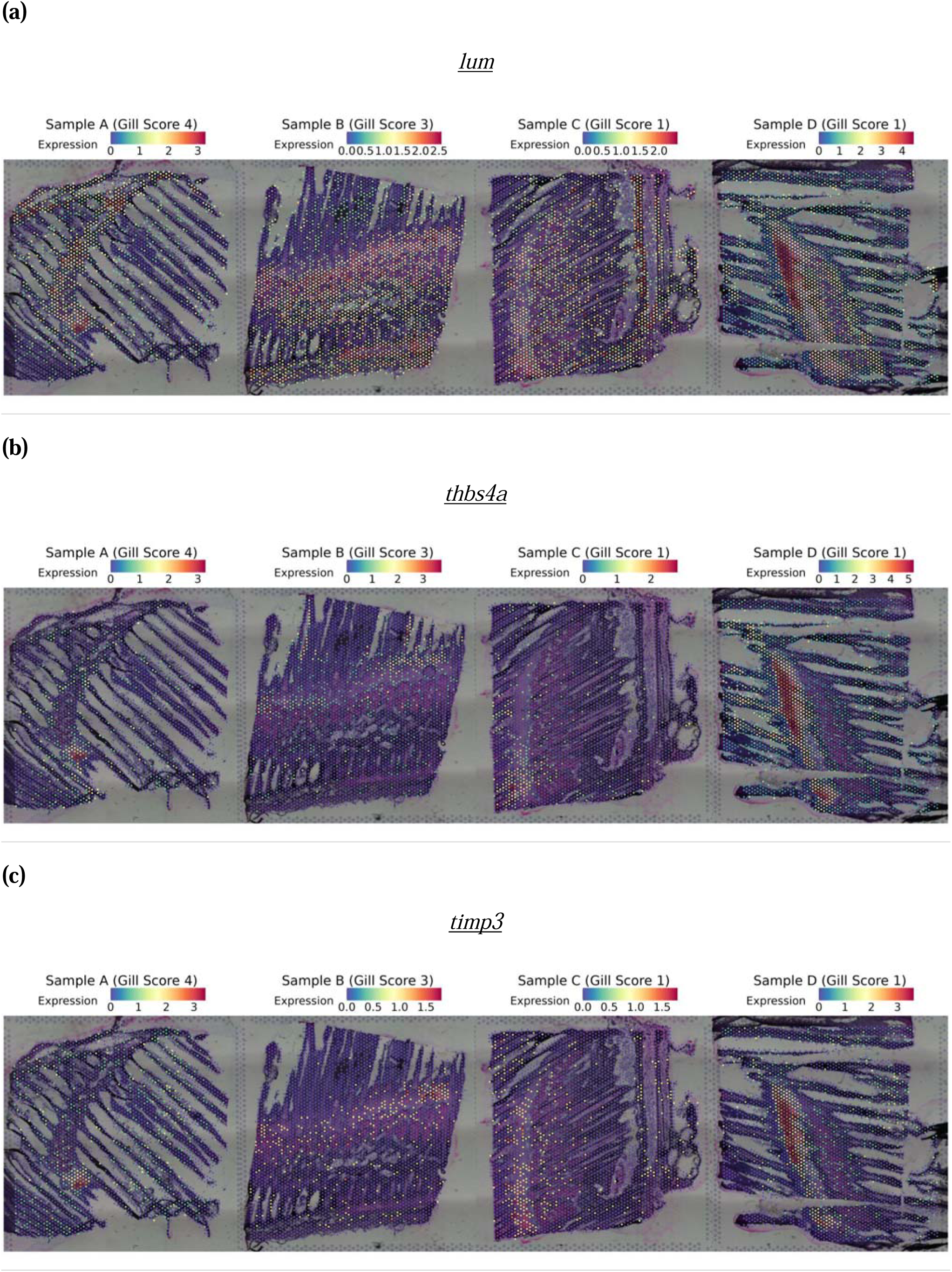
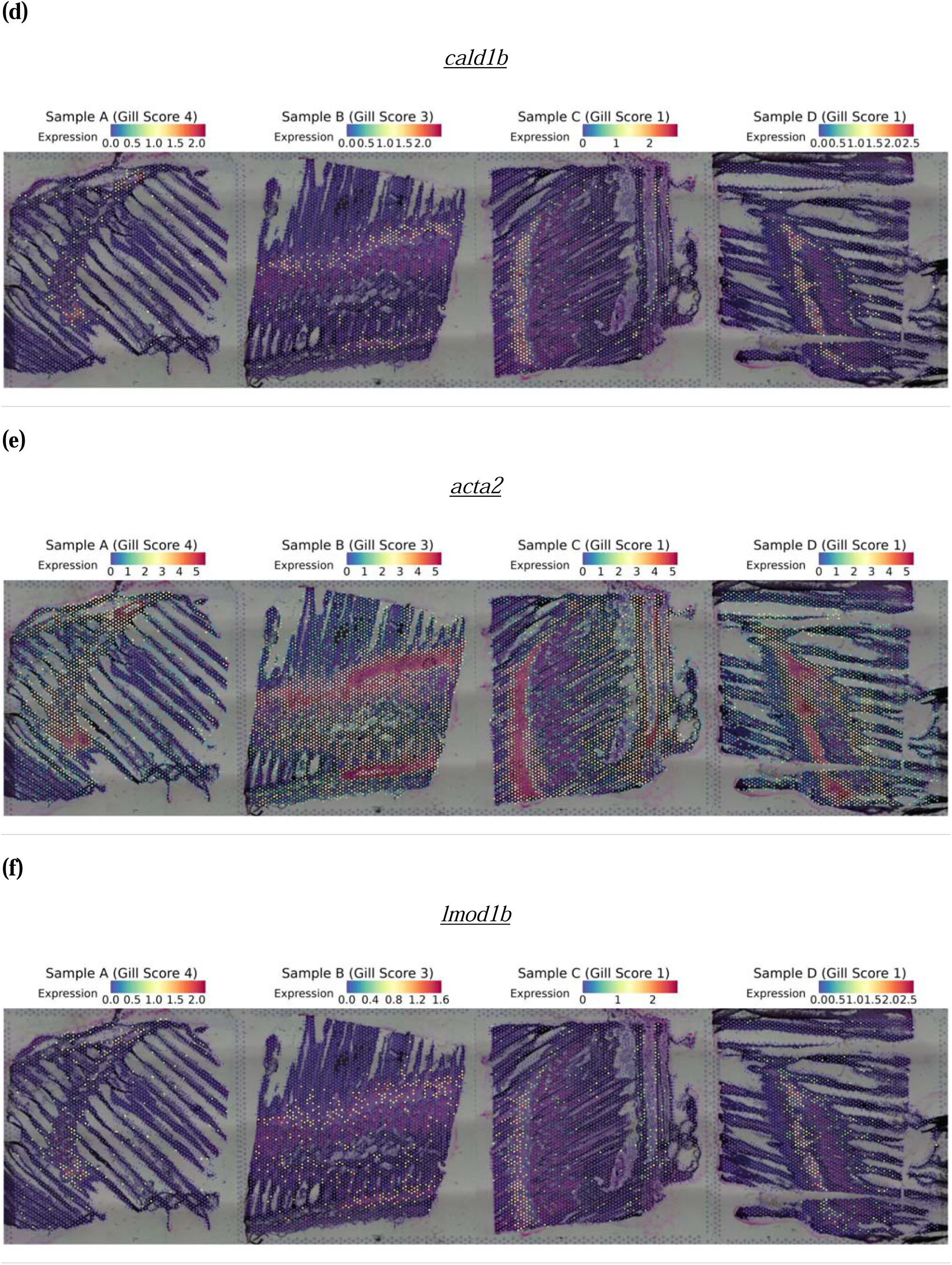
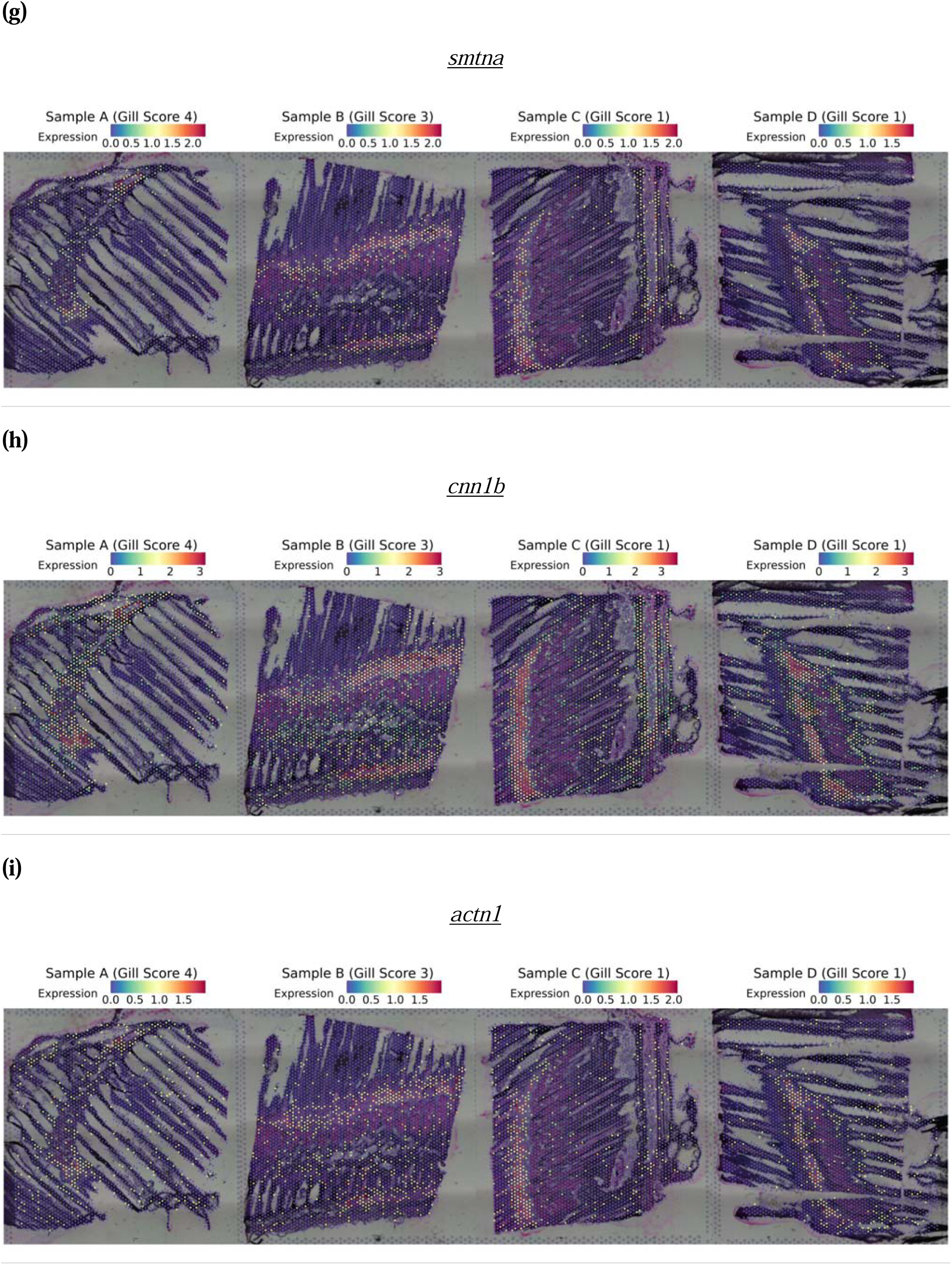
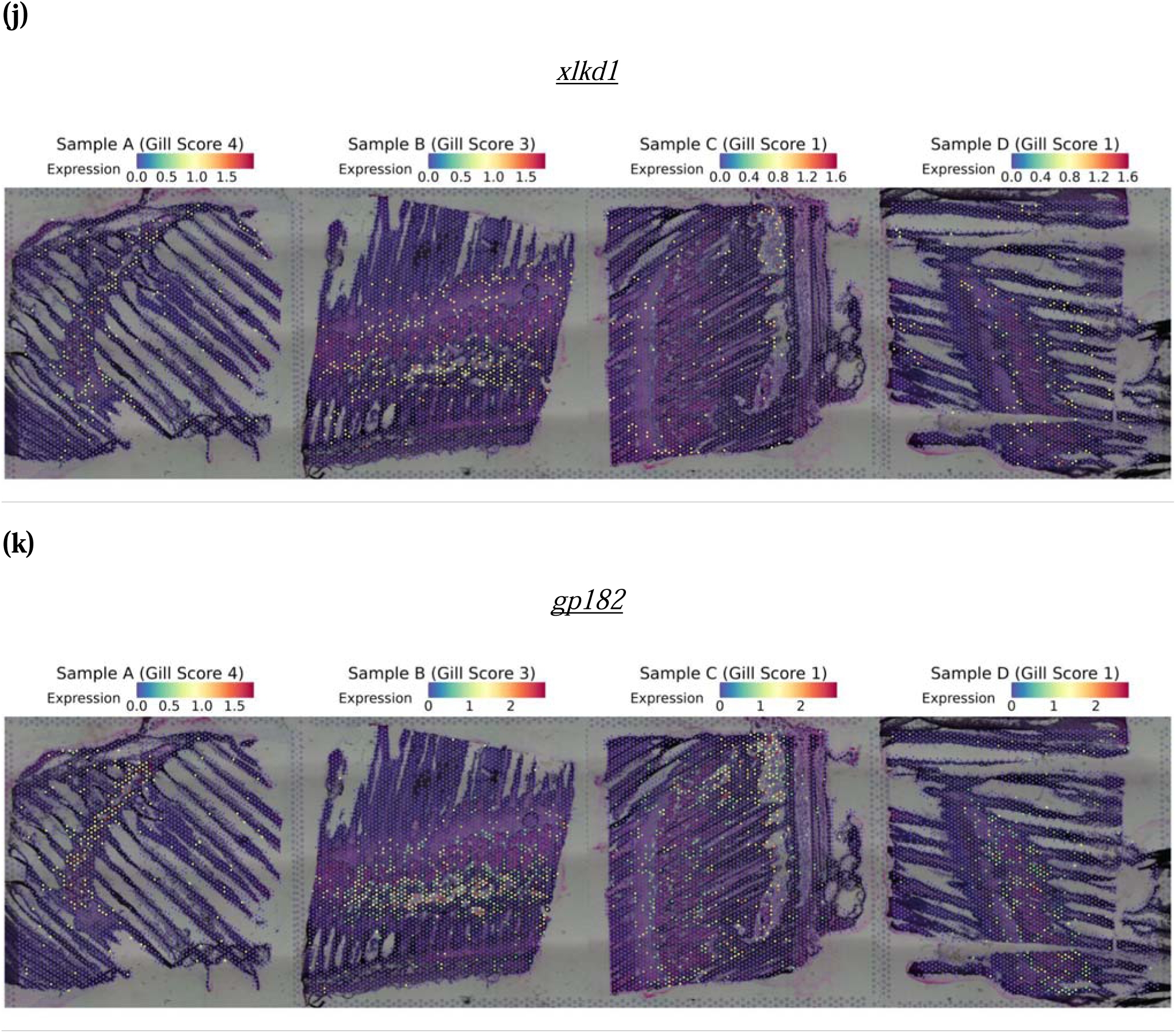
Spatial expression of representative marker genes for the gill arch clusters, grouped by function. Connective tissue and extracellular matrix remodelling: (a) *lum* (lumican; Ensembl ID: ENSSSAG00000020979; Cluster 7), (b) *thbs4a* (thrombospondin 4a; Ensembl ID: ENSSSAG00000071963; UniProt: A0A1S3PF42; Cluster 7) and (c) *timp3* (TIMP metallopeptidase inhibitor 3; Ensembl ID: ENSSSAG00000073256; Entrez Gene ID: LOC106609153; Cluster 7). Smooth muscle contractility: (d) *cald1b* (caldesmon 1b, non-muscle caldesmon-like isoform X1; Ensembl ID: ENSSSAG00000100045; Entrez Gene ID: LOC106573078; UniProt: A0A1S3MJG3; Cluster 8), (e) *acta2* (actin alpha 2, smooth muscle; Ensembl ID: ENSSSAG00000044179; UniProt: A0A1S3Q4A0; Cluster 8), (f) *lmod1b* (leiomodin 1b, smooth muscle; Ensembl ID: ENSSSAG00000064672; Entrez Gene ID: LOC106566054; UniProt: A0A1S3LEU6; Cluster 8), (g) *smtna* (smoothelin a; Ensembl ID: ENSSSAG00000050262; Entrez Gene ID: LOC106563188; UniProt: A0A1S3KZ62; Cluster 8), (h) *cnn1b* (calponin 1, basic, smooth muscle, b; Ensembl ID: ENSSSAG00000122078; Cluster 8) and (i) *actn1* (actinin, alpha 1; Ensembl ID: ENSSSAG00000114975; Cluster 8). Vascular and lymphatic vessel activity: (j) *xlkd1* (lymphatic vessel endothelial hyaluronic acid receptor 1; Ensembl ID: ENSSSAG00000072759; Cluster 9) and (k) *gp182* (G-protein coupled receptor 182; Ensembl ID: ENSSSAG00000088534). Each panel corresponds to one tissue section from Samples A to D. The colour intensity reflects normalised expression, using SCTransform, where higher values indicate greater relative expression following sequencing depth correction.

#### 3.2.3 Skeletal Muscle cluster

Marker genes in Cluster 3 highlight skeletal muscle components alongside signatures of high metabolic activity. Spatially, this cluster corresponds to muscle fibres within the gill (**Figure 10**). These fibers are primarily surrounded by areas of gill arch clusters that exhibit a smooth muscle transcriptional profile (**Figure 9d - i**). These transcriptional features reflect two main functional signatures: skeletal muscle contractile machinery and elevated metabolic activity supporting sustained muscle function. Skeletal muscle identity is supported by strong expression of contractile apparatus genes, including *tnnc2.2* (**Figure 11a**), *mlrb* (**Figure 11b**), and *actc1a* (**Figure 11c**), which collectively form the structural and regulatory machinery of striated muscle fibres. Elevated metabolic activity is indicated by strong expression of glycolytic and mitochondrial respiration genes, including *g3p* (**Figure 11d**), *cx6a1* (**Figure 11e**), and *cox4i2* (**Figure 11f**), supporting high ATP production required for muscle contraction. While these markers define the core functions of this cluster, additional related genes identified in this skeletal muscle cluster are detailed in the **Supplementary Note 10**.

**Figure 10.**
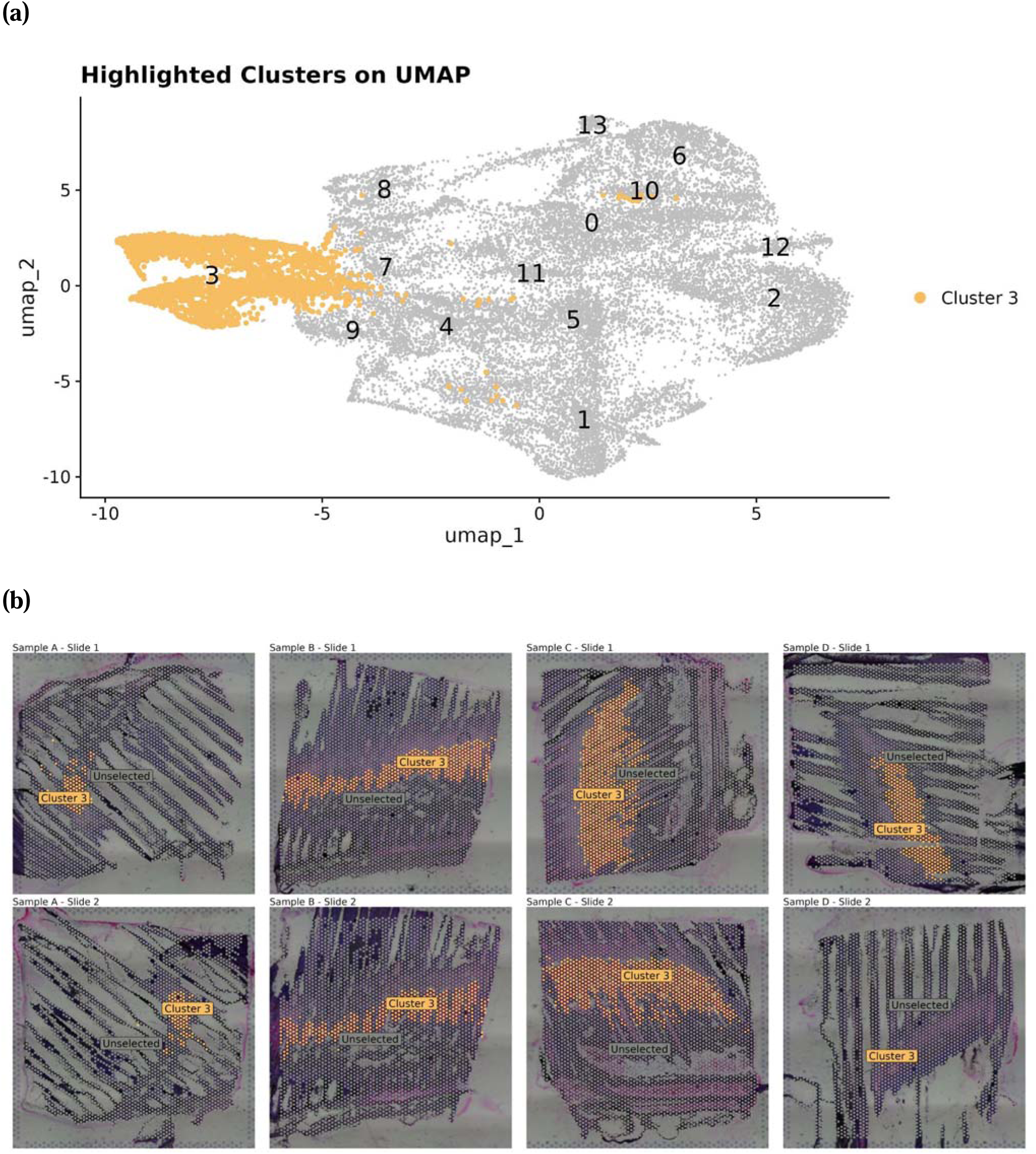
(a) UMAP projection subset highlighting a transcriptionally distinct cluster, Cluster 3 (•). Spatial spots that are not part of Cluster 3 are shown as (•). Clusters were identified using the Louvain algorithm at a resolution of 0.3, following PCA-based dimensionality reduction and construction of a Shared Nearest Neighbour (SNN) graph. Corresponding spatial mappings are shown in eight panels in (b), with the sample name indicated above each panel.

**Figure 11.**
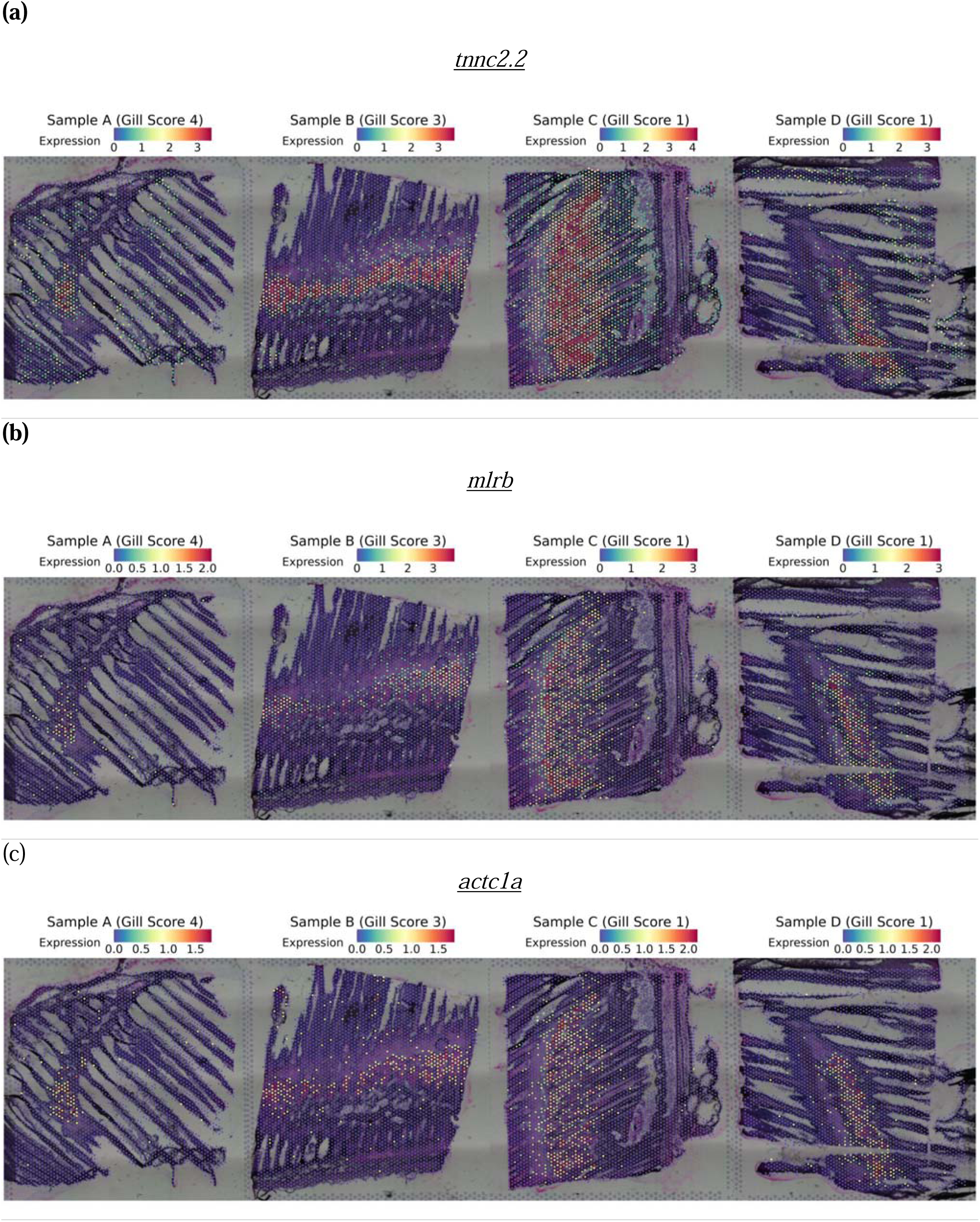
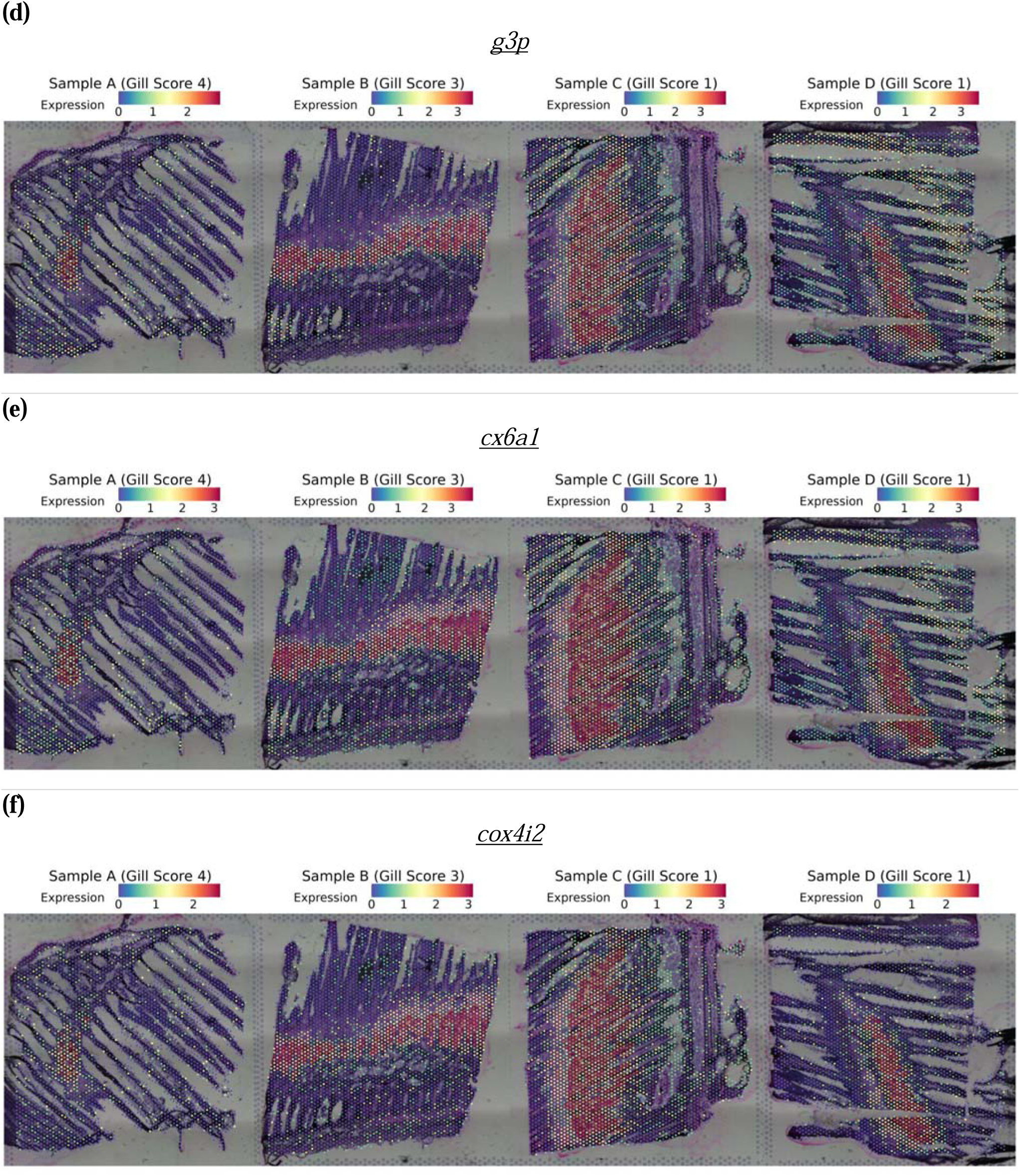
Spatial expression of representative marker genes for the skeletal muscle cluster (Cluster 3), grouped by function. Contractile apparatus: (a) *tnnc2.2* (troponin C2, fast skeletal type, tandem duplicate 2; Ensembl ID: ENSSSAG00000066537), (b) *mlrb* (myosin regulatory light chain 2B, cardiac muscle isoform; Ensembl ID: ENSSSAG00000062577) and (c) *actc1a* (actin alpha cardiac muscle 1a; Ensembl ID: ENSSSAG00000073229). Glycolysis: (d) *g3p* (glyceraldehyde-3-phosphate dehydrogenase; Ensembl ID: ENSSSAG00000073639). Oxidative phosphorylation: (e) *cx6a1* (cytochrome c oxidase subunit 6A, mitochondrial; Ensembl ID: ENSSSAG00000041369) and (f) *cox4i2* (cytochrome c oxidase subunit 4I2; Ensembl ID: ENSSSAG00000063792). Each panel corresponds to one tissue section from Samples A to D. The colour intensity reflects normalised expression, using SCTransform, where higher values indicate greater relative expression following sequencing depth correction.

#### 3.2.4 Haemoglobulin cluster

Top marker genes in Cluster 1 indicate transcriptional signatures associated with erythroid activity and vascular integrity. Spatially, this cluster is primarily localised within blood vessels of the gill arch and filaments, consistent with its association with the broader circulatory network (**Figure 12**). These transcriptional profiles support two main functional signatures: erythroid oxygen transport and platelet-associated vascular maintenance. Erythroid identity is reflected by strong expression of haemoglobin genes including *hbaa2* (**Figure 13a**), *hbae4* (**Figure 13b**), and *hbe1* (**Figure 13a**), forming a transcriptional signature consistent with erythroid cells. Platelet-associated vascular functions are indicated by expression of *gp1bb* (**Figure 13d**), which contributes to platelet-mediated vascular integrity, and *septin3* (**Figure 13e**), a septin cytoskeletal protein associated with platelet structure and function. While these markers define the core functions of this cluster, additional related genes identified in this Haemoglobulin related cluster are detailed in the **Supplementary Note 11**.

**Figure 12.**
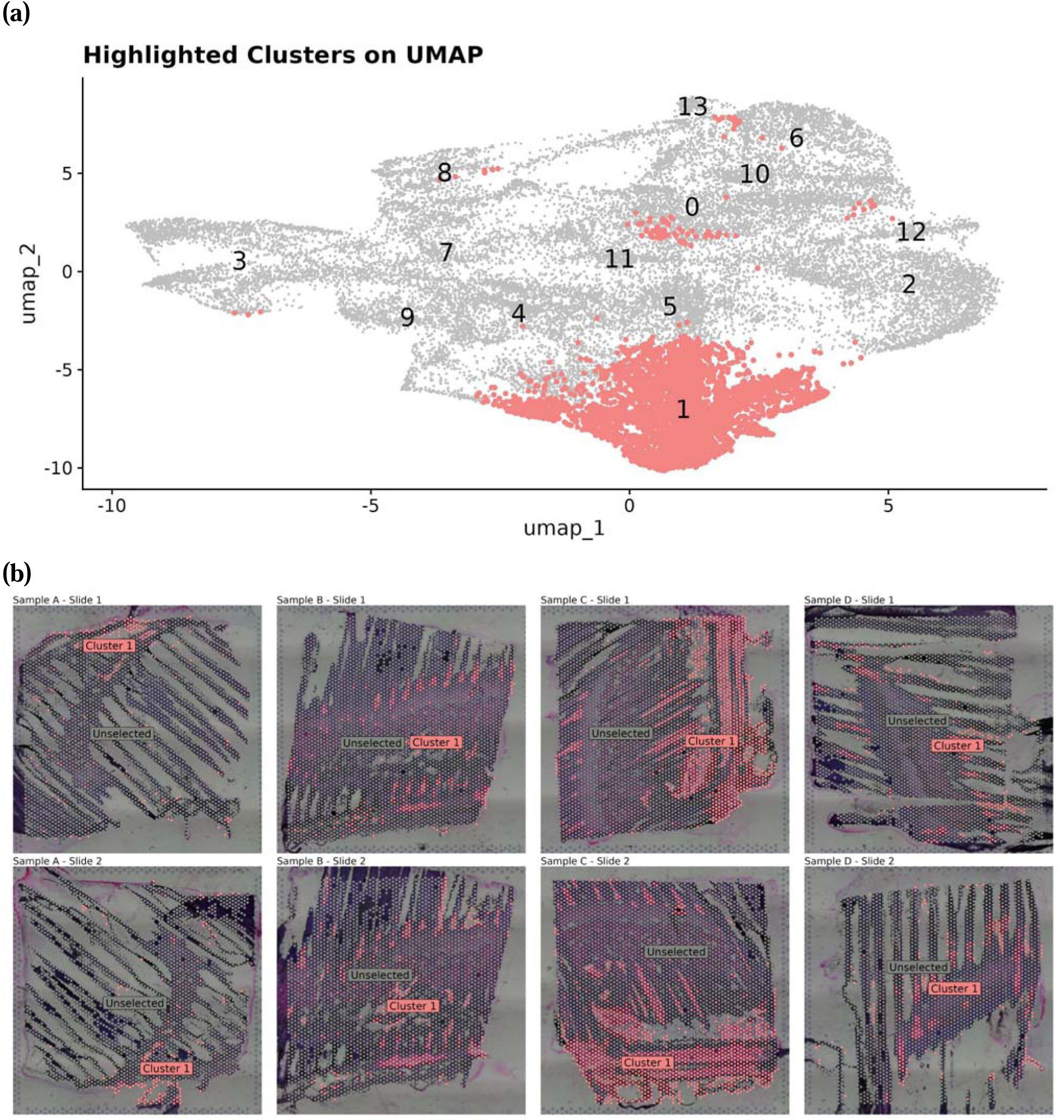
(a) UMAP projection subset highlighting a transcriptionally distinct cluster, Cluster 1 (•). Spatial spots not part of Cluster 1 are shown as (•). Clusters were identified using the Louvain algorithm at a resolution of 0.3, following PCA-based dimensionality reduction and construction of a Shared Nearest Neighbour (SNN) graph. Corresponding spatial mappings are shown in eight panels in (b), with the sample name indicated above each panel.

**Figure 13.**
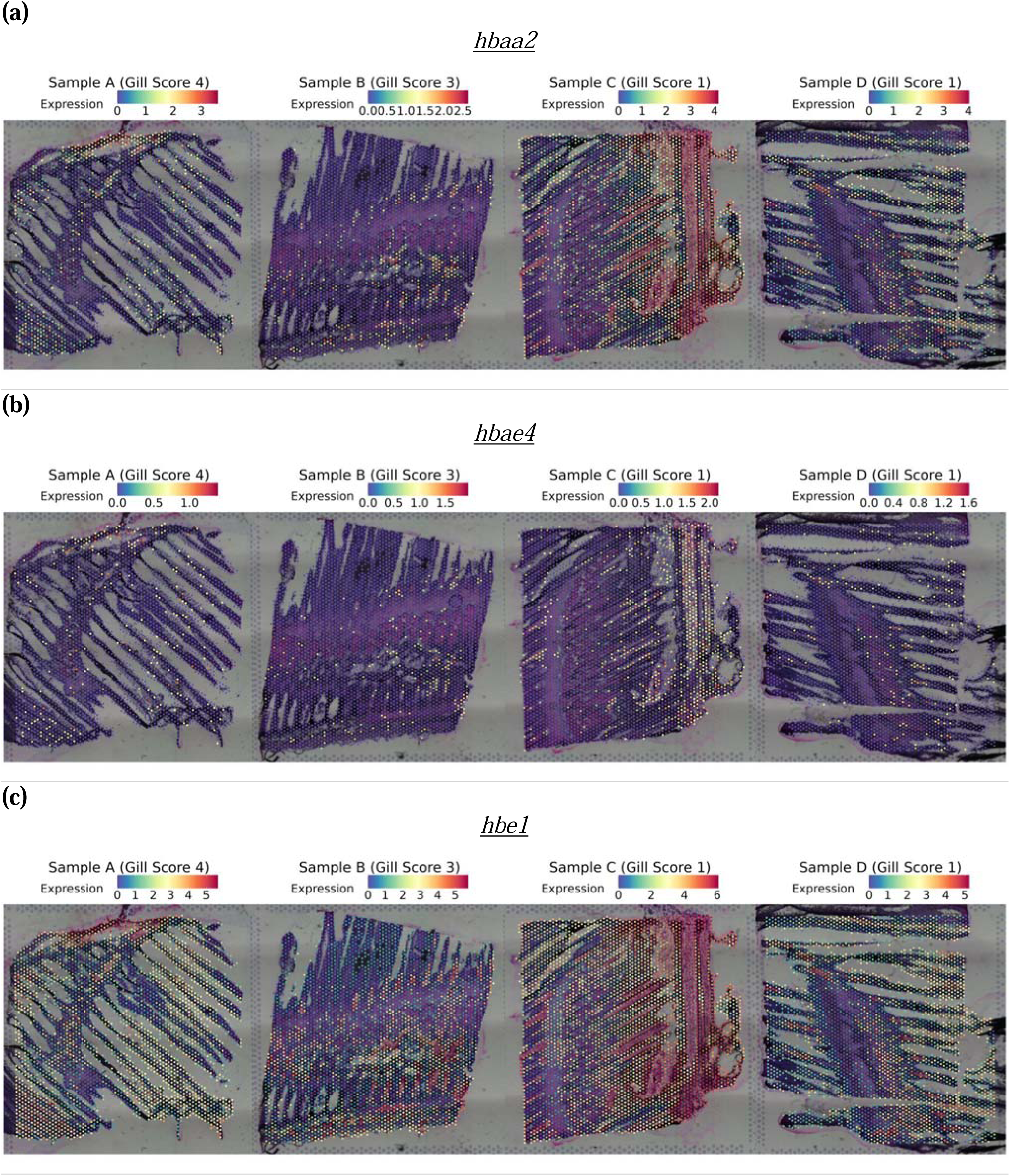
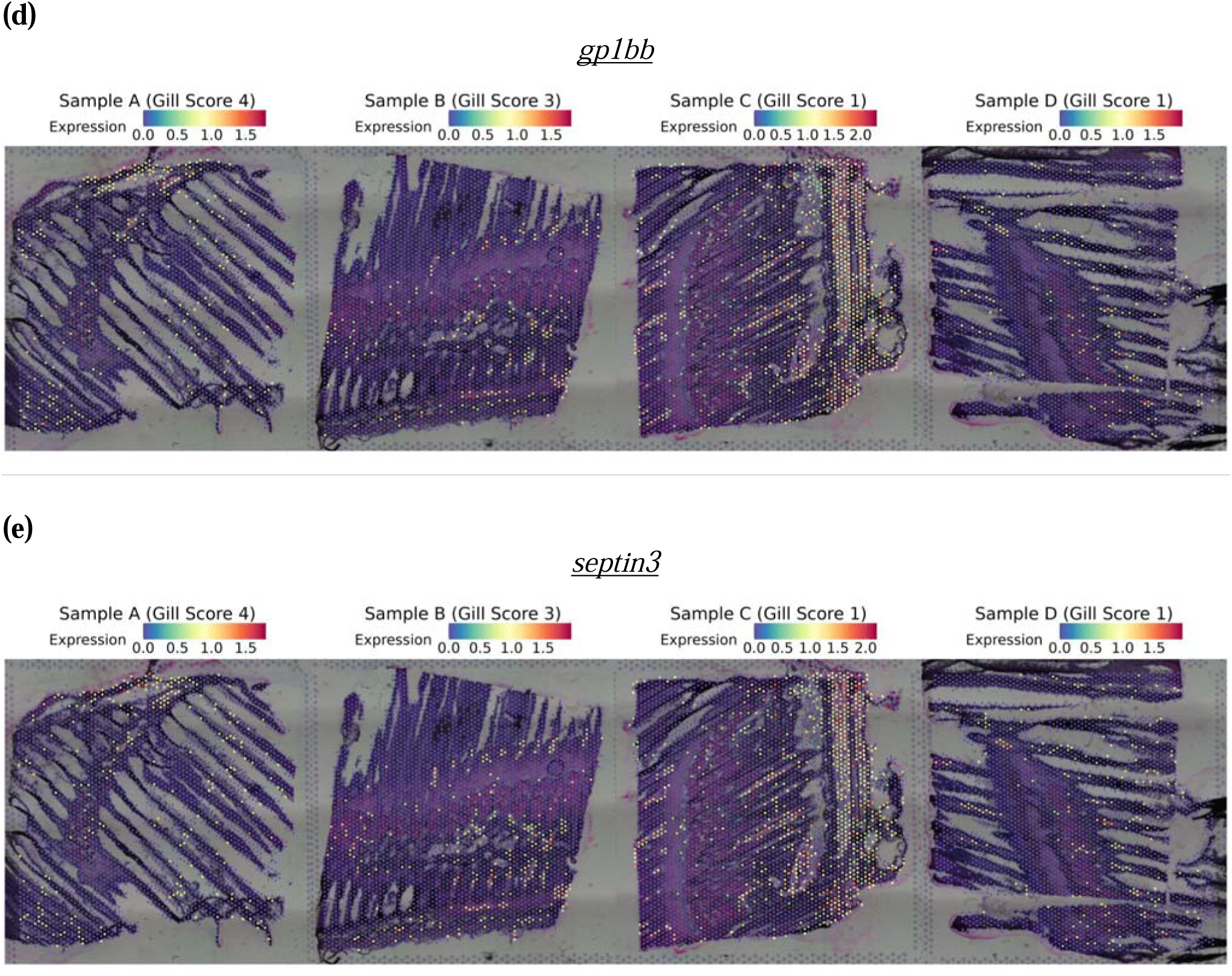
Spatial expression of representative marker genes for the haemoglobin cluster (Cluster 1), grouped by function. Erythroid oxygen transport: (a) *hbaa2* (haemoglobin, alpha adult 2; Ensembl ID: ENSSSAG00000065229), (b) *hbae4* (haemoglobin alpha embryonic-4; Ensembl ID: ENSSSAG00000046009) and (c) *hbe1* (haemoglobin embryonic-1; Ensembl ID: ENSSSAG00000065233). Thrombocyte-associated vascular maintenance: (d) *gp1bb* (glycoprotein Ib platelet subunit beta; Ensembl ID: ENSSSAG00000084006) and (e) *septin3* (septin 3; Ensembl ID: ENSSSAG00000007203). Each panel corresponds to one tissue section from Samples A to D. The colour intensity reflects normalised expression, using SCTransform, where higher values indicate greater relative expression following sequencing depth correction.

### 3.3 Immunological Functions spanning across the four main anatomical domains of the gill

#### 3.3.1 Spatial Distribution of Innate Immune Effectors and Pattern Recognition Receptors in the Gill

To recognise pathogen-associated molecular patterns (PAMPs) and initiate immune responses, several classes of pattern recognition receptors (PRRs) are involved. Elevated PRR expression was observed among the top marker genes across multiple clusters.

Among the pattern recognition receptors (PRRs) showing elevated expression, C-type lectins receptors (CLRs), a PRR subgroup, were enriched. CLRs are primarily found on the surface of innate immune cells such as macrophages and dendritic cells (Takeuchi and Akira, 2010). They are characterised by C-type lectin domains that bind carbohydrate structures (glycans) found on pathogens, such as fungal mannans, β-glucans, and bacterial components (Brown *et al*., 2018). Across multiple spatial clusters in the gill, three genes without official symbols but containing C-type lectin-like domains were significantly upregulated, each showing distinct spatial distributions. These included: ENSSSAG00000055826 (**Figure 14a**), which was highly expressed in both the gill arch and filaments; ENSSSAG00000003409 (**Figure 14b**), which was associated with the gill arch, showing expression linked to vascular and smooth muscle regions and differentially expressed in a domain likely involved in vascular and lymphatic functions; and ENSSSAG00000116244 (**Figure 14c**), which was scattered throughout the filaments but absent from the gill arch. Furthermore, another member of the C-type CLR family, ***mrc1*** (**Figure 14d**), was highly expressed in Cluster 9, corresponding to regions of the gill arch and filaments. These C-type lectin domain expressions highlight potential ongoing pathogen sensing and uptake by means of CLRs within Cluster 4 and 9 (filament and lamellae clusters), as well as Cluster 10 and 11 (gill arch clusters). The spatial proximity between Cluster 4 and Cluster 9 suggests coordinated immune activity, possibly involving lymphoid tissue, while the location of Clusters 10 and 11 along the gill filament may reflect roles in local immune surveillance.

**Figure 14.**
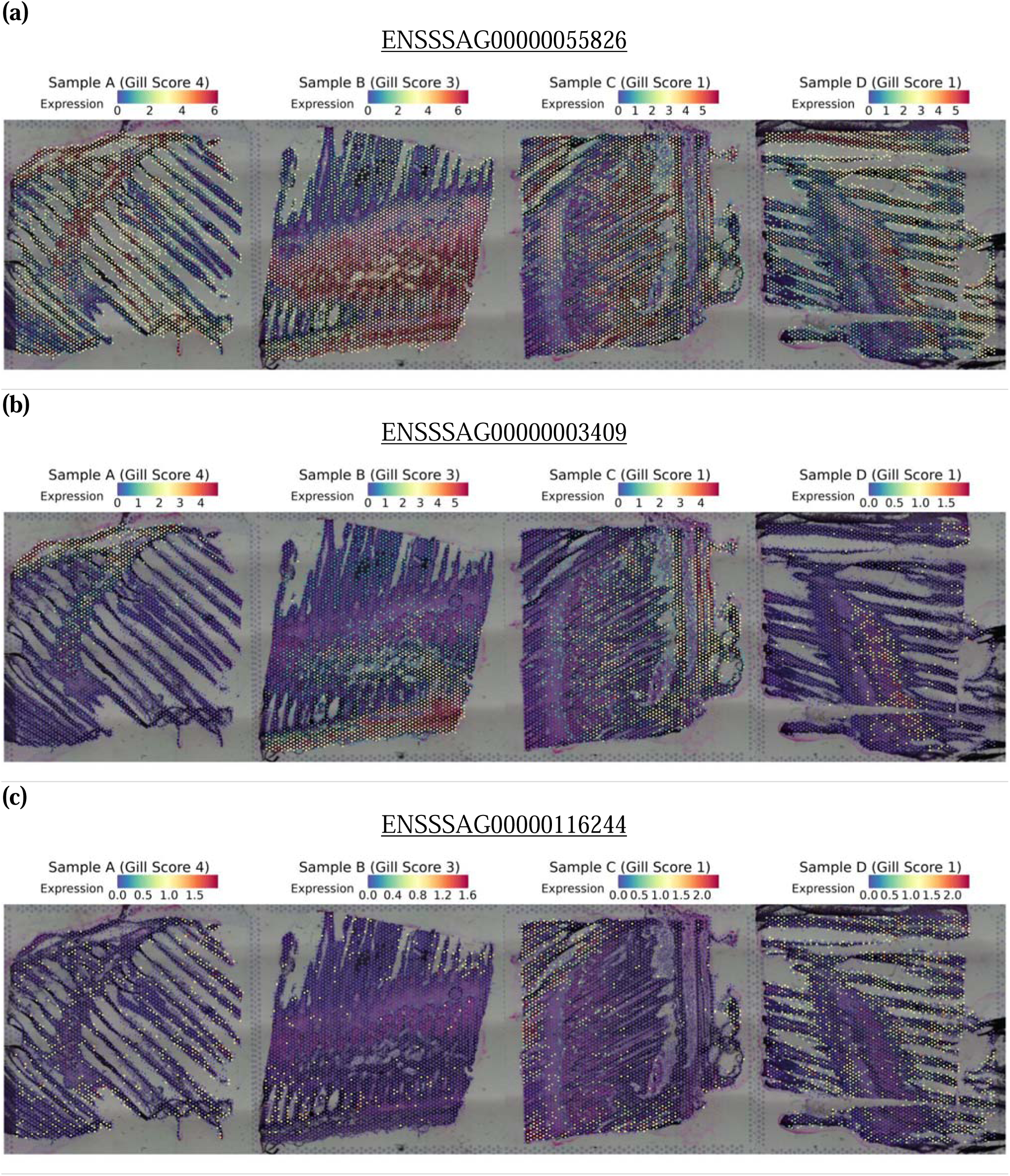
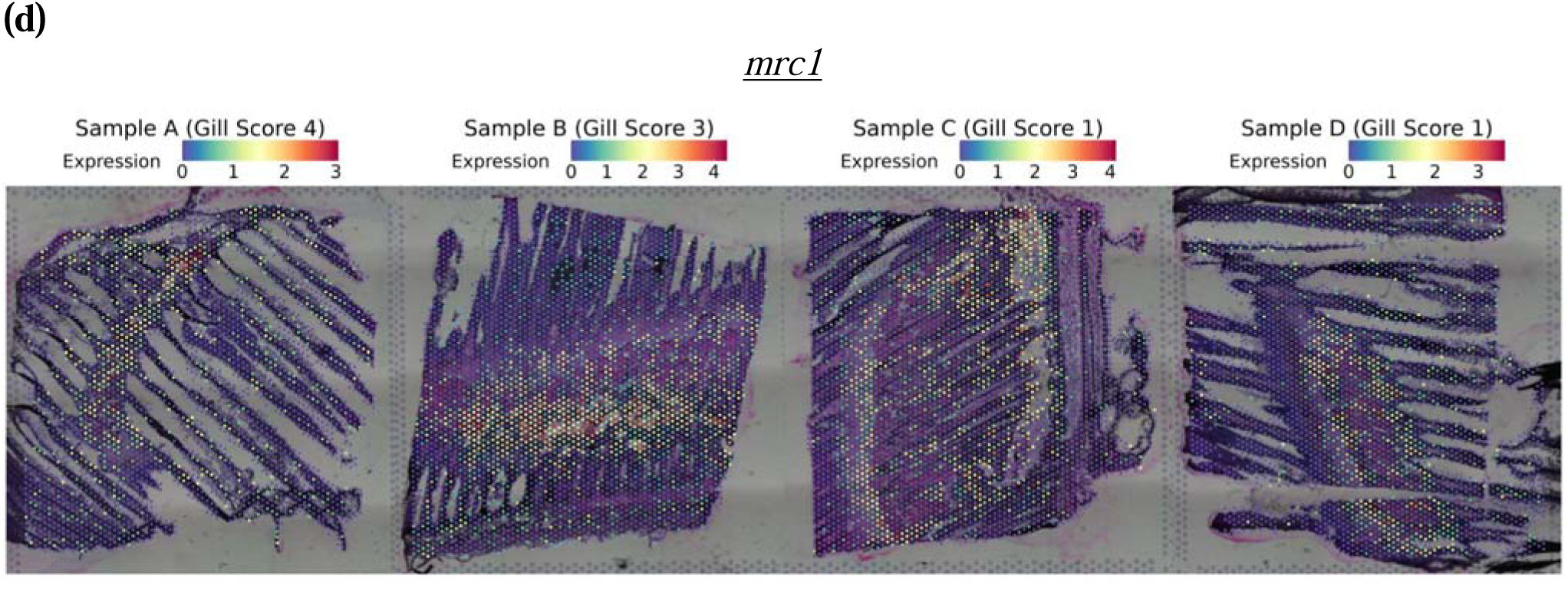
Spatial expression of genes encoding C-type lectin domain-containing proteins: (a) ENSSSAG00000055826 (C-type lectin-like domain-containing; Pfam: PF00059; InterPro: IPR001304; Clusters 4 and 11), (b) ENSSSAG00000003409 (C-type lectin-like domain-containing; Pfam: PF00059; InterPro: IPR001304; Cluster 4), (c) ENSSSAG00000116244 (C-type lectin-like domain-containing; Pfam: PF00059; InterPro: IPR001304; Cluster X) and (d) *mrc1* (mannose receptor C-type 1; Ensembl ID: ENSSSAG00000073565; Cluster 9). Each panel corresponds to one tissue section from Samples A to D. The colour intensity reflects normalised expression, using SCTransform, where higher values indicate greater relative expression following sequencing depth correction.

Rhamnose-binding lectins (RBLs), members of the lectin family, have been identified as PRRs in invertebrates and teleost fish (Bovo and Ballarin, 2023; Zhang et al., 2025). These RBLs are known for their ability to bind bacterial surface carbohydrates, thereby facilitating immune recognition. The RBL *sal* (**Figure 15**) was differentially expressed in Cluster 0 and showed localisation along the gill filament but was absent from the gill arch. This spatial pattern suggests that *sal* may contribute to localised immune surveillance in the filament, potentially through the recognition of bacterial components and the initiation of inflammatory responses or phagocytosis.

**Figure 15:**
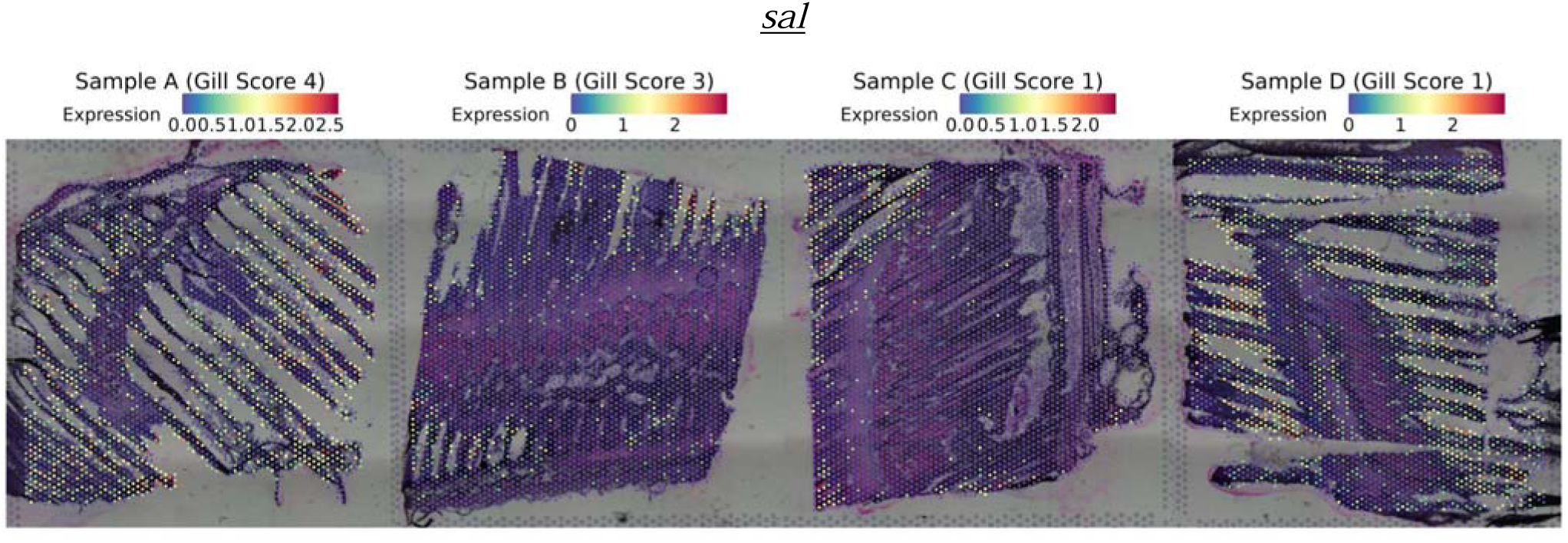
Spatial expression of *sal* (rhamnose-binding lectin; Ensembl ID: ENSSSAG00000005097; Cluster 0). Each panel corresponds to one tissue section from Samples A to D. The colour intensity reflects normalised expression, using SCTransform, where higher values indicate greater relative expression following sequencing depth correction.

Antimicrobial peptides and enzymes are key components of the innate immune response in fish, contributing to first-line defence against aquatic pathogen (Magnadóttir, 2006). C-type lysozymes are among these effectors, functioning as enzymatic molecules that degrade bacterial cell walls particularly effective against Gram-positive bacteria (Saurabh and Sahoo, 2008). Widely conserved across fish species, C-type lysozymes have been studied for both their expression profiles and antimicrobial properties (Chen *et al*., 2022; Kim *et al*., 2023). In the spatial transcriptomics, *ENSSSAG00000001998* (**Figure 16a**), a lysozyme-like domain containing gene, was enriched in Cluster 2, suggesting localised lysozyme activity within the secondary gill lamellae and potential involvement in bacterial cell wall degradation. Similarly, cathelicidin antimicrobial peptides play a critical role in mucosal defence (Chang *et al*., 2006; D’Este *et al*., 2016). In rainbow trout, cathelicidins are synthesised as precursors containing a conserved cathelin-like domain, from which the mature Gly/Ser-rich peptides are released; these peptides display broad-spectrum, salt-sensitive antimicrobial activity linked to structural features such as a positively charged N-terminus and central repetitive motifs (Chang *et al*., 2005, 2006; D’Este *et al*., 2016). One of the top marker genes in Clusters 6 and 13, *cath1a* (**Figure 16b**), was identified as corresponding to regions of the gill filament and lamellae, indicating active peptide-mediated defence at these sites. Collectively, the expression of lysozyme- and cathelicidin-related genes across distinct spatial clusters supports the presence of layered innate immune mechanisms within the gill epithelium.

**Figure 16.**
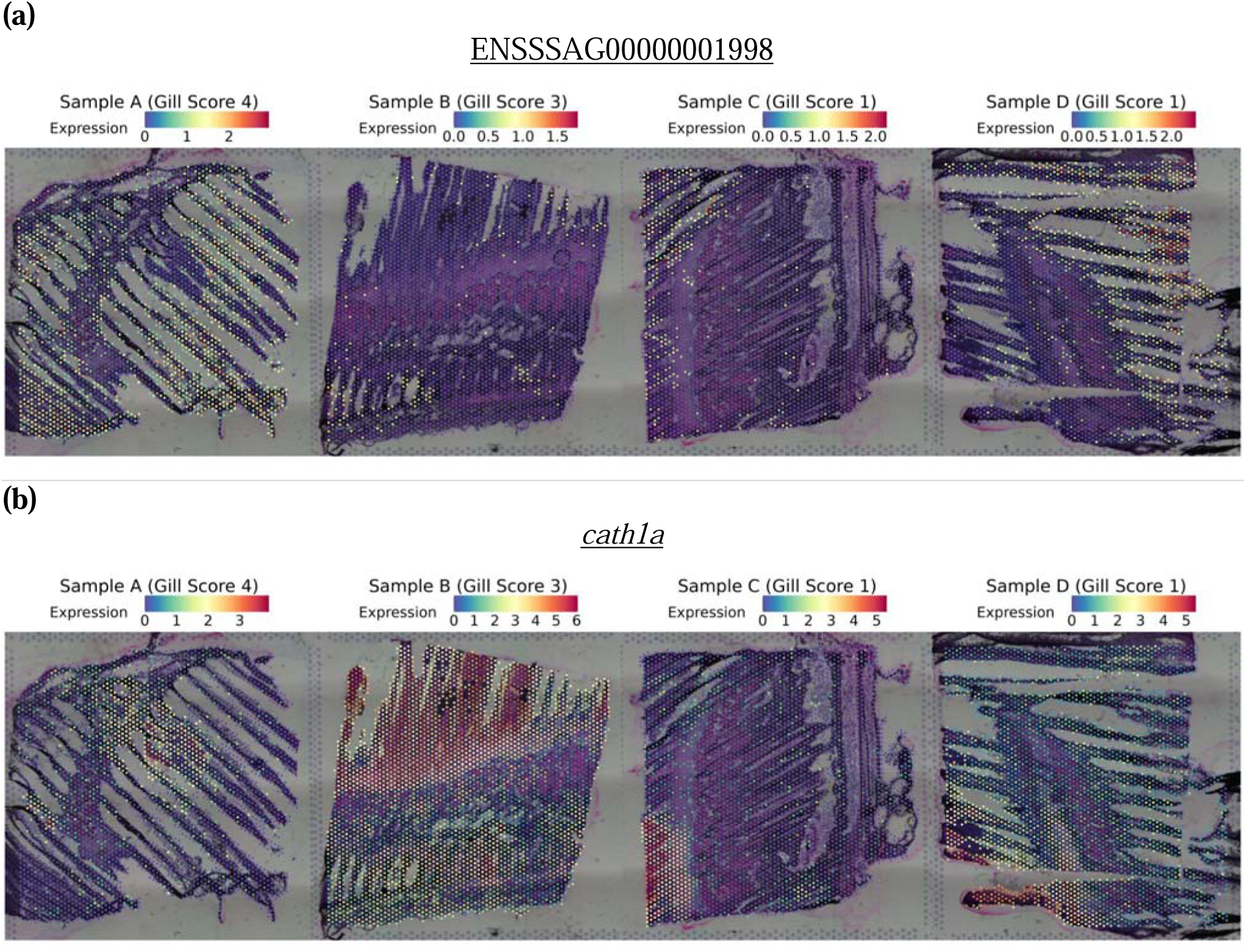
Spatial expression of genes encoding a lysozyme-like domain-containing protein and a cathelicidin: (a) ENSSSAG00000001998 (lysozyme-like domain-containing; Pfam: PF00062; InterPro: IPR001916; Cluster 2) and (b) *cath1a* (cathelicidin antimicrobial peptide; Ensembl ID: ENSSSAG00000074950; Clusters 6 and 13). Each panel corresponds to one tissue section from Samples A to D. The colour intensity reflects normalised expression, using SCTransform, where higher values indicate greater relative expression following sequencing depth correction.

In addition to enzymatic and peptide-mediated defences, cellular innate immune effectors also showed distinct spatial distributions within the gill. A signature marker of neutrophils, *mpx* (**Figure 17**), which is involved in generating potent oxidants for antimicrobial activity, was one of the top marker genes in Cluster 11. The spatial expression pattern of *mpx* indicated its localisation within both the gill lamellae and filaments. Such elevated expression suggests a high abundance of neutrophils in these regions, reflecting active cellular innate immune responses and highlighting the oxidative burst as a key mechanism for pathogen clearance at this mucosal interface.

**Figure 17.**
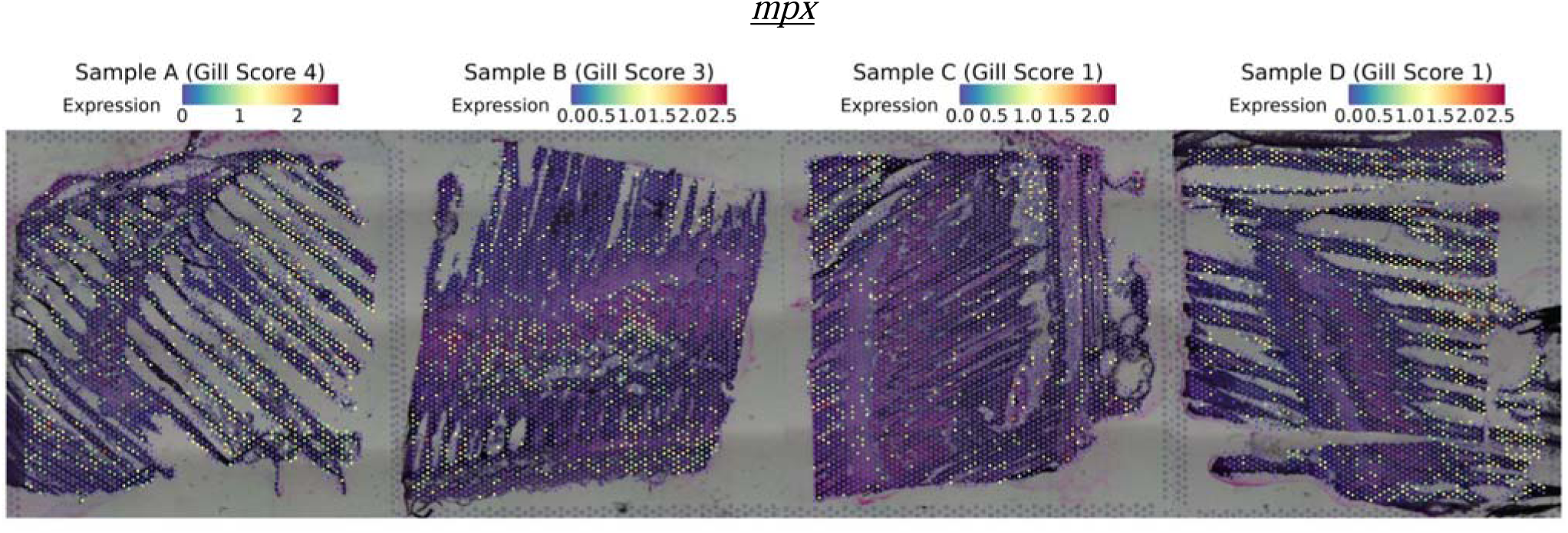
Spatial expression of *mpx* (myeloid-specific peroxidase; Ensembl ID: ENSSSAG00000000157). Each panel corresponds to one tissue section from Samples A to D. The colour intensity reflects normalised expression, using SCTransform, where higher values indicate greater relative expression following sequencing depth correction.

#### 3.3.2 Chemotactic Signalling and Immune Cell Recruitment in the Gill

The recruitment of immune cells to sites of challenge is strongly indicated by the presence of multiple chemokines as top marker genes across distinct spatial clusters in the gill. C-C motif chemokine 28 (*ccl28*), known in mammals for its role in immune cells recruitment and antimicrobial defence, is highly expressed in mucosal tissues such as the skin and gill, suggesting a conserved function in mucosal immunity (Hieshima *et al*., 2003; Muñoz-Atienza *et al*., 2019). In the spatial transcriptomics dataset, *ccl28* (**Figure 18a**) was among the top markers in Cluster 0, located within the gill filament, indicating a potential role in maintaining frontline immune surveillance. Similarly, ENSSSAG00000033609 (**Figure 18b**), a top marker gene in Cluster 10, is known in teleosts to mediate the recruitment of lymphocytes and macrophages during infection, indicating ongoing immune cell trafficking within the gill filament (Bjørgen *et al*., 2019; Sepahi *et al*., 2017; Wei *et al*., 2021). In Cluster 13, *cxl10* (**Figure 18c**) was highly expressed; its functional homolog in the Japanese flounder (PoCXCL10) is known to regulate leukocyte migration, cytokine production, and phagocytosis, in addition to exhibiting direct antimicrobial activity through its membrane-disruptive C-terminal α-helix (Li et al., 2022). Furthermore, ENSSSAG00000045448 (**Figure 18d**) is also among the top markers in Cluster 13. Interleukin-1 is a key pro-inflammatory cytokine in salmon involved in orchestrating immune responses via the IL-1 receptor pathway (Morrison *et al*., 2012). Its expression at the gill filament and lamellae further indicates that this region serves as a site of inflammatory signalling and cellular activation. Therefore, the presence of ccl28, ccl19-like, cxl10, and interleukin-1 as top markers across spatial clusters highlights the capacity of the gill for localised chemotactic signalling, supporting immune cell recruitment and coordination of early immune responses at mucosal surfaces.

**Figure 18.**
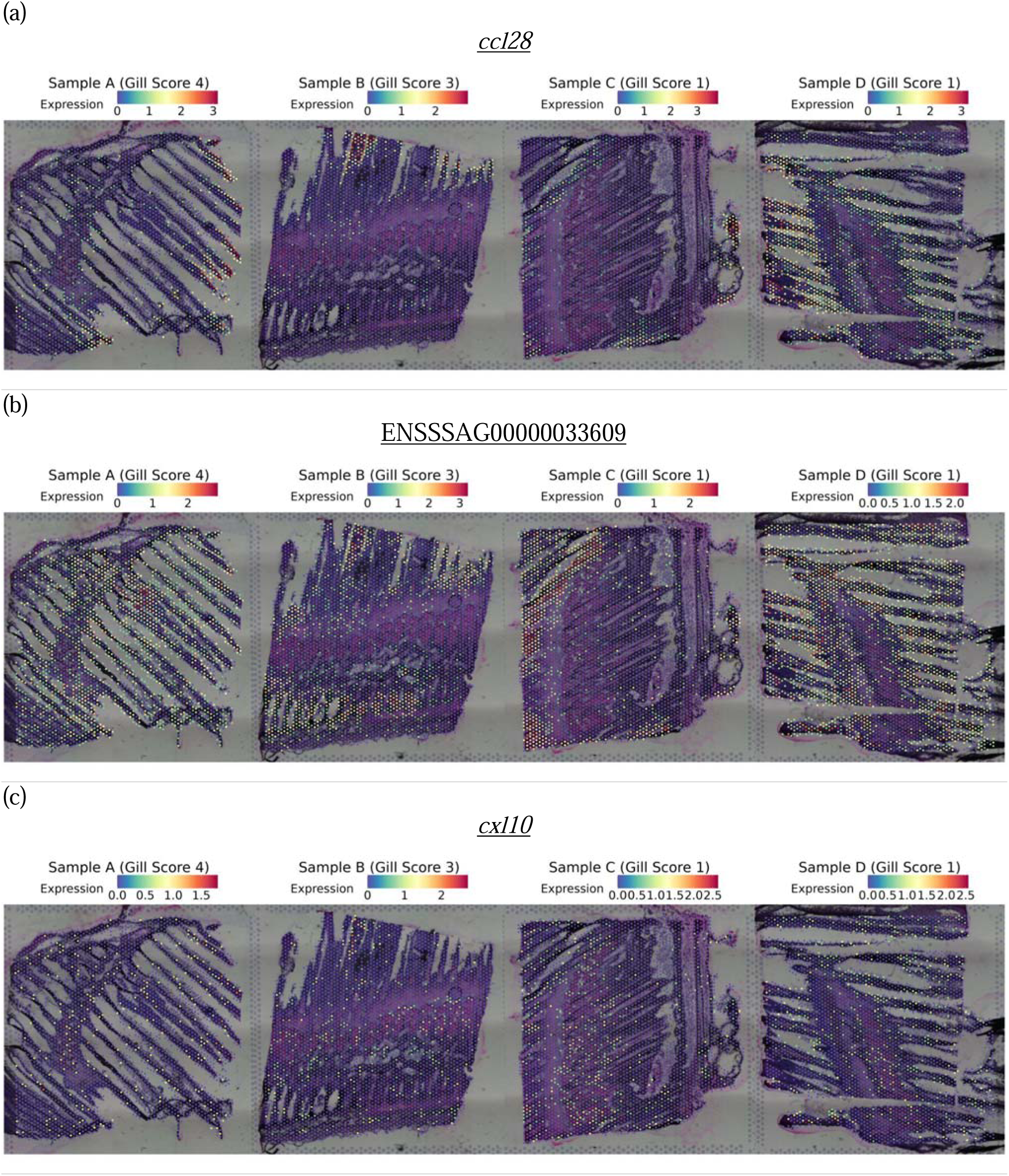
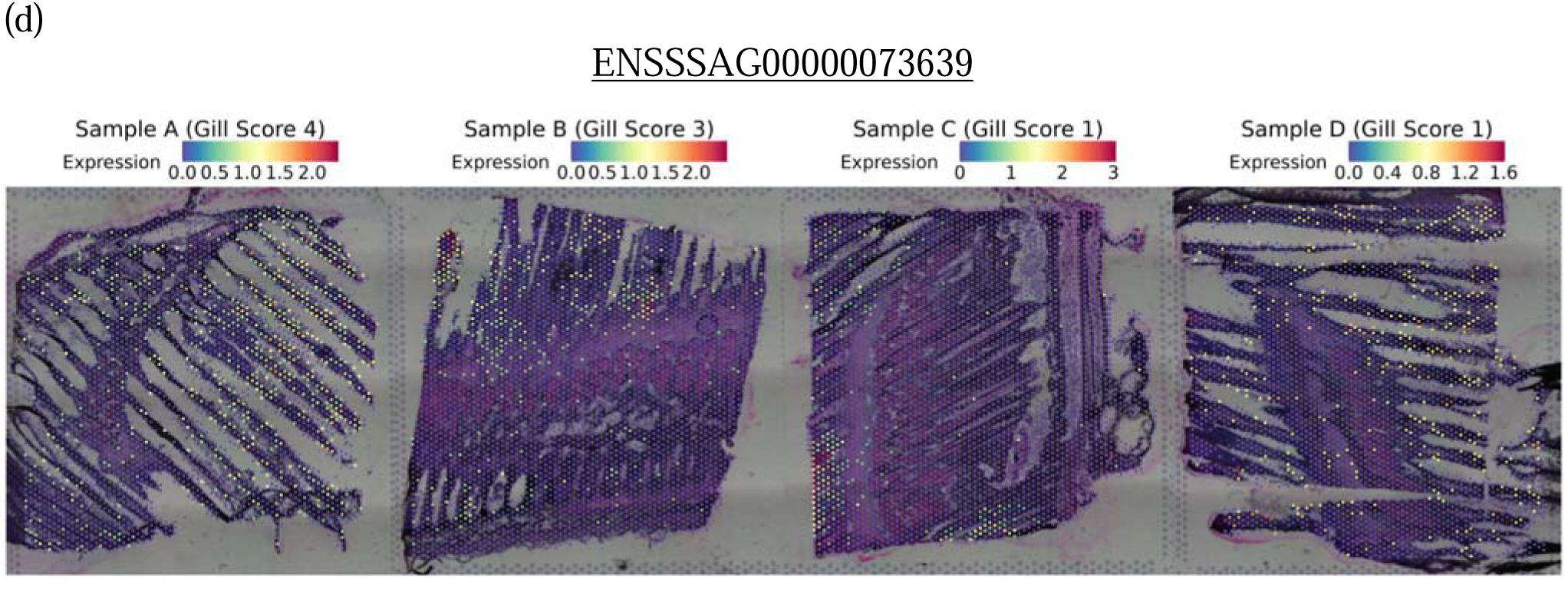
Spatial expression of genes encoding chemokines and interleukin-1: (a) *ccl28* (C-C motif chemokine 28; Ensembl ID: ENSSSAG00000065916; Cluster 0), (b) ENSSSAG00000033609 (C-C motif chemokine 19-like; Entrez Gene ID: LOC106563358; UniProt: A0A1S3L039; Cluster 10), (c) *cxl10* (C-X-C motif chemokine 10; Ensembl ID: ENSSSAG00000061390; Cluster 13) and (d) ENSSSAG00000045448 (interleukin-1; Entrez Gene ID: LOC106570815; UniProt: A0A1S3M701; Cluster 13). Each panel corresponds to one tissue section from Samples A to D. The colour intensity reflects normalised expression, using SCTransform, where higher values indicate greater relative expression following sequencing depth correction.

#### 3.3.3 Spatial Mapping of B Lymphocyte Markers and Antibody Production in the Gill

Spatial transcriptomic analysis of the gill identified three genes which, despite lacking official gene symbols, each exhibited strong spatial expression patterns and contained domains associated with immunoglobulin-like superfamilies. These genes are ENSSSAG00000108673 (**Figure 19a**), ENSSSAG00000100297 (**Figure 19b**), and ENSSSAG00000113740 (**Figure 19c**). To determine the specific type of immunoglobulin encoded by these genes, BLASTp sequence similarity searches were performed on their predicted amino acid sequences. The top BLASTp hits for ENSSSAG00000108673 included immunoglobulin lambda-1 light chain from *Salmo trutta* (XP_029631187.1, 89.80% identity) and immunoglobulin lambda-like polypeptide 1 precursor from *S. salar* (ACI66447.1, 91.25% identity). Similarly, ENSSSAG00000100297 showed highest similarity to immunoglobulin lambda-1 light chain isoform X2 from *S. salar* (XP_014057998.1, 100.00% identity). For ENSSSAG00000113740, the top hit was to an Ig lambda chain C region from *Oncorhynchus mykiss* (XP_036840305.1, 96.45% identity), with an Ig kappa chain V region from *S. salar* (ACM09401.1, 100.00% identity) also appearing in the top results. These BLASTp identifications indicate that ENSSSAG00000108673, ENSSSAG00000100297, and ENSSSAG00000113740 correspond to immunoglobulin light chains (either lambda or kappa types) which are essential components of antibodies. This supports the presence of B lymphocytes and rules out the possibility that these transcripts represent novel immunoglobulin-like transcripts (NILTs). Unlike NILTs, which are germline-encoded and do not form classical light or heavy chains, true immunoglobulin light chains are produced exclusively by B cells through V(D)J recombination (Wcisel *et al*., 2023). The localisation of these transcripts within the lamellae or gill arch region supports the presence of B lymphocytes in the gill-associated lymphoid tissue, indicating active humoral immune responses at this mucosal surface.

**Figure 19.**
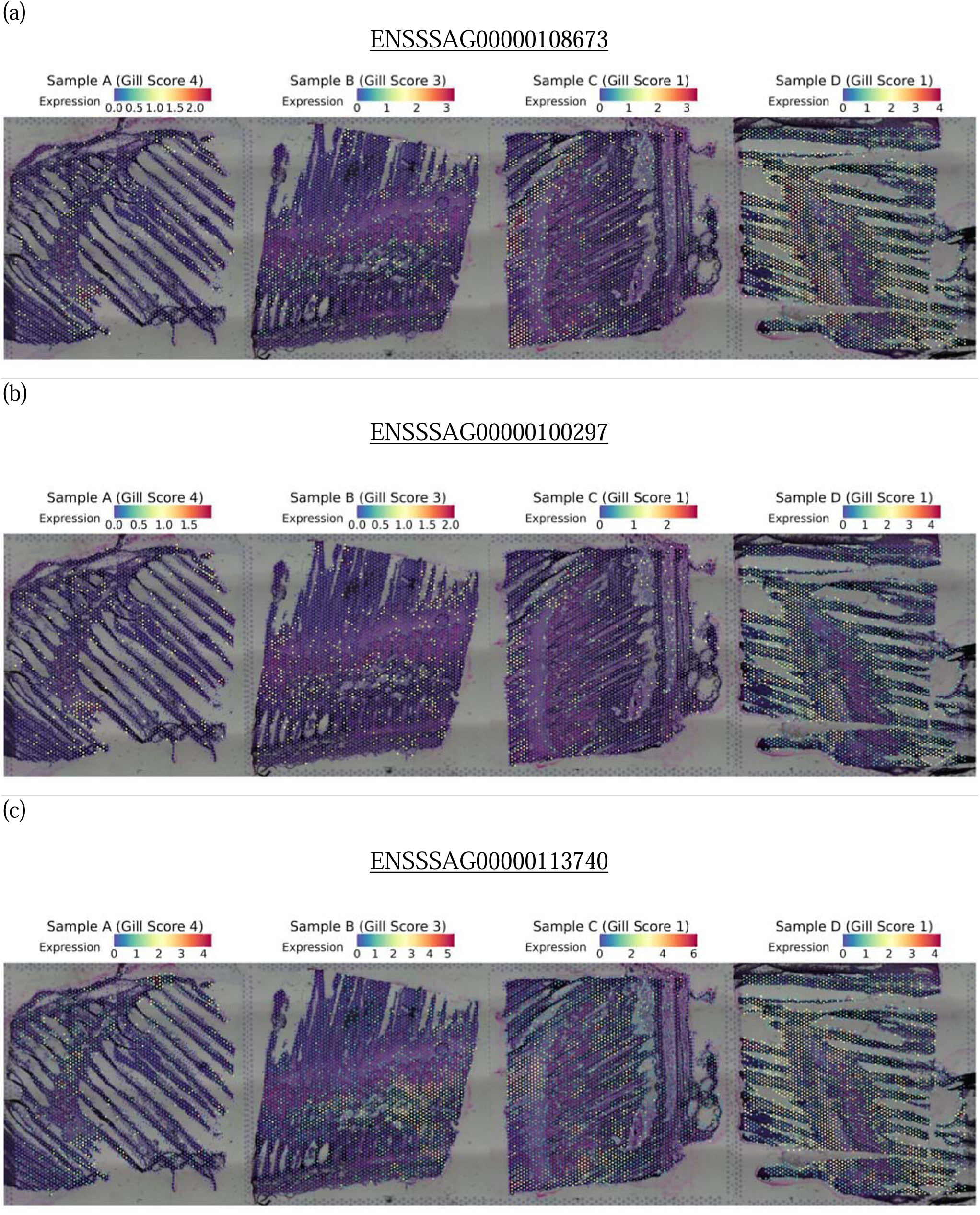
Spatial expression of immunoglobulin-like domain-containing genes: (a) ENSSSAG00000108673 (immunoglobulin-like domain superfamily; Pfam: immunoglobulin V-set and C1-set domains; Cluster 0), (b) ENSSSAG00000100297 (immunoglobulin-like domain superfamily; Pfam: immunoglobulin V-set domain; Cluster 0) and (c) ENSSSAG00000113740 (immunoglobulin-like domain superfamily; Pfam: immunoglobulin C1-set domain; Cluster 4). Each panel corresponds to one tissue section from Samples A to D. The colour intensity reflects normalised expression, using SCTransform, where higher values indicate greater relative expression following sequencing depth correction.

Based on the SVGs and DEGs, the gill of *S. salar* displays a spatially organised immune system capable of recognising pathogens, initiating inflammatory and antimicrobial responses, modulating immune cell activity, and maintaining tissue integrity despite environmental challenges. While spatial transcriptomics from 10X genomic Visium reveals the regional organisation of gene expression, one limitation is that each spatial spot (∼55 μm in diameter) may capture a mixture of distinct cell types. As a result, the gene expression profiles observed within a given cluster represent an aggregate signal from multiple cells, which can obscure the specific contributions of individual cell populations.

### 3.4 Integrating Spatial and Single Nuclei Transcriptomics in Gills

To address cellular heterogeneity within the gill-associated lymphoid tissue (GIALT) and to gain a higher-resolution understanding of the spatial arrangement of specific immune and structural components that support mucosal defence, our spatial transcriptomics dataset was integrated with single-nucleus RNA sequencing data from West et al. (2021). GIALT encompasses both the immune cell populations directly engaged in pathogen recognition and response, as well as the surrounding epithelial and structural framework that maintains the mucosal barrier and facilitates immune cell trafficking. Briefly, West et al. (2021) identified 20 distinct cell clusters in their sn-RNA experiment, including pavement cells, erythrocytes, four subtypes of vascular cells, pillar cells, mitochondrion-rich cells (MRCs) from both freshwater (FW) and seawater (SW) conditions, T cells, two subtypes of epithelial cells, goblet cells, non-differentiated cells, neuroepithelial cells, monocytes, lymphatic cells, dendritic cells, accessory cells, and fibrocytes. The integration was performed using RCTD in doublet mode, which constrains each spatial spot to contain at most two cell types and assumes that spots may represent transcriptomes from up to two distinct cells. Across the eight samples, an average of 51.2% of spatial spots were classified as singlets, representing areas dominated by a single cell type, 31.2% were categorised as doublets, and 17.6% were rejected due to unassigned cell type profiles (**Table 2**). Doublet spots reflect the close spatial proximity of two cell types, which may suggest potential cell–cell interactions; such interactions are especially important in the gill, where cooperation between structural and immune cells is essential for preserving barrier integrity and coordinating mucosal defence.

**Table 2.** Proportion of spatial spots that are classified as singlets, doublets and rejected for each sample following from RCTD.

| Sample | Singlets | Doublets | Rejects |
| --- | --- | --- | --- |
| Slide 1 Sample A | 1504 (45.8%) | 1331 (40.5%) | 452 (13.8%) |
| Slide 1 Sample B | 1942 (51.8%) | 1239 (33.1%) | 565 (15.1%) |
| Slide 1 Sample C | 2212 (57.6%) | 758 (19.7%) | 871 (22.7%) |
| Slide 1 Sample D | 1741 (48.8%) | 1392 (39.0%) | 438 (12.3%) |
| Slide 2 Sample A | 1341 (45.0%) | 831 (27.9%) | 808 (27.1%) |
| Slide 2 Sample B | 1708 (51.6%) | 1104 (33.4%) | 498 (15.0%) |
| Slide 2 Sample C | 2316 (59.5%) | 736 (18.9%) | 840 (21.6%) |
| Slide 2 Sample D | 1380 (50.6%) | 1005 (36.8%) | 344 (12.6%) |

The spatial distribution of all 20 recovered cell types is shown across the four gill samples in **Figure 20 to Figure 23**. Five cell types with particular immunological relevance were examined in further detail: T cells, dendritic cells, monocytes, lymphatic cells, and goblet cells. T cells were consistently localised at the filament and lamellar regions rather than the arch, and frequently co-occurred with goblet cells, erythrocytes, and vascular cells, positioning them within the peripheral epithelial-vascular mucosal interface. Dendritic cells clustered predominantly at the gill arch, while also extending into the filaments, and most frequently co-localised with vascular cells and erythrocytes, consistent with a sentinel role bridging antigen entry and vascular exchange. Monocytes were distributed diffusely across the gill without clear regional restriction, and were consistently found in close spatial association with erythrocytes and dendritic cells, reflecting flexible integration into vascular, epithelial, and immune niches. Lymphatic cells, defined by *Prox1* expression following West *et al*. (2021), were abundant throughout both the arch and filaments and were consistently found alongside dendritic, epithelial, vascular, and erythrocyte populations. Goblet cells were concentrated within the filament and lamellar region and most frequently co-localised with epithelial cells, T cells, erythrocytes, and vascular cells. Full singlet/doublet proportions and detailed co-localisation frequencies for each cell type, per sample, are provided in the **Supplementary Note 11.**

**Figure 20.**
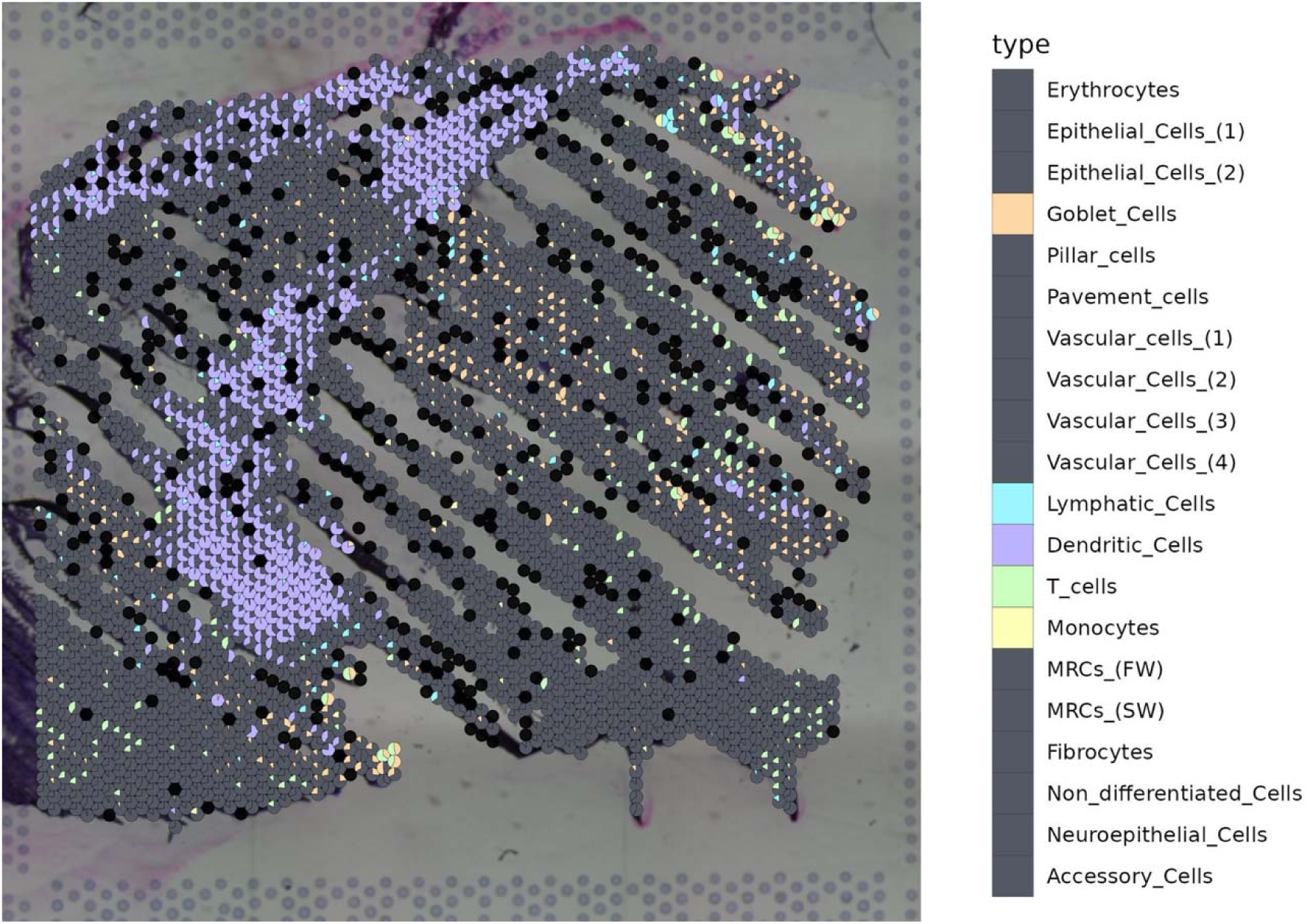
Spatial distribution of cell types in Atlantic salmon gills for sample A (gross gill score: 4), with each spot on the gills visualised as a pie chart. The pie chart distribution is based on the cell type proportions for each spot, as assigned by the RCTD deconvolution algorithm. This assignment was performed by integrating the spatial data with single-nuclei transcriptomics data from West *et al*. (2021) on Atlantic salmon gill cells. Each cell type is coloured as indicated in the legend. Spots identified as a singlet are shown as a single colour, while doublets (spots with up to two assigned cell types) are represented by multi-coloured pie charts. Spots that were rejected due to unassigned cell type profiles are shown as black dots.

**Figure 21.**
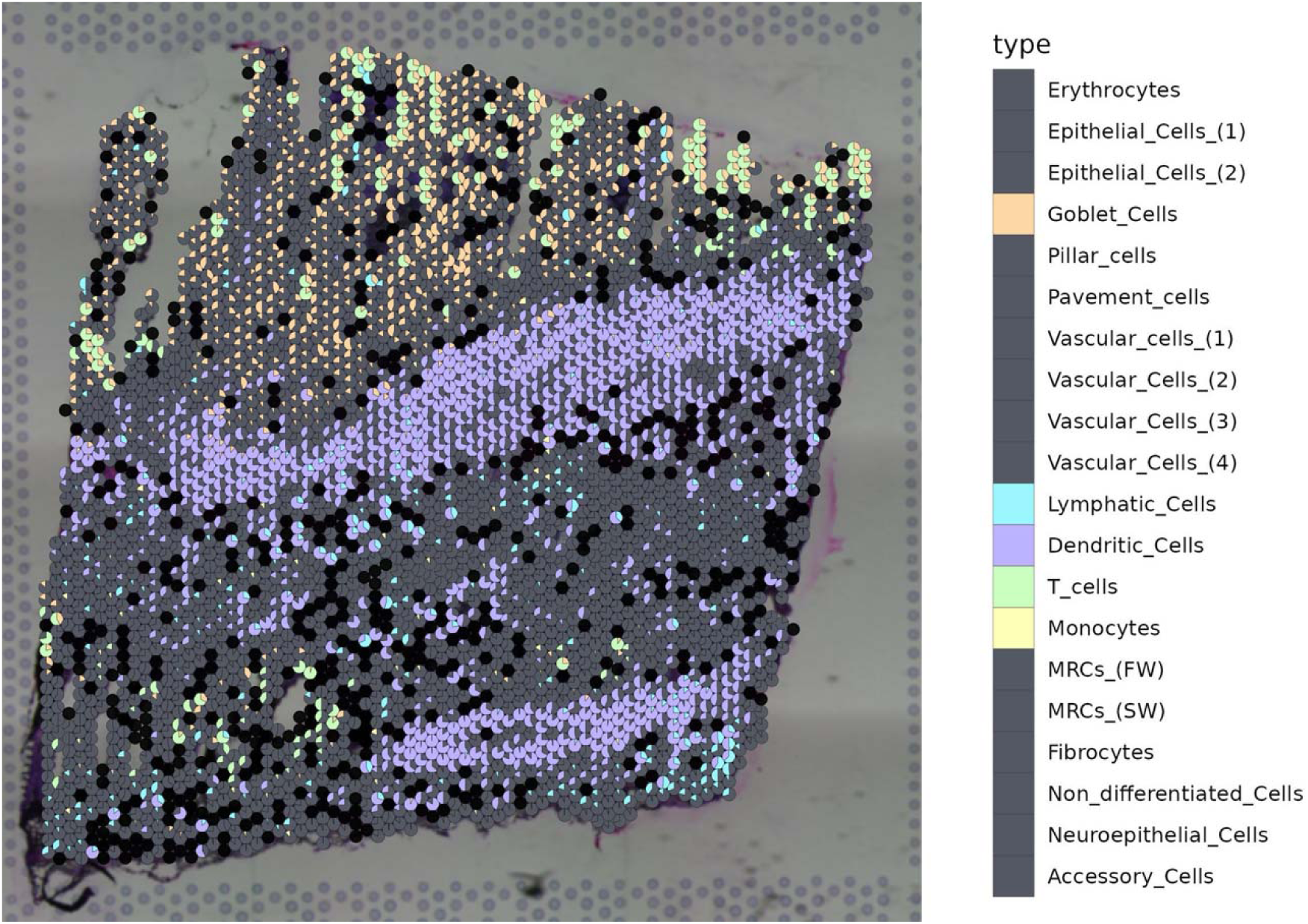
Spatial distribution of cell types in Atlantic salmon gills for sample B (gross gill score: 3), with each spot on the gills visualised as a pie chart. The pie chart distribution is based on the cell type proportions for each spot, as assigned by the RCTD deconvolution algorithm. This assignment was performed by integrating the spatial data with single-nuclei transcriptomics data from West et al. (2021) on Atlantic salmon gill cells. Each cell type is coloured as indicated in the legend. Spots identified as a singlet are shown as a single colour, while doublets (spots with up to two assigned cell types) are represented by multi-coloured pie charts. Spots that were rejected due to unassigned cell type profiles are shown as black dots.

**Figure 22.**
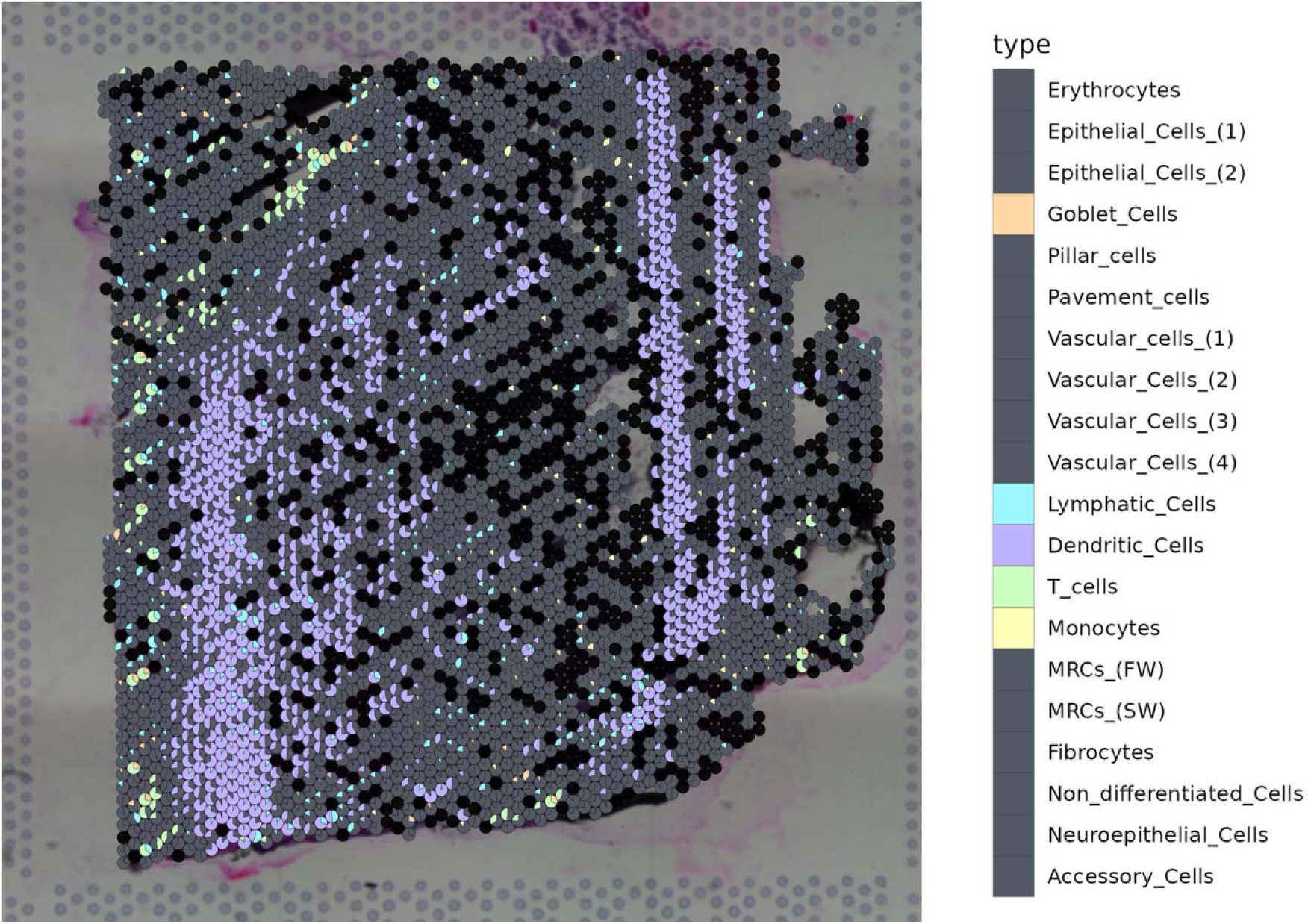
Spatial distribution of cell types in Atlantic salmon gills for sample C (gross gill score: 1), with each spot on the gills visualised as a pie chart. The pie chart distribution is based on the cell type proportions for each spot, as assigned by the RCTD deconvolution algorithm. This assignment was performed by integrating the spatial data with single-nuclei transcriptomics data from West et al. (2021) on Atlantic salmon gill cells. Each cell type is coloured as indicated in the legend. Spots identified as a singlet are shown as a single colour, while doublets (spots with up to two assigned cell types) are represented by multi-coloured pie charts. Spots that were rejected due to unassigned cell type profiles are shown as black dots.

**Figure 23.**
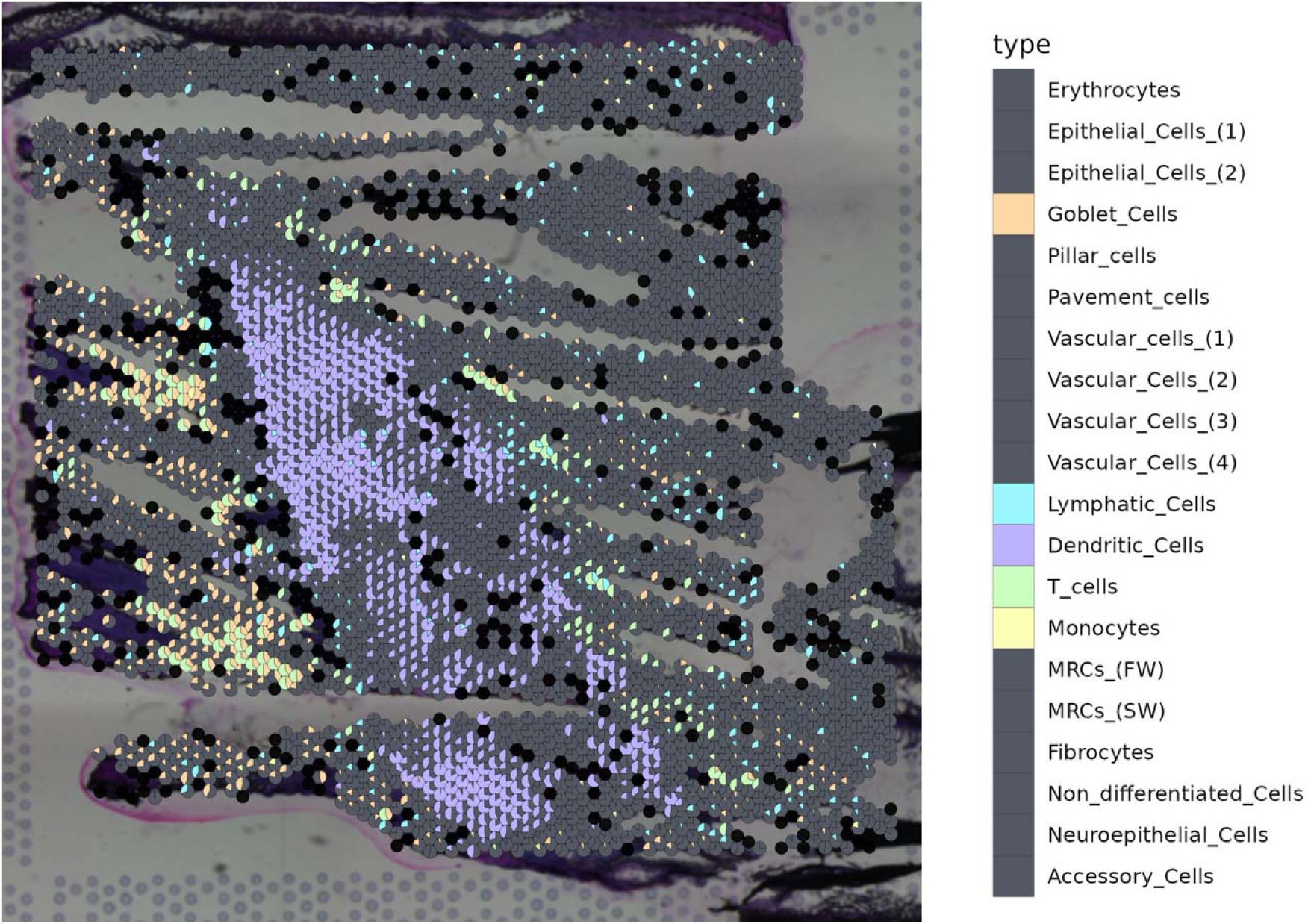
Spatial distribution of cell types in Atlantic salmon gills for sample D (gross gill score: 1), with each spot on the gills visualised as a pie chart. The pie chart distribution is based on the cell type proportions for each spot, as assigned by the RCTD deconvolution algorithm. This assignment was performed by integrating the spatial data with single-nuclei transcriptomics data from West et al. (2021) on Atlantic salmon gill cells. Each cell type is coloured as indicated in the legend. Spots identified as a singlet are shown as a single colour, while doublets (spots with up to two assigned cell types) are represented by multi-coloured pie charts. Spots that were rejected due to unassigned cell type profiles are shown as black dots.

## 4 Discussion

This study presents the first spatial transcriptomic atlas of the Atlantic salmon gill and, to our knowledge, the first application of this technology to a teleost gill affected by *Neoparamoeba perurans*. Our study design, a natural exposure system, does not allow us to test which elements of the gill immunological repertoire are specifically up- or downregulated in response to the parasite. It does, however, provide a fascinating snapshot of the infected gill to form the basis of future work.

Five main findings emerge. First, the infected gill comprised fourteen spatial clusters resolved into four anatomical compartments with coherent functional signatures: filament and lamellar epithelium, gill arch, vasculature and skeletal muscle. Second, adaptive immune signals are regionally organised within this architecture: immunoglobulin light chain transcripts and dendritic cells were enriched toward the gill arch, whereas antimicrobial effectors and several pattern recognition receptors were distributed along the filaments. Third, the filament epithelium carries the strongest pro-inflammatory and antimicrobial signals in the dataset, including IL-1, *cxcl10*, cathelicidin and the neutrophil marker *mpx*. Fourth, T cells mapped to the filament and lamellae rather than the arch, the same compartment in which chemokines were enriched. Fifth, within the limits of four fish, we found no consistent association between cellular composition and gill score.

The compartments recovered by unsupervised clustering correspond closely to known gill anatomy. Keratin-rich epithelium and ionocyte transporters (*nhe3*, *clic1*, *trpv6*) define the filament and lamellae (Leguen *et al*., 2015; Su *et al*., 2024). Connective tissue, smooth muscle and lymphatic markers define the arch (Jackson *et al*., 2001; Liu and Li, 2022). Erythroid and skeletal muscle signatures are confined to vessels and muscle fibres. This correspondence indicates that the clusters are anatomically meaningful, and it provides the frame within which the immune signals below can be interpreted.

Gill-associated lymphoid tissue (GIALT) comprises a diffuse component, with immune cells scattered through the filament and lamellar epithelium, and organised components. The organised components are the interbranchial lymphoid tissue (ILT) and the amphibranchial lymphoid tissue (ALT) near the base of the filaments (Dalum *et al*., 2021; Haugarvoll *et al*., 2008; Koppang *et al*., 2015). Our data are consistent with this organisation, although they do not fully resolve it. Immunoglobulin light chain transcripts, which in teleosts are restricted to B cells (Perdiguero *et al*., 2021), were enriched toward the gill arch. Dendritic cells also mapped predominantly to the arch, and the lymphatic endothelial marker *lyve1* (*xlkd1*) was enriched in an arch cluster. Whether this corresponds to the ILT, the ALT or both requires histological confirmation that the 55 µm spot size and plane of section do not provide. Lymphatic endothelial cells identified by deconvolution were, by contrast, distributed across both arch and filaments. Antimicrobial effectors, including cathelicidin and a lysozyme-like gene, localised to the filaments and lamellae, as did the rhamnose-binding lectin *sal* and one C-type lectin. This is consistent with a diffuse surveillance layer at the primary site of parasite contact. The presence of B cell transcripts fits with the local humoral activity reported in AGD. Previous studies found fluctuating mucosal IgM during outbreaks (Marcos-López *et al*., 2017) and increased ILT lymphocyte density during early infection (Norte dos Santos *et al*., 2014). Whether infection expands this organised compartment cannot be determined without uninfected controls, and direct comparison with naïve gills is a clear priority for future work.

The histological hallmarks of AGD include epithelial hyperplasia, lamellar fusion, mucous cell hyperplasia, and infiltration of neutrophils and macrophages (Adams and Nowak, 2001, 2004). The filament epithelium was the focus of inflammatory and antimicrobial gene expression in our dataset. IL-1, *cxcl10* and cathelicidin were enriched in cluster 13, and the neutrophil marker *mpx* in cluster 11. IL-1β has previously been localised to AGD lesions (Morrison et al., 2012; Botwright et al., 2021b), and neutrophil infiltration is a consistent histological feature of the disease (Adams *et al*., 2004; Adams and Nowak, 2001). However, these genes also contribute to baseline mucosal defence, and without uninfected controls we cannot distinguish an infection-induced response from constitutive expression. Mucin genes did not feature among cluster markers despite mucous cell hyperplasia being characteristic of AGD (Marcos-López *et al*., 2018; Valdenegro-Vega *et al*., 2014). This probably reflects the dispersed distribution of mucous cells at 55 µm resolution rather than their absence. Linking these signals to the parasite directly was not possible here. Amoebae were not observed in the H&E-stained sections, despite all four fish being qPCR-positive with visible gross lesions. *N. perurans* is fragile and prone to rupture on freezing (Robertson, 2024), and amoebae were most likely lysed and lost from the unfixed cryosections during staining; the same processing also obscured lamellar fusion. In addition, the absence of a complete, uncontaminated *N. perurans* genome prevented reliable identification of amoeba-derived transcripts. Fixation-compatible spatial platforms, combined with *in situ* detection of the parasite and a curated reference genome, would allow the transcriptional basis of AGD lesions to be tested directly.

T cells were resolved by deconvolution against the *cd3e*-defined population of West *et al*. (2021). They localised to the filament and lamellae rather than the arch, and frequently shared spots with goblet, vascular and erythroid cells. This places T cells at the epithelial surface of the filament, the site of parasite attachment, and not only within organised lymphoid tissue. Two chemokines enriched in the same compartment could contribute to their recruitment. A CCL19-like chemokine, enriched in cluster 10, mediates lymphocyte chemotaxis in teleosts (Bjørgen *et al*., 2019; Chen *et al*., 2018). *cxcl10*, enriched in cluster 13, belongs to the IFN-γ-inducible CXCL9–11/CXCR3 axis that recruits T cells and macrophages (Valdés *et al*., 2024). We did not test whether T cell-assigned spots coincide with these clusters. The presence of a Th1-associated chemokine is notable, given that previous studies have described Th2-type cytokine expression (*il4/13*) in AGD-affected gills (Botwright *et al*., 2021; Marcos-López *et al*., 2018). The reference annotation does not resolve T cell subsets, so the function of these cells remains open.

Samples A and B (gill scores 4 and 3) and C and D (score 1) spanned a range of clinical severity. However, no cell–cell association was consistently enriched in both severe samples relative to both mild samples, and variation in doublet composition appeared largely sample-specific. With two fish per group this absence of pattern is uninformative rather than negative. There is also a problem of scale: gross gill score summarises the whole arch, whereas each section samples a few millimetres of it, so section-level pathology need not track the clinical score. Sampling lesioned and non-lesioned regions within the same gill would remove this mismatch and is better suited to spatial approaches than between-fish severity comparisons.

Taken together, our findings align closely with the existing AGD literature. They localise IL-1 and neutrophil signatures to the filament epithelium where lesions form, B cell transcripts to the region of organised lymphoid tissue, and T cells to the mucosal surface, while adding the spatial context that bulk and single-cell studies cannot provide. A key strength of the study is the successful integration of spatial and single-nucleus data. Lifting the West *et al*. (2021) atlas across genome builds and transferring its annotations by deconvolution assigned over 80% of spots to one or two cell types. The resulting positions were anatomically plausible, showing that existing salmon single-cell resources can be reused to interpret spatial data without generating new reference atlases. The main limitations, four fish and no uninfected controls, mean that the atlas describes how immune function is organised in the infected gill rather than how infection alters it. Both are straightforward to address. Higher-resolution and fixation-compatible platforms such as Visium HD and Xenium, applied to paired lesioned and unaffected tissue from infected and naïve fish, would resolve individual lesions and separate infection-induced from constitutive responses. The same approach extends naturally to other gill diseases and to evaluating treatments, vaccines and selective breeding at the tissue level. The framework established here provides the anatomical, cellular and immunological reference for that work.

## 7 Author Contributions

The Author Contributions section is mandatory for all articles, including articles by sole authors. If an appropriate statement is not provided on submission, a standard one will be inserted during the production process. The Author Contributions statement must describe the contributions of individual authors referred to by their initials and, in doing so, all authors agree to be accountable for the content of the work. Please see <u>here</u> for full authorship criteria.

## 9 Acknowledgments

The authors thank the animal care staff at the Marine Institute, Galway, Ireland, for their expert husbandry and assistance with the research animals. We also thank the reviewers for their constructive comments on the manuscript.

## 10 Data Availability Statement

The raw and processed sequencing data generated in this study have been deposited in the NCBI Gene Expression Omnibus (GEO) and will be publicly available upon publication of this manuscript. The accession number will be provided prior to final publication.

## Reference

Adams, M.B., Ellard, K. and Nowak, B.F. (2004), “Gross pathology and its relationship with histopathology of amoebic gill disease (AGD) in farmed Atlantic salmon, Salmo salar L.”, Journal of Fish Diseases, Vol. 27 No. 3, pp. 151–161, doi: 10.1111/J.1365-2761.2004.00526.X

Adams, M.B. and Nowak, B.F. (2001), “Distribution and structure of lesions in the gills of Atlantic salmon, Salmo salar L., affected with amoebic gill disease”, Journal of Fish Diseases, Vol. 24 No. 9, pp. 535–542, doi: 10.1046/j.1365-2761.2001.00330.x

Andresen, A.M.S., Taylor, R.S., Grimholt, U., Daniels, R.R., Sun, J., Dobie, R., Henderson, N.C., et al. (2024), “Mapping the cellular landscape of Atlantic salmon head kidney by single cell and single nucleus transcriptomics”, *Fish and Shellfish Immunology*, Academic Press, Vol. 146, doi: 10.1016/j.fsi.2024.109357

Bjørgen, H., Løken, O.M., Aas, I.B., Fjelldal, P.G., Hansen, T., Austbø, L. and Koppang, E.O. (2019), “Visualization of CCL19-like transcripts in the ILT, thymus and head kidney of Atlantic salmon (Salmo salar L.)”, *Fish & Shellfish Immunology*, Academic Press, Vol. 93, pp. 763–765, doi: 10.1016/J.FSI.2019.08.044

Bjørgen, H., Malik, S., Rimstad, E., Vaadal, M., Nyman, I.B., Koppang, E.O. and Tengs, T. (2024), “Cellular heterogeneity in red and melanized focal muscle changes in farmed Atlantic salmon (Salmo salar) visualized by spatial transcriptomics”, *Cell and Tissue Research*, Springer Science and Business Media Deutschland GmbH, Vol. 395 No. 2, pp. 199–210, doi: 10.1007/s00441-023-03850-x

Botwright, N.A., Mohamed, A.R., Slinger, J., Lima, P.C. and Wynne, J.W. (2021), “Host-Parasite Interaction of Atlantic salmon (Salmo salar) and the Ectoparasite Neoparamoeba perurans in Amoebic Gill Disease”, *Frontiers in Immunology*, Frontiers Media S.A., Vol. 12, p. 672700, doi: 10.3389/FIMMU.2021.672700/XML/NLM

Botwright, N.A., Rusu, A., English, C.J., Hutt, O. and Wynne, J.W. (2020), “A High Throughput Viability Screening Method for the Marine Ectoparasite Neoparamoeba perurans: Neoparamoeba perurans High Throughput Screening Method”, *Protist*, Elsevier GmbH, Vol. 171 No. 6, doi: 10.1016/j.protis.2020.125773

Brown, G.D., Willment, J.A. and Whitehead, L. (2018), “C-type lectins in immunity and homeostasis”, *Nature Reviews. Immunology*, Nat Rev Immunol, Vol. 18 No. 6, pp. 374–389, doi: 10.1038/S41577-018-0004-8

Cable, D.M., Murray, E., Zou, L.S., Goeva, A., Macosko, E.Z., Chen, F. and Irizarry, R.A. (2022), “Robust decomposition of cell type mixtures in spatial transcriptomics”, *Nature Biotechnology*, Nature Research, Vol. 40 No. 4, pp. 517–526, doi: 10.1038/S41587-021-00830-W;SUBJMETA=114,2019,212,2415,61,631;KWRD=BIOTECHNOLOGY,GENOMICS,STATISTICAL+METHODS,TRANSCRIPTOMICS

Chang, C.I., Pleguezuelos, O., Zhang, Y.A., Zou, J. and Secombes, C.J. (2005), “Identification of a Novel Cathelicidin Gene in the Rainbow Trout, Oncorhynchus mykiss”, *Infection and Immunity*, American Society for Microbiology, Vol. 73 No. 8, p. 5053, doi: 10.1128/IAI.73.8.5053-5064.2005

Chang, C.I., Zhang, Y.A., Zou, J., Nie, P. and Secombes, C.J. (2006), “Two Cathelicidin Genes Are Present in both Rainbow Trout (Oncorhynchus mykiss) and Atlantic Salmon (Salmo salar)”, Antimicrobial Agents and Chemotherapy, Vol. 50 No. 1, p. 185, doi: 10.1128/AAC.50.1.185-195.2006

Chen, F., Lu, X.J., Nie, L., Ning, Y.J. and Chen, J. (2018), “Molecular characterization of a CC motif chemokine 19-like gene in ayu (Plecoglossus altivelis) and its role in leukocyte trafficking”, *Fish & Shellfish Immunology*, Academic Press, Vol. 72, pp. 301–308, doi: 10.1016/J.FSI.2017.11.012

Chen, H., Chen, X., Song, T.Y. and Ge, J.Q. (2022), “Expression Profiles of Zebrafish (Danio rerio) Lysozymes and Preparation of c-Type Lysozyme with High Bacteriolytic Activity against Vibrio vulnificus”, *Antibiotics (Basel, Switzerland)*, Antibiotics (Basel), Vol. 11 No. 12, doi: 10.3390/ANTIBIOTICS11121803

Dalum, A.S., Kraus, A., Khan, S., Davydova, E., Rigaudeau, D., Bjørgen, H., López-Porras, A., et al. (2021), “High-Resolution, 3D Imaging of the Zebrafish Gill-Associated Lymphoid Tissue (GIALT) Reveals a Novel Lymphoid Structure, the Amphibranchial Lymphoid Tissue”, *Frontiers in Immunology*, Frontiers Media S.A., Vol. 12, doi: 10.3389/fimmu.2021.769901

Denisenko, E., Guo, B.B., Jones, M., Hou, R., De Kock, L., Lassmann, T., Poppe, D., et al. (2020), “Systematic assessment of tissue dissociation and storage biases in single-cell and single-nucleus RNA-seq workflows”, *Genome Biology*, BioMed Central, Vol. 21 No. 1, pp. 1–25, doi: 10.1186/S13059-020-02048-6/FIGURES/4

D’Este, F., Benincasa, M., Cannone, G., Furlan, M., Scarsini, M., Volpatti, D., Gennaro, R., et al. (2016), “Antimicrobial and host cell-directed activities of Gly/Ser-rich peptides from salmonid cathelicidins”, *Fish & Shellfish Immunology*, Fish Shellfish Immunol, Vol. 59, pp. 456–468, doi: 10.1016/J.FSI.2016.11.004

Downes, J., Henshilwood, K., Collins, E., Ryan, A., O’Connor, I., Rodger, H., MacCarthy, E., et al. (2015), “A longitudinal study of amoebic gill disease on a marine Atlantic salmon farm utilising a real-time PCR assay for the detection of Neoparamoeba perurans”, *Aquaculture Environment Interactions*, Inter-Research Science Center, Vol. 7 No. 3, pp. 239–251, doi: 10.3354/aei00150

Efremova, M., Vento-Tormo, R., Park, J.E., Teichmann, S.A. and James, K.R. (2020), “Immunology in the Era of Single-Cell Technologies”, *Annual Review of Immunology*, Annual Reviews Inc., Vol. 38 No. Volume 38, 2020, pp. 727–757, doi: 10.1146/ANNUREV-IMMUNOL-090419-020340/CITE/REFWORKS

Gervais, O., Peñaloza, C., Gratacap, R., Papadopoulou, A., Beltrán, M., Henderson, N.C., Houston, R.D., et al. (2023), “Understanding host response to infectious salmon anaemia virus in an Atlantic salmon cell line using single-cell RNA sequencing”, *BMC Genomics*, BioMed Central Ltd, Vol. 24 No. 1, pp. 1–12, doi: 10.1186/S12864-023-09254-Z/PEER-REVIEW

Gjessing, M.C., Krasnov, A., Timmerhaus, G., Brun, S., Afanasyev, S., Dale, O.B. and Dahle, M.K. (2020), “The Atlantic Salmon Gill Transcriptome Response in a Natural Outbreak of Salmon Gill Pox Virus Infection Reveals New Biomarkers of Gill Pathology and Suppression of Mucosal Defense”, *Frontiers in Immunology*, Frontiers Media S.A., Vol. 11, p. 566294, doi: 10.3389/FIMMU.2020.02154/BIBTEX

Grindberg, R. V., Yee-Greenbaum, J.L., McConnell, M.J., Novotny, M., O’Shaughnessy, A.L., Lambert, G.M., Araúzo-Bravo, M.J., et al. (2013), “RNA-sequencing from single nuclei”, Proceedings of the National Academy of Sciences of the United States of America, Proc Natl Acad Sci U S A, Vol. 110 No. 49, pp. 19802–19807, doi: 10.1073/PNAS.1319700110,

Haugarvoll, E., Bjerkås, I., Nowak, B.F., Hordvik, I. and Koppang, E.O. (2008), “Identification and characterization of a novel intraepithelial lymphoid tissue in the gills of Atlantic salmon”, Journal of Anatomy, Vol. 213 No. 2, pp. 202–209, doi: 10.1111/j.1469-7580.2008.00943.x

Hieshima, K., Ohtani, H., Shibano, M., Izawa, D., Nakayama, T., Kawasaki, Y., Shiba, F., et al. (2003), “CCL28 Has Dual Roles in Mucosal Immunity as a Chemokine with Broad-Spectrum Antimicrobial Activity”, *The Journal of Immunology*, Oxford University Press (OUP), Vol. 170 No. 3, pp. 1452–1461, doi: 10.4049/JIMMUNOL.170.3.1452,

Jackson, D.G., Prevo, R., Clasper, S. and Banerji, S. (2001), “LYVE-1, the lymphatic system and tumor lymphangiogenesis”, *Trends in Immunology*, Elsevier, Vol. 22 No. 6, pp. 317–321, doi: 10.1016/S1471-4906(01)01936-6

Kashima, Y., Sakamoto, Y., Kaneko, K., Seki, M., Suzuki, Y. and Suzuki, A. (2020), “Single-cell sequencing techniques from individual to multiomics analyses”, *Experimental and Molecular Medicine*, Springer Nature, Vol. 52 No. 9, pp. 1419–1427, doi: 10.1038/S12276-020-00499-2;TECHMETA=23,38,39,45,91;SUBJMETA=1949,2254,514,61,631;KWRD=NEXT-GENERATION+SEQUENCING,RNA+SEQUENCING

Kim, G., Sohn, H., Omeka, W.K.M., Lim, C., Elvitigala, D.A.S. and Lee, J. (2023), “Functional characterization and expression analysis of c-type and g-like-type lysozymes in yellowtail clownfish (Amphiprion clarkii)”, *Fisheries and Aquatic Sciences*, Korean Society of Fisheries and Aquatic Sciences, Vol. 26 No. 3, pp. 188–203, doi: 10.47853/FAS.2023.E16

Koppang, E.O., Kvellestad, A. and Fischer, U. (2015), Mucosal Health in Aquaculture

Kube, P.D., Taylor, R.S. and Elliott, N.G. (2012), “Genetic variation in parasite resistance of Atlantic salmon to amoebic gill disease over multiple infections”, *Aquaculture*, Elsevier, Vol. 364–365, pp. 165–172, doi: 10.1016/J.AQUACULTURE.2012.08.026

Leguen, I., Le Cam, A., Montfort, J., Peron, S. and Fautrel, A. (2015), “Transcriptomic Analysis of Trout Gill Ionocytes in Fresh Water and Sea Water Using Laser Capture Microdissection Combined with Microarray Analysis”, *PLOS ONE*, Public Library of Science, Vol. 10 No. 10, p. e0139938, doi: 10.1371/JOURNAL.PONE.0139938

Li, B., Zhang, W., Guo, C., Xu, H., Li, L., Fang, M., Hu, Y., et al. (2022), “Benchmarking spatial and single-cell transcriptomics integration methods for transcript distribution prediction and cell type deconvolution”, Nature Methods 2022 19:6, Nature Publishing Group, Vol. 19 No. 6, pp. 662–670, doi: 10.1038/s41592-022-01480-9

Li, H., Sun, Y. and Sun, L. (2022), “A Teleost CXCL10 Is Both an Immunoregulator and an Antimicrobial”, *Frontiers in Immunology*, Frontiers Media S.A., Vol. 13, p. 917697, doi: 10.3389/FIMMU.2022.917697/BIBTEX

Liu, H. and Li, B. (2022), “Separation and identification of collagen peptides derived from enzymatic hydrolysate of Salmo salar skin and their anti-inflammatory activity in lipopolysaccharide (LPS)-induced RAW264.7 inflammatory model”, *Journal of Food Biochemistry*, John Wiley & Sons, Ltd, Vol. 46 No. 7, p. e14122, doi: 10.1111/JFBC.14122

Lowe, R., Shirley, N., Bleackley, M., Dolan, S. and Shafee, T. (2017), “Transcriptomics technologies”, *PLoS Computational Biology*, Public Library of Science, Vol. 13 No. 5, p. e1005457, doi: 10.1371/JOURNAL.PCBI.1005457

Magnadóttir, B. (2006), “Innate immunity of fish (overview)”, *Fish & Shellfish Immunology*, Academic Press, Vol. 20 No. 2, pp. 137–151, doi: 10.1016/J.FSI.2004.09.006

Marcos-López, M., Calduch-Giner, J.A., Mirimin, L., MacCarthy, E., Rodger, H.D., O’Connor, I., Sitjà-Bobadilla, A., et al. (2018), “Gene expression analysis of Atlantic salmon gills reveals mucin 5 and interleukin 4/13 as key molecules during amoebic gill disease”, Scientific Reports 2018 8:1, Nature Publishing Group, Vol. 8 No. 1, pp. 1–15, doi: 10.1038/s41598-018-32019-8

Marcos-López, M., Espinosa Ruiz, C., Rodger, H.D., O’Connor, I., MacCarthy, E. and Esteban, M.Á. (2017), “Local and systemic humoral immune response in farmed Atlantic salmon (Salmo salar L.) under a natural amoebic gill disease outbreak”, *Fish and Shellfish Immunology*, Academic Press, Vol. 66, pp. 207–216, doi: 10.1016/j.fsi.2017.05.029

Marcos-López, M. and Rodger, H.D. (2020), “Amoebic gill disease and host response in Atlantic salmon (Salmo salar L.): A review”, *Parasite Immunology*, Blackwell Publishing Ltd, Vol. 42 No. 8, doi: 10.1111/pim.12766

McLaughlin, C.N., Qi, Y., Quake, S.R., Luo, L. and Li, H. (2022), “Isolation and RNA sequencing of single nuclei from Drosophila tissues”, *STAR Protocols*, Cell Press, Vol. 3 No. 2, doi: 10.1016/J.XPRO.2022.101417

Morrison, R.N., Young, N.D. and Nowak, B.F. (2012), “Description of an Atlantic salmon (Salmo salar L.) type II interleukin-1 receptor cDNA and analysis of interleukin-1 receptor expression in amoebic gill disease-affected fish”, *Fish & Shellfish Immunology*, Academic Press, Vol. 32 No. 6, pp. 1185–1190, doi: 10.1016/J.FSI.2012.03.005

Moses, L. and Pachter, L. (2022), “Museum of spatial transcriptomics”, *Nature Methods*, Nature Research, 1 May, doi: 10.1038/s41592-022-01409-2

Muñoz-Atienza, E., Aquilino, C., Syahputra, K., Al-Jubury, A., Araújo, C., Skov, J., Kania, P.W., et al. (2019), “CK11, a Teleost Chemokine with a Potent Antimicrobial Activity”, *The Journal of Immunology*, Oxford Academic, Vol. 202 No. 3, pp. 857–870, doi: 10.4049/JIMMUNOL.1800568

Norte dos Santos, C.C., Adams, M.B., Leef, M.J. and Nowak, B.F. (2014), “Changes in the interbranchial lymphoid tissue of Atlantic salmon (Salmo salar) affected by amoebic gill disease”, *Fish and Shellfish Immunology*, Academic Press, Vol. 41 No. 2, pp. 600–607, doi: 10.1016/j.fsi.2014.10.003

Perdiguero, P., Morel, E., Díaz-Rosales, P. and Tafalla, C. (2021), “Individual B cells transcribe multiple rearranged immunoglobulin light chains in teleost fish”, *IScience*, Elsevier Inc., Vol. 24 No. 6, p. 102615, doi: 10.1016/J.ISCI.2021.102615/ATTACHMENT/E57530E4-0FA9-467B-91C6-4CE83F3FCDA1/MMC1.PDF

Pham, D., Tan, X., Balderson, B., Xu, J., Grice, L.F., Yoon, S., Willis, E.F., et al. (2023), “Robust mapping of spatiotemporal trajectories and cell–cell interactions in healthy and diseased tissues”, *Nature Communications*, Nature Research, Vol. 14 No. 1, pp. 1–25, doi: 10.1038/S41467-023-43120-6;SUBJMETA=114,1305,2391,327,4028,631,67,692;KWRD=CANCER+MICROENVIRONMENT,CELLULAR+SIGNALLING+NETWORKS,MACHINE+LEARNING

Power, C., Shirakashi, S., Bott, N.J. and Nowak, B.F. (2025), “Parasites in Aquaculture”, *Aquatic Parasitology: Ecological and Environmental Concepts and Implications of Marine and Freshwater Parasites*, Springer, Cham, pp. 595–620, doi: 10.1007/978-3-031-83903-0_23

Rao, A., Barkley, D., França, G.S. and Yanai, I. (2021), “Exploring tissue architecture using spatial transcriptomics”, Nature, Vol. 596 No. 7871, pp. 211–220, doi: 10.1038/s41586-021-03634-9

Robertson, B. (2024), Developing Protocols for Successful Cryopreservation of Neoparamoeba Perurans, the Causative Agent of Amoebic Gill Disease (Master’s Thesis)

Rozas-Serri, M. (2019), “Gill diseases in marine salmon aquaculture with an emphasis on amoebic gill disease”, *CAB Reviews: Perspectives in Agriculture, Veterinary Science, Nutrition and Natural Resources*, CABI International, Vol. 14, doi: 10.1079/PAVSNNR201914032

Saurabh, S. and Sahoo, P.K. (2008), “Lysozyme: an important defence molecule of fish innate immune system”, *Aquaculture Research*, John Wiley & Sons, Ltd, Vol. 39 No. 3, pp. 223–239, doi: 10.1111/J.1365-2109.2007.01883.X

Sepahi, A., Tacchi, L., Casadei, E., Takizawa, F., LaPatra, S.E. and Salinas, I. (2017), “CK12a, a CCL19-like Chemokine That Orchestrates both Nasal and Systemic Antiviral Immune Responses in Rainbow Trout”, *The Journal of Immunology*, Oxford University Press (OUP), Vol. 199 No. 11, pp. 3900–3913, doi: 10.4049/jimmunol.1700757

Su, Y., Yu, S.E., Sun, Y.X., Zhang, L., Tan, Y., Zhang, Y.Y., Wang, S., et al. (2024), “Genome-wide identification and quantification of salinity-responsive Na+/K+-ATPase α-subunits in three salmonids”, *Aquaculture*, Elsevier B.V., Vol. 582, doi: 10.1016/j.aquaculture.2023.740514

Sun, S.-S., Fu, P.-C., Zhou, X.-J., Cheng, Y.-W., Zhang, F.-Q., Chen, S.-L. and Gao, Q.-B. (2018), “The Complete Plastome Sequences of Seven Species in Gentiana sect. Kudoa (Gentianaceae): Insights Into Plastid Gene Loss and Molecular Evolution”, Frontiers in Plant Science, Vol. 9, doi: 10.3389/fpls.2018.00493

Sveen, L.R., Robinson, N., Krasnov, A., Daniels, R.R., Vaadal, M., Karlsen, C., Ytteborg, E., et al. (2023), “Transcriptomic landscape of Atlantic salmon (Salmo salar L.) skin”, G3: Genes, Genomes, Genetics, Genetics Society of America, Vol. 13 No. 11, doi: 10.1093/g3journal/jkad215

Takeuchi, O. and Akira, S. (2010), “Pattern Recognition Receptors and Inflammation”, *Cell*, Elsevier B.V., Vol. 140 No. 6, pp. 805–820, doi: 10.1016/J.CELL.2010.01.022/ASSET/4CC56ED2-02E5-40B4-AD20-359E7CB048EF/MAIN.ASSETS/GR5.JPG

Taylor, R.S., Muller, W.J., Cook, M.T., Kube, P.D. and Elliott, N.G. (2009), “Gill observations in Atlantic salmon (Salmo salar, L.) during repeated amoebic gill disease (AGD) field exposure and survival challenge”, Aquaculture, Vol. 290 No. 1–2, pp. 1–8, doi: 10.1016/j.aquaculture.2009.01.030

Valdenegro-Vega, V.A., Crosbie, P., Bridle, A., Leef, M., Wilson, R. and Nowak, B.F. (2014), “Differentially expressed proteins in gill and skin mucus of Atlantic salmon (Salmo salar) affected by amoebic gill disease”, *Fish and Shellfish Immunology*, Academic Press, Vol. 40 No. 1, pp. 69–77, doi: 10.1016/j.fsi.2014.06.025

Valdés, N., Espinoza, D., Pareja-Barrueto, C., Olate, N., Barraza-Rojas, F., Benavides-Larenas, A., Cortés, M., et al. (2024), “Expression and regulation of the CXCL9-11 chemokines and CXCR3 receptor in Atlantic salmon (Salmo salar)”, *Frontiers in Immunology*, Frontiers Media SA, Vol. 15, p. 1455457, doi: 10.3389/FIMMU.2024.1455457/BIBTEX

Wcisel, D.J., Dornburg, A., McConnell, S.C., Hernandez, K.M., Andrade, J., de Jong, J.L.O., Litman, G.W., et al. (2023), “A highly diverse set of novel immunoglobulin-like transcript (NILT) genes in zebrafish indicates a wide range of functions with complex relationships to mammalian receptors”, *Immunogenetics*, Springer Science and Business Media Deutschland GmbH, Vol. 75 No. 1, pp. 53–69, doi: 10.1007/S00251-022-01270-9/FIGURES/6

Wei, W., Wang, J., Min, Q., Jia, Z., Chen, K., Feng, H. and Zou, J. (2021), “CCL19 variants mediate chemotactic response via CCR7 in grass carp Ctenopharyngodon idella”, *Developmental and Comparative Immunology*, Elsevier Ltd, Vol. 122, doi: 10.1016/j.dci.2021.104127

West, A.C., Mizoro, Y., Wood, S.H., Ince, L.M., Iversen, M., Jørgensen, E.H., Nome, T., et al. (2021), “Immunologic Profiling of the Atlantic Salmon Gill by Single Nuclei Transcriptomics”, *Frontiers in Immunology*, Frontiers Media S.A., Vol. 12, doi: 10.3389/fimmu.2021.669889

Williams, C.G., Lee, H.J., Asatsuma, T., Vento-Tormo, R. and Haque, A. (2022), “An introduction to spatial transcriptomics for biomedical research”, *Genome Medicine*, BioMed Central Ltd, Vol. 14 No. 1, pp. 1–18, doi: 10.1186/S13073-022-01075-1/FIGURES/1

Xia, C., Fan, J., Emanuel, G., Hao, J. and Zhuang, X. (2019), “Spatial transcriptome profiling by MERFISH reveals subcellular RNA compartmentalization and cell cycle-dependent gene expression”, *Proceedings of the National Academy of Sciences of the United States of America*, National Academy of Sciences, Vol. 116 No. 39, pp. 19490–19499, doi: 10.1073/PNAS.1912459116/-/DCSUPPLEMENTAL

Young, N.D., Crosbie, P.B.B., Adams, M.B., Nowak, B.F. and Morrison, R.N. (2007), “Neoparamoeba perurans n. sp., an agent of amoebic gill disease of Atlantic salmon (Salmo salar)”, International Journal for Parasitology, Vol. 37 No. 13, pp. 1469–1481, doi: 10.1016/j.ijpara.2007.04.018

