## Supplementary Note 1 for "Spatial Transcriptomics Reveals Region-Specific Host Responses in Amoebic Gill Disease of Atlantic Salmon"

**Supplementary Note 1. Cryopreserved gill sample morphological integrity selection**

To ensure suitability for downstream spatial transcriptomics, gill samples were assessed for morphological integrity. Cryopreserved gills embedded in Optimal Cutting Temperature (OCT) compound were sectioned at a thickness of 10 µm using a cryostat. Sections were stained with Haematoxylin and Eosin (H&E) using a standardized protocol. Briefly, slides were processed on a Leica ST5020 autostainer. Sections were rinsed in water before immersion in Haematoxylin Z (RBA-4201-00A, CellPath) for 3 minutes, followed by rinsing in tap water and differentiation in 1% acid alcohol for 1 second. After a subsequent rinse, nuclear staining was enhanced using STWS, followed by a final rinse. The cytoplasm of sections was stained using eosin y (RRSP34-F, Atom Scientific) for 45 seconds. Sections were then rinsed in water, dehydrated through graded alcohols (70%, 100% ×2; 20821.365, VWR), cleared in xylene (×3; UN1307, Genta), and coverslipped on a Leica ST5030 coverslipper using DPX mountant (SEA-130-00A, CellPath). Once dried, slides were imaged at 20× magnification using a Leica Aperio AT2 digital slide scanner. Digital images were reviewed using ImageScope software (Leica) to evaluate tissue quality. Selection criteria for inclusion in spatial transcriptomics analysis included: (1) Clear preservation of gill morphology, including intact primary and secondary lamellae, and (2) minimal artefacts such as tearing, folding, freezing damage, or epithelial detachment.  Only samples meeting these quality standards were selected for 10X Visium spatial transcriptomics. Scanned H&E-stained sections from all 10 samples are shown below (**Figure S1.1)**. Of the ten cryopreserved gill samples, sample 1, sample 3, sample 4, sample 6, and sample 9 are excluded from the spatial gene expression profiling due to epithelial detachment were observed.

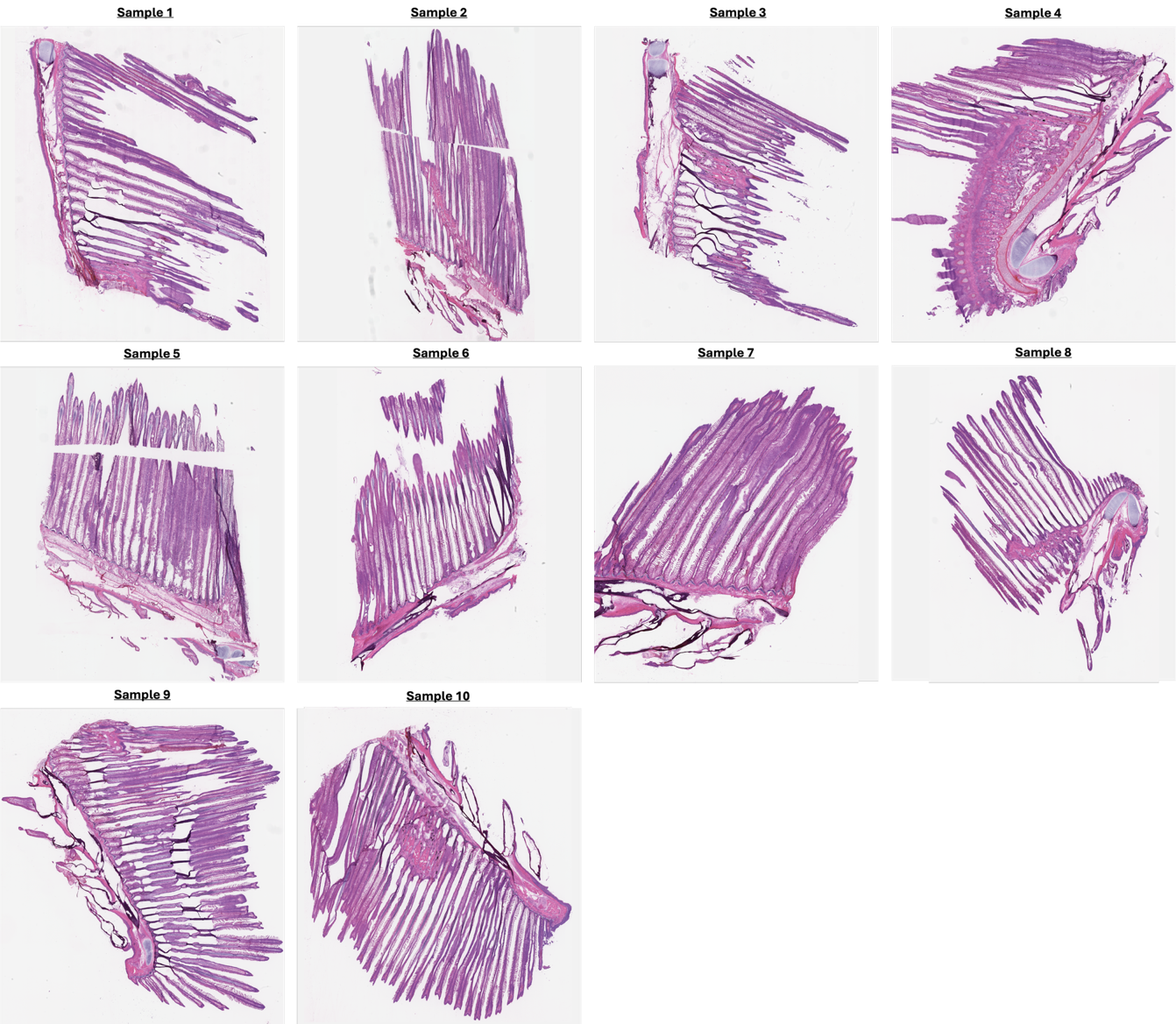

Figure S1.1	cryopreserved gill sample morphological integrity selection
