## Supplementary Note 2 for "Spatial Transcriptomics Reveals Region-Specific Host Responses in Amoebic Gill Disease of Atlantic Salmon"

**Supplementary Note 2.** RNA Integrity Number (RIN) Assessment for Sample Selection in 10X Visium Spatial Transcriptomics

Ten 10 µm cryosections from each cryopreserved sample (n = 10) were stored at −80 °C in Eppendorf tubes prior to RNA extraction, which was performed using the Qiagen RNeasy kit. Extracted RNA was then analysed using the Agilent Bioanalyzer with Nano chips to determine the RNA Integrity Number (RIN). The RIN is a standardized numerical value ranging from 1 to 10, where a score of 10 indicates highly intact RNA and 1 indicates fully degraded RNA. For optimal performance in 10X Visium spatial transcriptomics, 10X Genomics recommends using samples with a RIN of at least 7 to ensure RNA integrity suitable for spatial gene expression profiling. Of the ten cryopreserved gill samples, sample 2 (sample D), sample 5 (sample B), sample 7 (sample C), and sample 8 (sample A) having the RIN score above 9 were selected for spatial gene expression profiling. The figure below summarises the RIN scores for each of the 10 samples with the corresponding electropherograms shown to visualise RNA integrity.

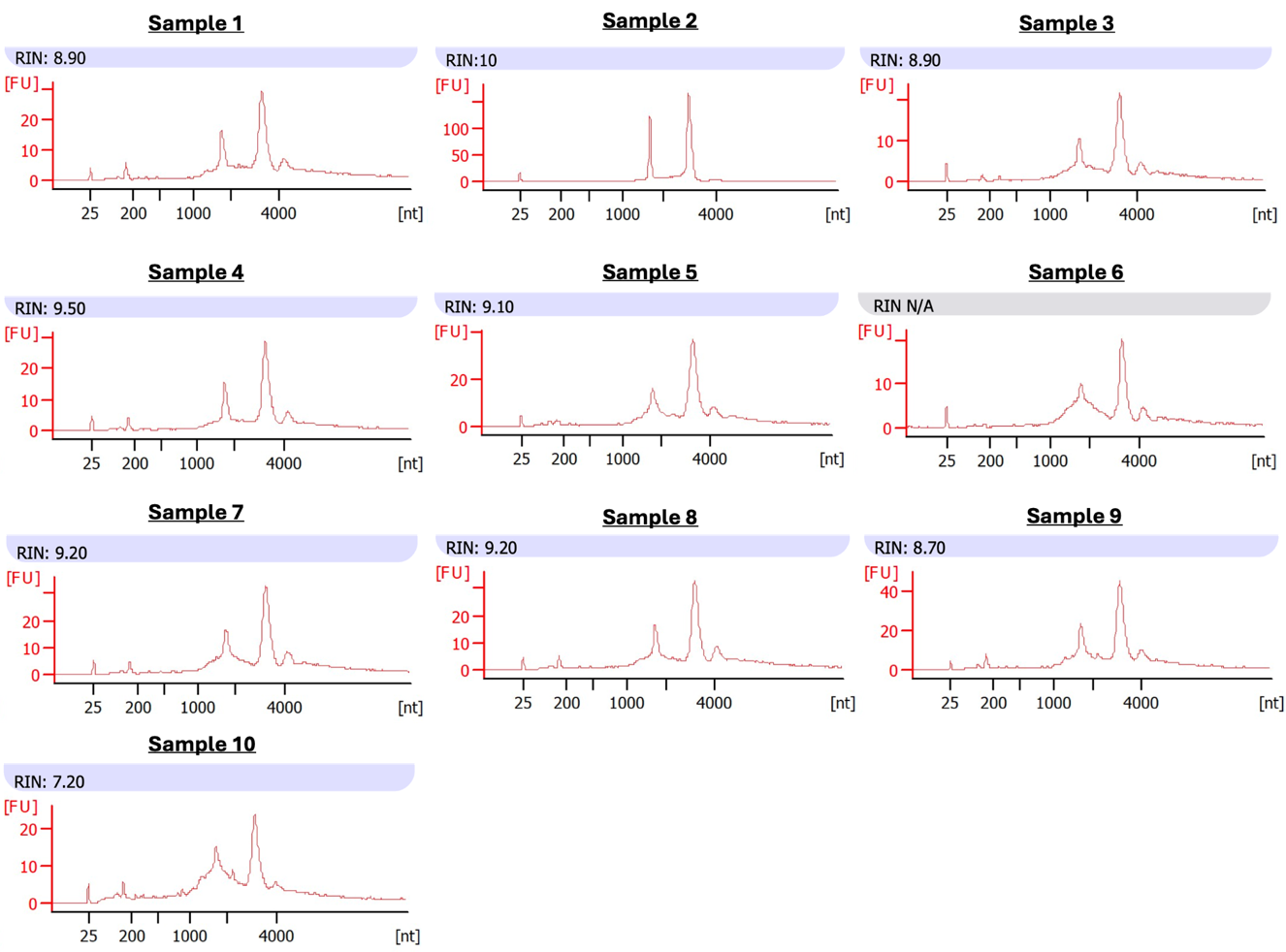

**Figure S2.1**   cryopreserved gill sample morphological integrity selection
