## Supplementary Note 3 for "Spatial Transcriptomics Reveals Region-Specific Host Responses in Amoebic Gill Disease of Atlantic Salmon"

**Supplementary Note 3. Permeability time determination**

Six minutes permeability time appears to be the brightest with gill filaments features clearly visualised under fluorescent macroscope. The gill sections on the 10X visium gene expression slide is therefore permeabilised for 6 minutes.

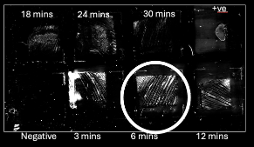

Figure S3.1 Cryopreserved gill sample morphological integrity selection
