## Supplementary Note 4 for "Spatial Transcriptomics Reveals Region-Specific Host Responses in Amoebic Gill Disease of Atlantic Salmon"

**Supplementary Note 4. Elbow Plot**

Elbow plot is a visualisation that plots explained variance against the number of principal components after principal component analysis, allowing for the selection of a subset of PCs that explain the majority of the variance without including excessive noise. The elbow point determined in **Figure S4.1** was 15.

 
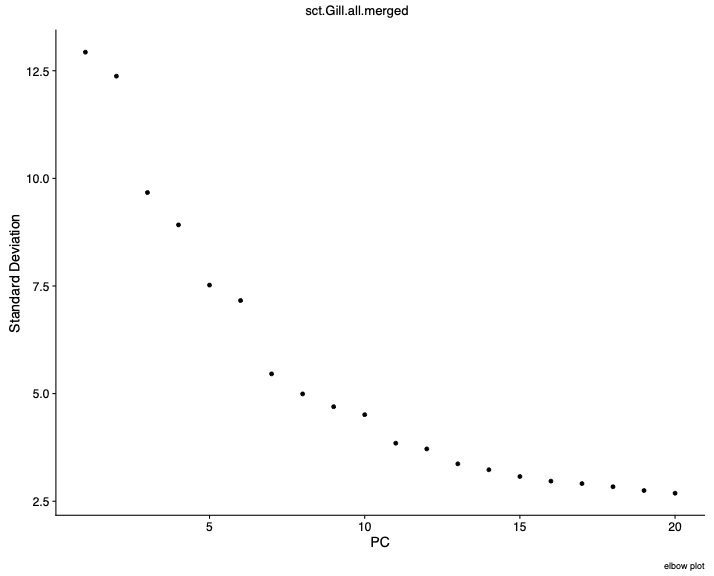

**Figure S4.1** Elbow plot
