## Supplementary Note 5 for "Spatial Transcriptomics Reveals Region-Specific Host Responses in Amoebic Gill Disease of Atlantic Salmon"

**Supplementary Note 5. histological interpretation of AGD was limited by cryopreservation artefacts**

When assessing the severity of AGD pathology in Haematoxylin and Eosin (H&E) stained gills from four *S. salar*, careful consideration was given to potential artefactual damage caused by cryopreservation and subsequent handling. Compared to formalin-fixed materials, cryopreservation is known to induce additional artifacts for histology. Consequently, it was not always possible to confirm all four hallmark criteria of AGD (epithelial hyperplasia, lamellar fusion, vesicle formation in hyperplastic regions, and the presence of amoebae) in every section. Although epithelial hyperplasia and lamellar fusion were observed across samples A to D (Figure S5.1 A to D), some of these fusions may have been exacerbated by cryo-induced damage. These observations were made by comparing areas that appeared similar to hyperplasia and fusion at higher magnification (Figure S5.1 F) to healthier regions (Figure S5.1 E). Furthermore, vesicles in hyperplastic tissue and amoebae were particularly difficult to observe due to their fragility and susceptibility to cryopreservation effects. Importantly, the focal nature of AGD pathology, with diseased and healthy regions often present within the same gill arch, further complicates histological assessment, especially when sample integrity is compromised.

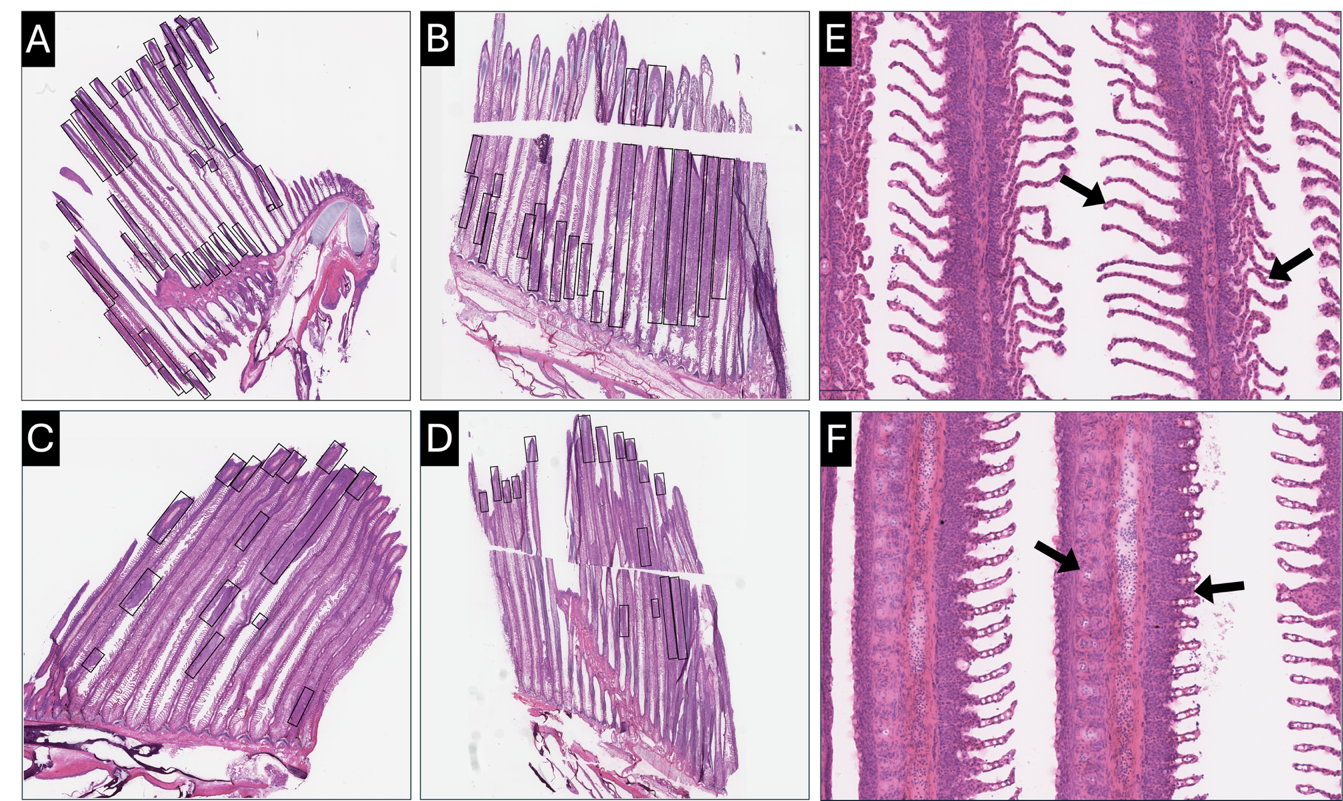

Figure S5.1 Haematoxylin and Eosin (H&E)-stained cryopreserved gill sections from Atlantic salmon (*Salmo salar*) affected by Amoebic Gill Disease (AGD) display varying degrees of gill pathology with severity categorised by gross gill scores and qPCR results. (A) Sample A, a section from a fish with a macroscopic gill score of 4. (B) Sample B, a section from a fish with a gill score of 3 (C) Sample C, a section from a fish with a gill score of 1. (D) Sample D, a section from a fish with a gill score of 1, showing moderate lamellar fusion with reduced interlamellar space and epithelial thickening. Rectangular outlines in A to D highlight regions of potential epithelial hyperplasia and secondary lamellar fusion, likely exacerbated by cryopreservation artifacts. (E) High magnification image of healthy gill lamellae, as highlighted by the arrows, showing distinct and separate secondary lamellae. (F) High magnification image of less healthy gill lamellae, as highlighted by the arrows, displaying extensive fusion and epithelial hyperplasia.

Given the challenges in definitively assessing AGD lesions from the histological images alone, disease severity was primarily classified based on gross gill scores (Figure 1) and confirmed by qPCR results (Figure 2). These combined data indicated samples A and B as severe AGD cases, with samples C and D showing milder pathology. While histopathological assessment is visually difficult, spatial transcriptomics might provide a valuable, high-resolution molecular perspective to further characterise these different severity levels. It is important to note that since the classification of severe versus milder cases was based on gross pathology rather than specific lesions on the spatial transcriptomics slides, caution is needed when making direct comparisons between these groups. The number of samples was constrained by practical considerations. Access to *Neoparamoeba perurans*-positive gills was limited to a small number of suitable fish within the available collection window, and the high cost of Visium spatial transcriptomics restricted the number of slides that could be processed. Technical replication was prioritised, with two slides prepared per gill region where feasible, to evaluate spatial and assay-level consistency. In contrast, biological replication, defined as sampling from independent fish, was limited. As a result, formal statistical comparisons between samples, such as severity-stratified differential gene expression analysis, were not conducted. Instead, observed differences are presented descriptively and interpreted as exploratory findings.
