## Supplementary Note 6 for "Spatial Transcriptomics Reveals Region-Specific Host Responses in Amoebic Gill Disease of Atlantic Salmon"

**Supplementary Note 6. Spatial Transcriptomics Data Quality Control**

To profile the transcriptomic landscape of gills in *S. salar* exhibiting varying gross severity of AGD, as determined by macroscopic scoring and qPCR quantification, the 10X Genomics Visium Spatial Gene Expression platform was employed. Four representative samples (sample A, B, C, and D) were selected for spatial transcriptomic analysis. For each sample, technical replicates were included to ensure data reliability and reproducibility (Figure S6.1).

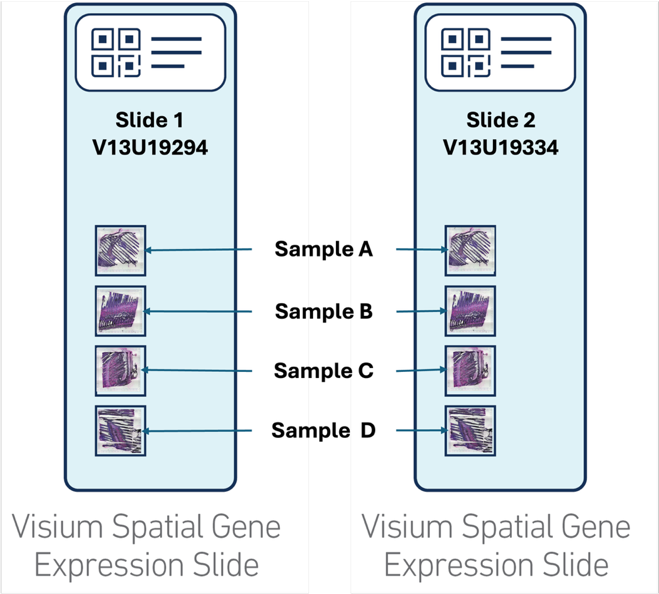

Figure S6.1 Layout of gills sections from Atlantic sample (*Salmo salar*) on two 10X Genomics Visium spatial gene expression slides used for spatial transcriptomic analysis. Each slide (V13U19294 and V13U19334) contains four gills sections representing different levels of AGD severity (Samples A, B, C, and D). This resulted in a total of eight spatial capture areas: V13U19294A (Slide 1 – Samples A), V13U19294B (Slide 1 – Samples B), V13U19294C (Slide 1 – Samples C), V13U19294D (Slide 1 – Samples D), V13U19334A (Slide 2 – Samples A), V13U19334B (Slide 2 – Samples B), V13U19334C (Slide 2 – Samples C), and V13U19334D (Slide 2 – Samples D).

All spatial transcriptomic libraries were sequenced to high depth, yielding between 238.8 and 245.3 million reads per sample across eight samples. Barcode and UMI detection rates were high, with valid barcodes ranging from 96.8 to 97.2% and valid UMIs from 99.0 to 99.4%. Base quality was uniformly high across samples, with Q30 scores exceeding 92.5% for barcodes, RNA reads, and UMIs. Across samples, each slide contained 4,992 tissue-covered spots, with per-spot median gene counts ranging from 484 to 1,290, and median UMI counts from 768 to 3,641. Total genes detected per sample ranged from 32,999 to 37,566.

To ensure unbiased downstream analysis, quality control filtering was applied by excluding spatial transcriptomics spots with fewer than 200 detected unique genes, as well as those not overlying tissue regions, determined by manual annotation in Loupe Browser. Such filtration retained a total of 27,356 high-quality spots across eight tissue sections (Figure S6.2)

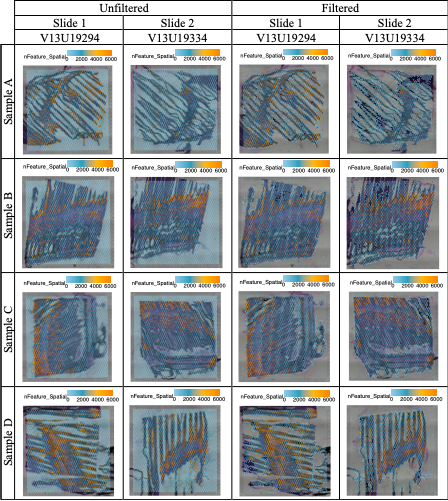

Figure S6.2 Spatial transcriptomics quality control across all spatial capture areas: V13U19294A, V13U19294B, V13U19294C, V13U19294D, V13U19334A, V13U19334B, V13U19334C, and V13U19334D. Spatial distribution of the number of unique genes (nFeature_Spatial) across four gills samples, each with technical replicates. Left panels show unfiltered data; right panels show the same tissues after quality control filtering. Spots with fewer than 200 detected genes or without tissue coverage (based on manual annotation) were excluded. Spatial feature plots (overlaid on H&E images) display the number of unique genes per spot. This filtering step effectively removes low-quality or outlier spots to ensure robust downstream analysis.
