## Supplementary Note 10 for "Spatial Transcriptomics Reveals Region-Specific Host Responses in Amoebic Gill Disease of Atlantic Salmon"

**Supplementary Note 10.** **Transcriptional Signatures of Skeletal Muscle and High Aerobic Metabolism in 3.2.3 Skeletal Muscle cluster (Cluster 3)**

A large proportion of the top marker genes in Cluster 3 are associated with striated muscle contraction. Notable genes linked to the contractile apparatus, including ***tnnc2.2*** (troponin C2, fast skeletal type, tandem duplicate 2; Ensembl ID: ENSSSAG00000066537; Cluster 3; S10.1), ***mlrb*** (myosin regulatory light chain 2B, cardiac muscle isoform; Ensembl ID: ENSSSAG00000062577; NCBI Gene: mlrb; Cluster 3; S10.2), and various actin isoforms such as ***acta1-like*** (actin alpha 1, skeletal muscle; Ensembl ID: ENSSSAG00000007690; Cluster 3; S10.3) and ***actc1a*** (actin alpha cardiac muscle 1a; Ensembl ID: ENSSSAG00000073229; Cluster 3; S10.4) show relatively high expression within this cluster. With both skeletal and cardiac muscle isoforms rank among the top marker genes in Cluster 3, this indicates a diverse muscle population or specialised isoforms adapted to the gill environment. The high expression of ***fhl1a*** (four and a half LIM domains 1a; Ensembl ID: ENSSSAG00000063443; Cluster 3; S10.5), a cytoskeletal protein commonly found in skeletal and cardiac muscle, further supports the muscular identity of this cluster.

***tnnc2* (ENSSSAG00000066537)**

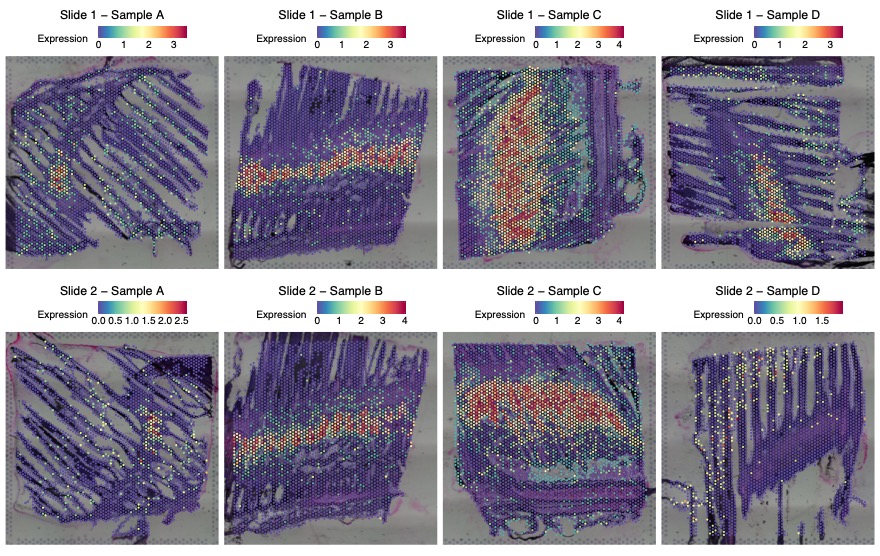

Figure S10.1 The spatial distribution of *tnnc2.2* (troponin C2, fast skeletal type, tandem duplicate 2; Ensembl ID: ENSSSAG00000066537) expression in eight tissue sections (Slide 1: Samples A to D; Slide 2: Samples A to D). Each panel corresponds to one tissue section, with expression values represented by colour intensity. The colour intensity reflects normalised expression, using SCTransform, where higher values indicate greater relative expression following sequencing depth correction.

***mlrb* (ENSSSAG00000062577)**

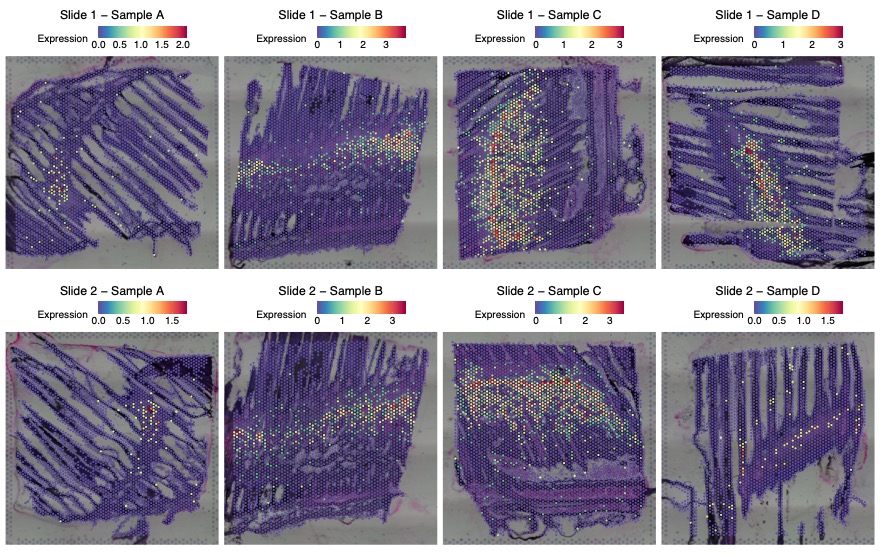

Figure S10.2 The spatial distribution of *mlrb* (myosin regulatory light chain 2B, cardiac muscle isoform; Ensembl ID: ENSSSAG00000062577; NCBI Gene: mlrb) expression in eight tissue sections (Slide 1: Samples A to D; Slide 2: Samples A to D). Each panel corresponds to one tissue section, with expression values represented by colour intensity. The colour intensity reflects normalised expression, using SCTransform, where higher values indicate greater relative expression following sequencing depth correction.

***acta1*-*like* (ENSSSAG00000007690)**

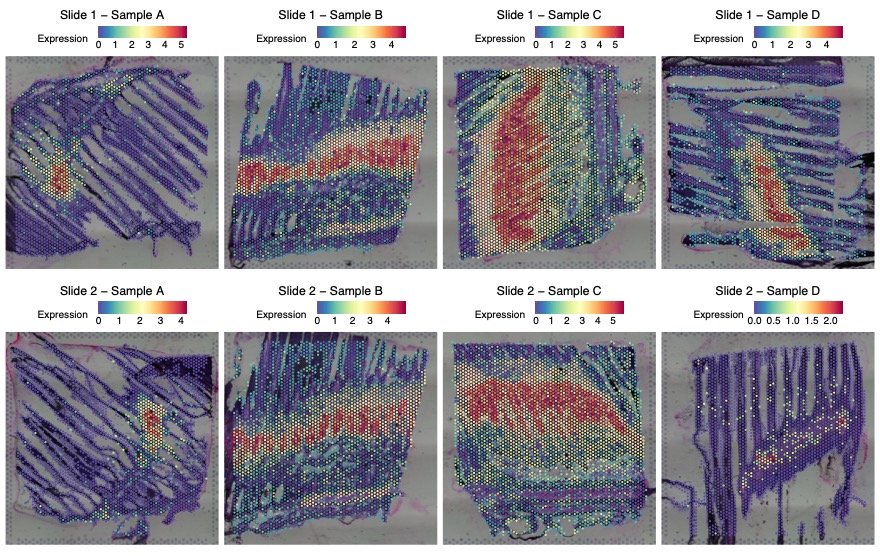

Figure S10.3 The spatial distribution of *acta1-like* (actin alpha 1, skeletal muscle; Ensembl ID: ENSSSAG00000007690) expression in eight tissue sections (Slide 1: Samples A to D; Slide 2: Samples A to D). Each panel corresponds to one tissue section, with expression values represented by colour intensity. The colour intensity reflects normalised expression, using SCTransform, where higher values indicate greater relative expression following sequencing depth correction.

***actc1a* (ENSSSAG00000073229)**

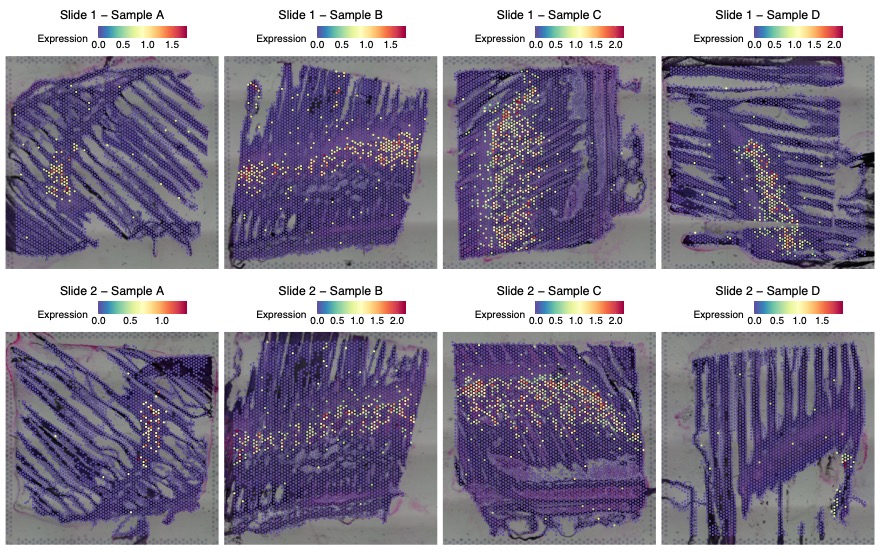

Figure S10.4 The spatial distribution of *actc1a* (actin alpha cardiac muscle 1a; Ensembl ID: ENSSSAG00000073229) expression in eight tissue sections (Slide 1: Samples A to D; Slide 2: Samples A to D). Each panel corresponds to one tissue section, with expression values represented by colour intensity. The colour intensity reflects normalised expression, using SCTransform, where higher values indicate greater relative expression following sequencing depth correction.

***fhl1a* (ENSSSAG00000063443)**

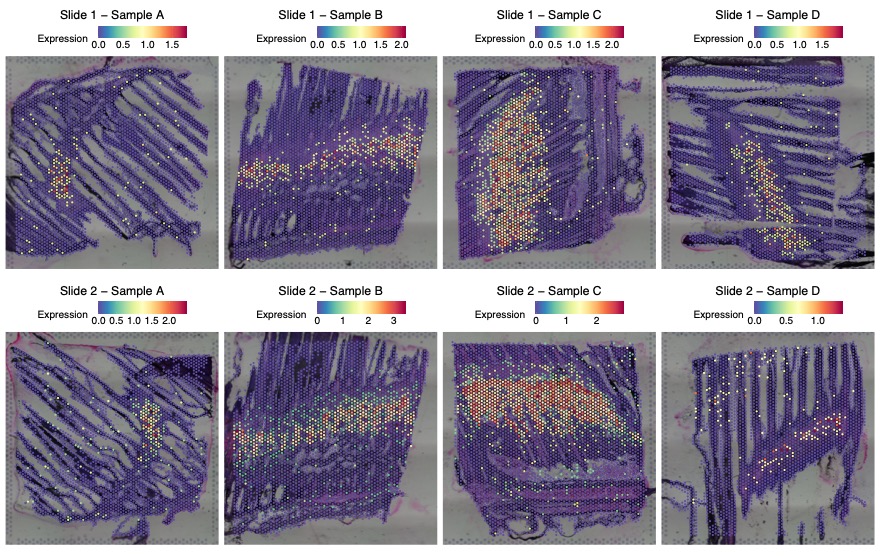

Figure S10.5 The spatial distribution of fhl1a (four and a half LIM domains 1a; Ensembl ID: ENSSSAG00000063443) expression in eight tissue sections (Slide 1: Samples A to D; Slide 2: Samples A to D). Each panel corresponds to one tissue section, with expression values represented by colour intensity. The colour intensity reflects normalised expression, using SCTransform, where higher values indicate greater relative expression following sequencing depth correction.

Complementing the contractile machinery, the top marker genes in Cluster 3 also display a strong signature of high-energy metabolism genes, reflecting the substantial ATP demands of muscle activity. Several isoforms of Glyceraldehyde-3-phosphate dehydrogenase were highly expressed in Cluster 3, including ***gapdh*** (Glyceraldehyde-3-phosphate dehydrogenase; Ensembl ID: ENSSSAG00000099398; Cluster 3; Figure S10.6), ***g3p*** (Glyceraldehyde-3-phosphate dehydrogenase; Ensembl ID: ENSSSAG00000067521; Cluster 3; Figure S10.7), and ***g3p*** (Glyceraldehyde-3-phosphate dehydrogenase; Ensembl ID: ENSSSAG00000073639; Cluster 3; Figure S10.8), all of which are enzymes that play a central role in glycolysis. Furthermore, multiple subunits of Cytochrome c oxidase, the terminal enzyme of the mitochondrial electron transport chain, were highly expressed in Cluster 3, such as ***cx6a1*** (cytochrome c oxidase subunit 6A, mitochondrial; Ensembl ID: ENSSSAG00000041369; Cluster 3; Figure S10.9) and ***cox4i2*** (cytochrome c oxidase subunit 4I2; Ensembl ID: ENSSSAG00000063792; Cluster 3; Figure S10.10). Such expression patterns indicate a reliance on efficient aerobic respiration for sustained energy production.

***gapdh* (ENSSSAG00000099398)**

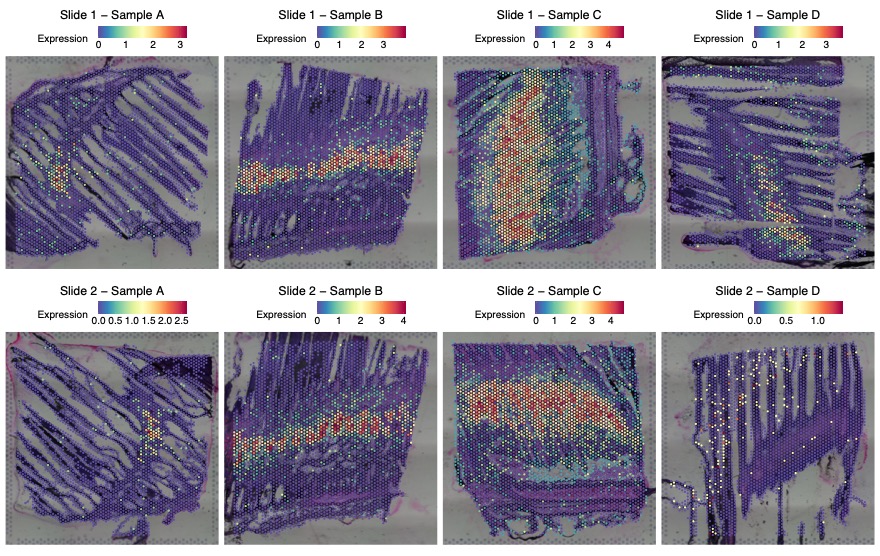

Figure S10.6 The spatial distribution of *gapdh* (Glyceraldehyde-3-phosphate dehydrogenase; Ensembl ID: ENSSSAG00000099398) expression in eight tissue sections (Slide 1: Samples A to D; Slide 2: Samples A to D). Each panel corresponds to one tissue section, with expression values represented by colour intensity. The colour intensity reflects normalised expression, using SCTransform, where higher values indicate greater relative expression following sequencing depth correction.

**g3g (ENSSSAG00000067521)**

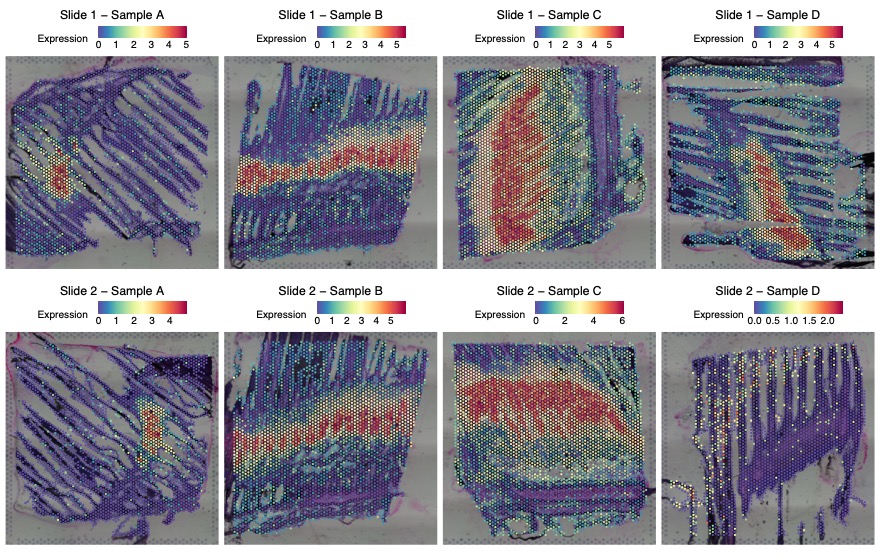

Figure S10.7 The spatial distribution of g3p (Glyceraldehyde-3-phosphate dehydrogenase; Ensembl ID: ENSSSAG00000067521) expression in eight tissue sections (Slide 1: Samples A to D; Slide 2: Samples A to D). Each panel corresponds to one tissue section, with expression values represented by colour intensity. The colour intensity reflects normalised expression, using SCTransform, where higher values indicate greater relative expression following sequencing depth correction.

***g3p*(ENSSSAG00000073639)**

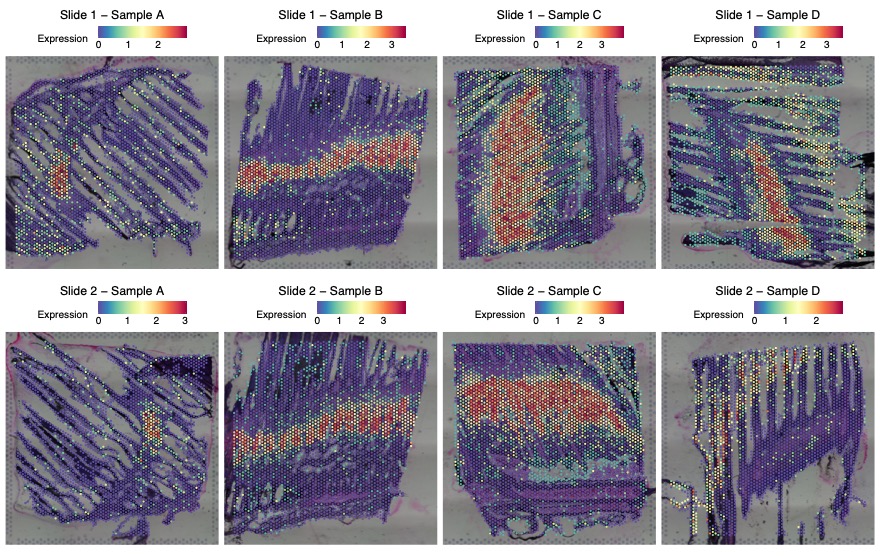

Figure S10.8 The spatial distribution of *g3p* (Glyceraldehyde-3-phosphate dehydrogenase; Ensembl ID: ENSSSAG00000073639) expression in eight tissue sections (Slide 1: Samples A to D; Slide 2: Samples A to D). Each panel corresponds to one tissue section, with expression values represented by colour intensity. The colour intensity reflects normalised expression, using SCTransform, where higher values indicate greater relative expression following sequencing depth correction.

***cx6a1* (ENSSSAG00000041369)**

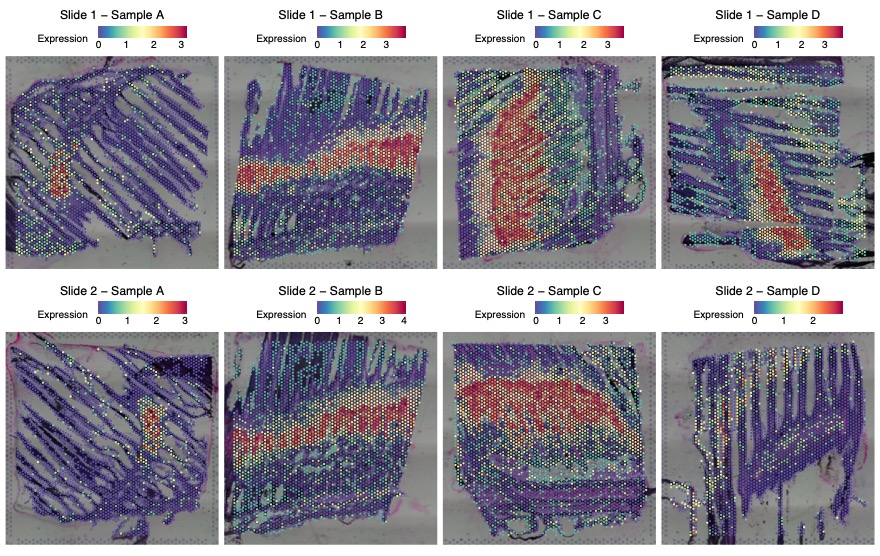

Figure S10.9 The spatial distribution of *cx6a1* (cytochrome c oxidase subunit 6A, mitochondrial; Ensembl ID: ENSSSAG00000041369) expression in eight tissue sections (Slide 1: Samples A to D; Slide 2: Samples A to D). Each panel corresponds to one tissue section, with expression values represented by colour intensity. The colour intensity reflects normalised expression, using SCTransform, where higher values indicate greater relative expression following sequencing depth correction.

***cox4i2* (ENSSSAG00000063792)**

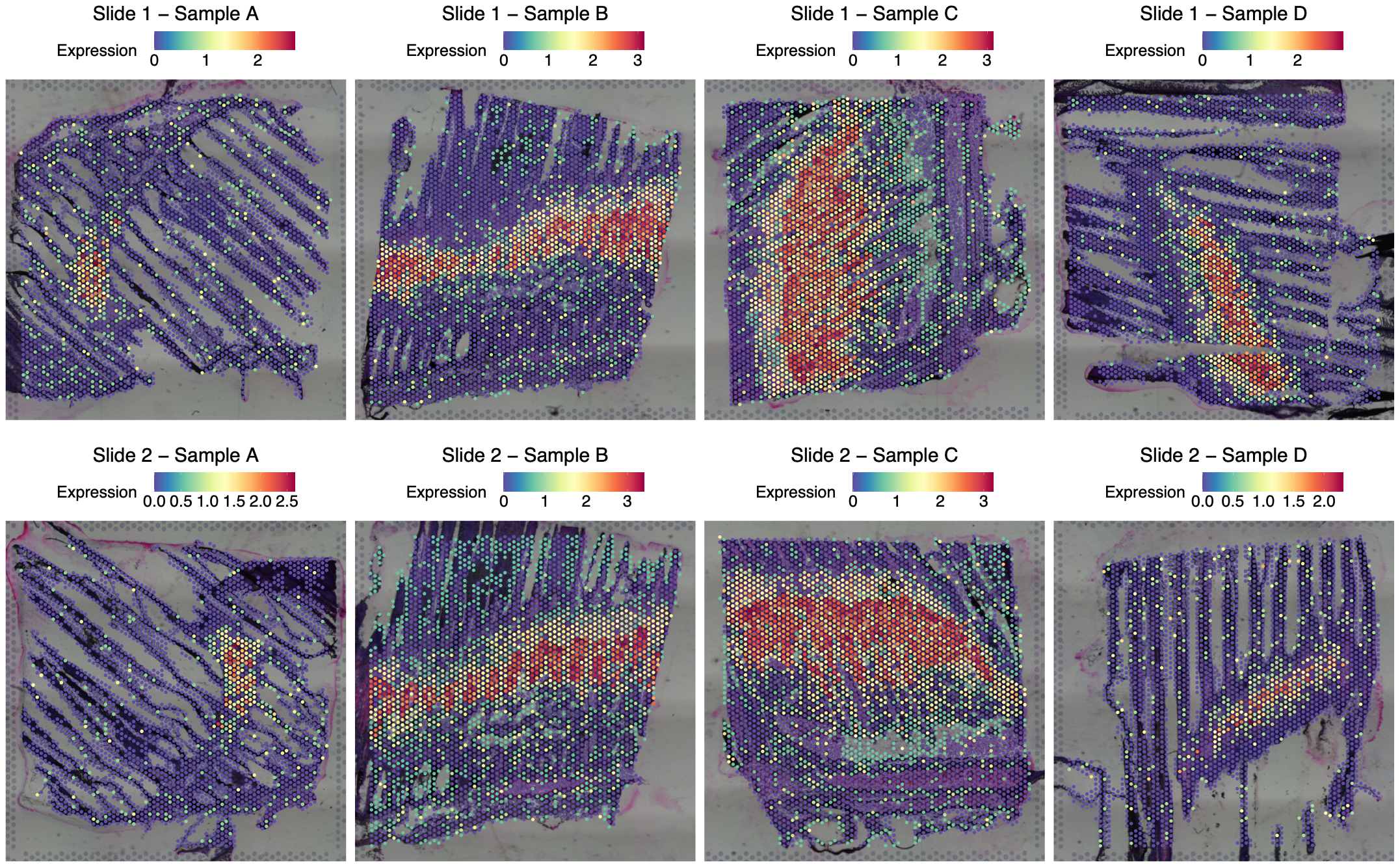

Figure S10.10 The spatial distribution of *cox4i2* (cytochrome c oxidase subunit 4I2; Ensembl ID: ENSSSAG00000063792) expression in eight tissue sections (Slide 1: Samples A to D; Slide 2: Samples A to D). Each panel corresponds to one tissue section, with expression values represented by colour intensity. The colour intensity reflects normalised expression, using SCTransform, where higher values indicate greater relative expression following sequencing depth correction.

Collectively, the gene expression profile of Cluster 3 suggests it represents a high proportion muscular component of the gill, likely composed of skeletal muscle. Enriched expression of contractile and aerobic metabolism genes may reflect a role in powering voluntary or reflexive gill movements essential for ventilation and respiration.
