## Supplementary Note 9 for "Spatial Transcriptomics Reveals Region-Specific Host Responses in Amoebic Gill Disease of Atlantic Salmon"

**Supplementary Note 9.** **Spatial Transcriptomic Characterisation of Gill Arch and Associated Smooth Muscle**

**9.1. Transcriptional Signatures of Connective Tissue Structure and Extracellular Matrix Remodelling in 3.2.2	Gill arch clusters**

The extracellular matrix (ECM) of the gill arch was marked by high expression of genes **ENSSSAG00000005527** (collagen alpha-1(I) chain; Entrez Gene ID: LOC106610502; UniProt: A0A1S3SM10; Cluster 7; Figure S9.1.1), which forms the major fibrillar scaffold of connective tissues; ***lum*** (lumican; Ensembl ID: ENSSSAG00000020979; Cluster 7; Figure S9.1.2), a small leucine-rich proteoglycan that regulates collagen fibrillogenesis; and **ENSSSAG00000085249** (Tropoelastin; InterPro: IPR003979; Cluster 7; Figure S9.1.3), the soluble precursor of elastin that contributes elasticity to vascular and connective tissues. ECM remodelling and regulatory functions were indicated by the expression of ***thbs4a*** (thrombospondin 4a; Ensembl ID: ENSSSAG00000071963; UniProt: A0A1S3PF42; Cluster 7; Figure S9.1.4), a matricellular glycoprotein implicated in cell matrix interactions, and ***timp3*** (TIMP metallopeptidase inhibitor 3; Ensembl ID: ENSSSAG00000073256; Entrez Gene ID: LOC106609153; Cluster 7; Figure S9.1.5), which inhibits matrix metalloproteinases to preserve ECM integrity. Collectively, these ECM associated genes within Metacluster 2 are likely contribute to both the structural integrity and the dynamic remodelling capacity of the connective tissues of the gill arch.

**ENSSSAG00000005527**

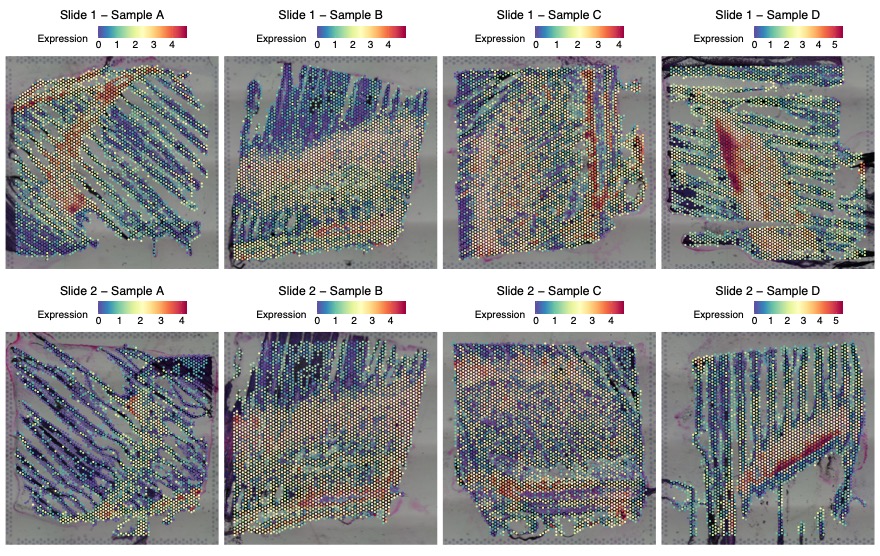

Figure S9.1.1 The spatial distribution of ENSSSAG00000005527 (collagen alpha-1(I) chain; Entrez Gene ID: LOC106610502; UniProt: A0A1S3SM10) expression in eight tissue sections (Slide 1: Samples A to D; Slide 2: Samples A to D). Each panel corresponds to one tissue section, with expression values represented by colour intensity. The colour intensity reflects normalised expression, using SCTransform, where higher values indicate greater relative expression following sequencing depth correction.

***lum* (ENSSSAG00000020979)**

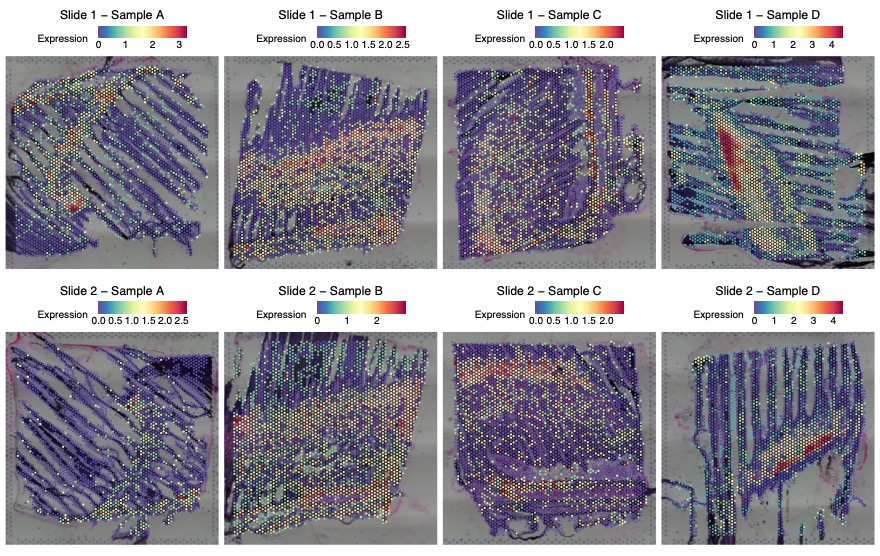

Figure S9.1.2 The spatial distribution of *lum* (lumican; Ensembl ID: ENSSSAG00000020979) expression in eight tissue sections (Slide 1: Samples A to D; Slide 2: Samples A to D). Each panel corresponds to one tissue section, with expression values represented by colour intensity. The colour intensity reflects normalised expression, using SCTransform, where higher values indicate greater relative expression following sequencing depth correction.

**ENSSSAG00000085249**

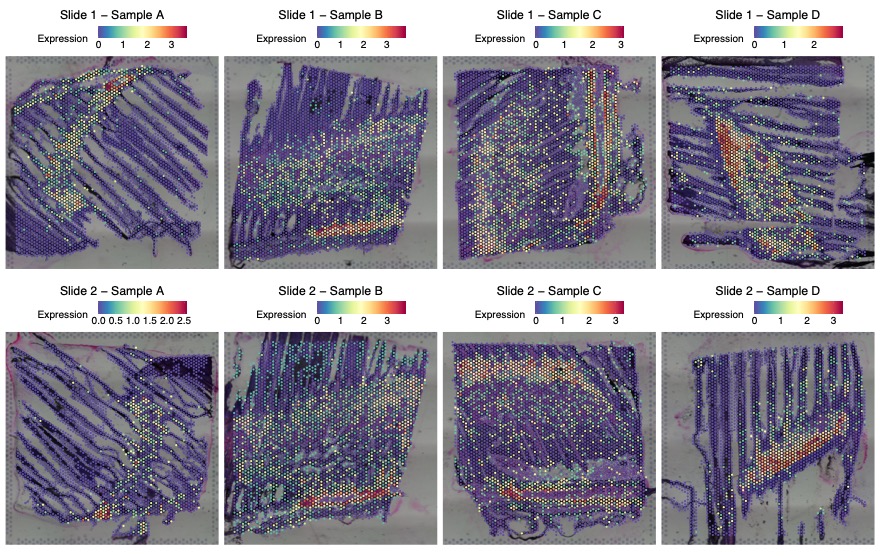

Figure S9.1.3 The spatial distribution of ENSSSAG00000085249 (Tropoelastin; InterPro: IPR003979) expression in eight tissue sections (Slide 1: Samples A to D; Slide 2: Samples A to D). Each panel corresponds to one tissue section, with expression values represented by colour intensity. The colour intensity reflects normalised expression, using SCTransform, where higher values indicate greater relative expression following sequencing depth correction.

***thbs4a* (ENSSSAG00000071963)**

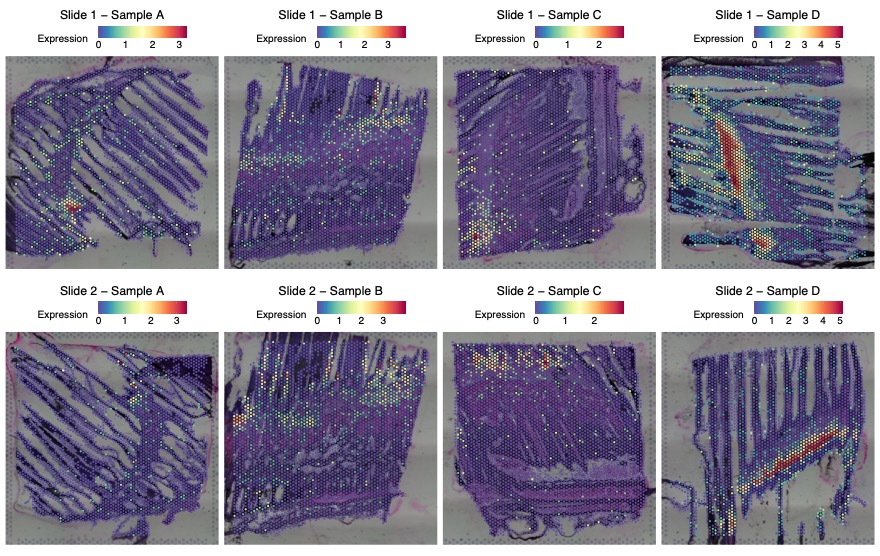

Figure S9.1.4 The spatial distribution of *thbs4a* (thrombospondin 4a; Ensembl ID: ENSSSAG00000071963; UniProt: A0A1S3PF42) expression in eight tissue sections (Slide 1: Samples A to D; Slide 2: Samples A to D). Each panel corresponds to one tissue section, with expression values represented by colour intensity. The colour intensity reflects normalised expression, using SCTransform, where higher values indicate greater relative expression following sequencing depth correction.

***timp3* (ENSSSAG00000073256)**

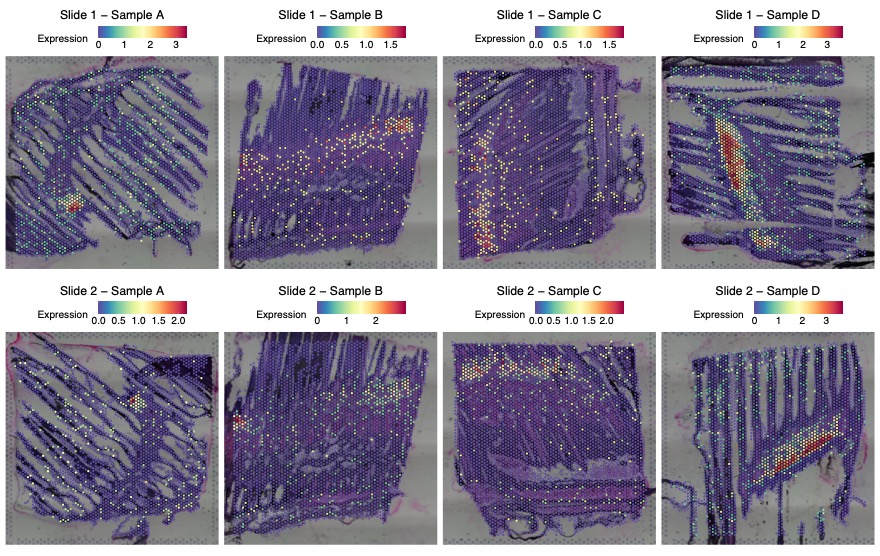

Figure S9.1.5 The spatial distribution of *timp3* (TIMP metallopeptidase inhibitor 3; Ensembl ID: ENSSSAG00000073256; Entrez Gene ID: LOC106609153) expression in eight tissue sections (Slide 1: Samples A to D; Slide 2: Samples A to D). Each panel corresponds to one tissue section, with expression values represented by colour intensity. The colour intensity reflects normalised expression, using SCTransform, where higher values indicate greater relative expression following sequencing depth correction.

**9.2 Smooth Muscle Contractility Signatures in 3.2.2 Gill arch clusters**

The gene expression profile within Cluster 8 is dominated by elements of smooth muscle contractility, with a significant proportion of its top marker genes aligning closely with smooth muscle structures of the gill arch. Key markers included a smooth muscle-associated cytoskeletal and contractile proteins: ***cald1b*** (caldesmon 1b, non-muscle caldesmon-like isoform X1; Ensembl ID: ENSSSAG00000100045; Entrez Gene ID: LOC106573078; UniProt: A0A1S3MJG3; Cluster 8; Figure S9.2.1), ***acta2*** (actin alpha 2, smooth muscle; Ensembl ID: ENSSSAG00000044179; UniProt: A0A1S3Q4A0; Cluster 8; Figure S9.2.2), **ENSSSAG00000047730** (desmin; Entrez Gene ID: LOC106574241; UniProt: A0A1S3MRL7; Cluster 8; Figure S9.2.3), ***lmod1b*** (leiomodin 1b (smooth muscle); Ensembl ID: ENSSSAG00000064672; Entrez Gene ID: LOC106566054; UniProt: A0A1S3LEU6; Cluster 8; Figure S9.2.4), ***smtna*** (smoothelin a; Ensembl ID: ENSSSAG00000050262; Entrez Gene ID: LOC106563188; UniProt: A0A1S3KZ62; Cluster 8; Figure S9.2.5), ***cnn1b*** (calponin 1, basic, smooth muscle, b; Ensembl ID: ENSSSAG00000122078; Cluster 8; Figure S9.2.6), and ***actn1*** (actinin, alpha 1; Ensembl ID: ENSSSAG00000114975; Cluster 8; Figure S9.2.7). Collectively, these genes form the core machinery for contraction, structural integrity, and dynamic control essential for efficient gill respiration and other physiological functions, highlighting the role of smooth muscle in regulating involuntary actions such as blood flow within vessels and fine-tuning movements that support skeletal muscle fibers.

***cald1b* (ENSSSAG00000100045)**

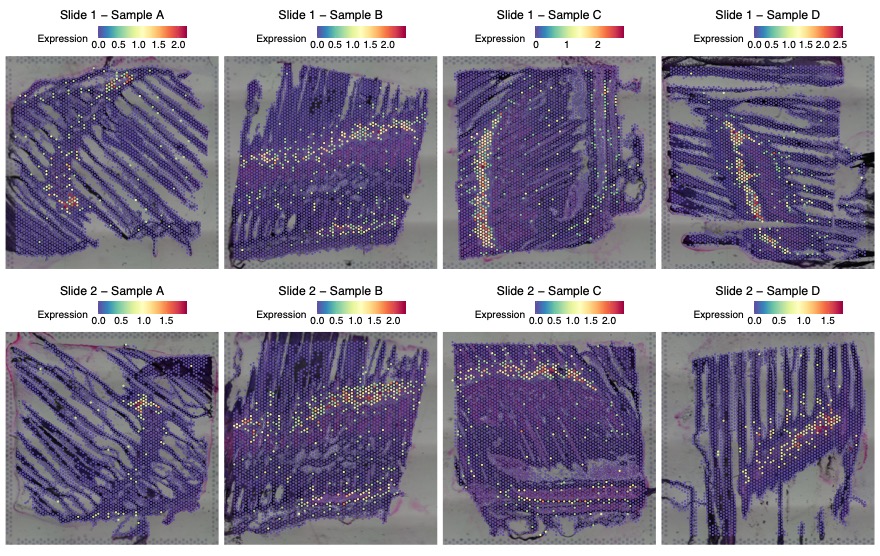

Figure S9.2.1 The spatial distribution of *cald1b* (caldesmon 1b, non-muscle caldesmon-like isoform X1; Ensembl ID: ENSSSAG00000100045; Entrez Gene ID: LOC106573078; UniProt: A0A1S3MJG3) expression in eight tissue sections (Slide 1: Samples A to D; Slide 2: Samples A to D). Each panel corresponds to one tissue section, with expression values represented by colour intensity. The colour intensity reflects normalised expression, using SCTransform, where higher values indicate greater relative expression following sequencing depth correction.

***acta2* (ENSSSAG00000044179)**

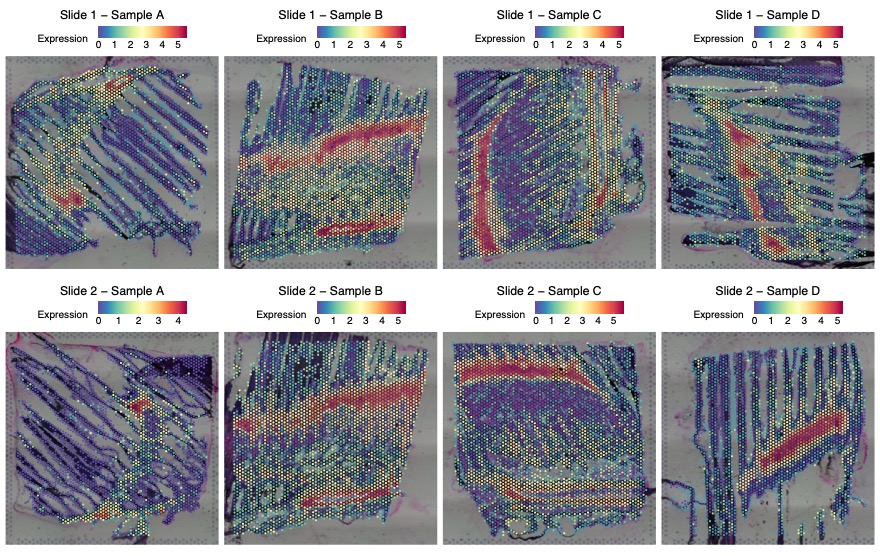

Figure S9.2.2 The spatial distribution of *acta2* (actin alpha 2, smooth muscle; Ensembl ID: ENSSSAG00000044179; UniProt: A0A1S3Q4A0) expression in eight tissue sections (Slide 1: Samples A to D; Slide 2: Samples A to D). Each panel corresponds to one tissue section, with expression values represented by colour intensity. The colour intensity reflects normalised expression, using SCTransform, where higher values indicate greater relative expression following sequencing depth correction.

**ENSSSAG00000047730**

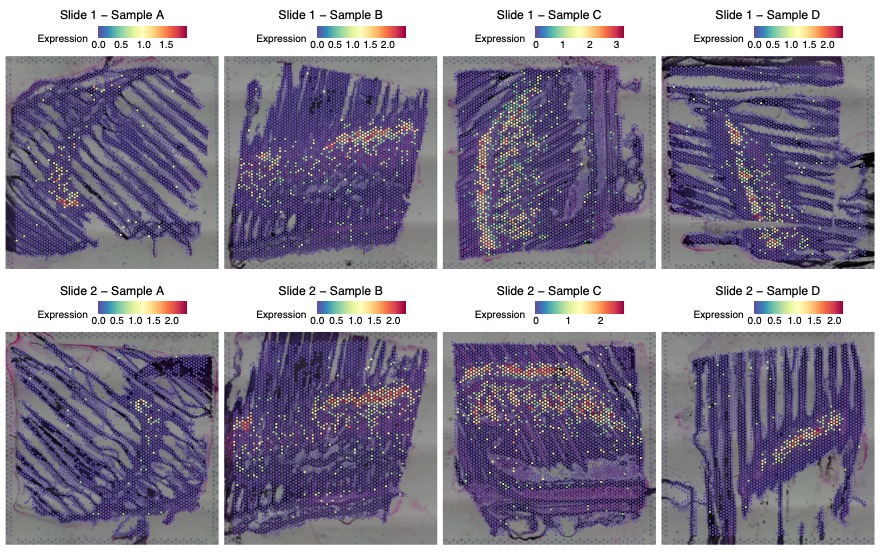

Figure S9.2.3 The spatial distribution of ENSSSAG00000047730 (desmin; Entrez Gene ID: LOC106574241; UniProt: A0A1S3MRL7) expression in eight tissue sections (Slide 1: Samples A to D; Slide 2: Samples A to D). Each panel corresponds to one tissue section, with expression values represented by colour intensity. The colour intensity reflects normalised expression, using SCTransform, where higher values indicate greater relative expression following sequencing depth correction.

***lmod1b* (ENSSSAG00000064672)**

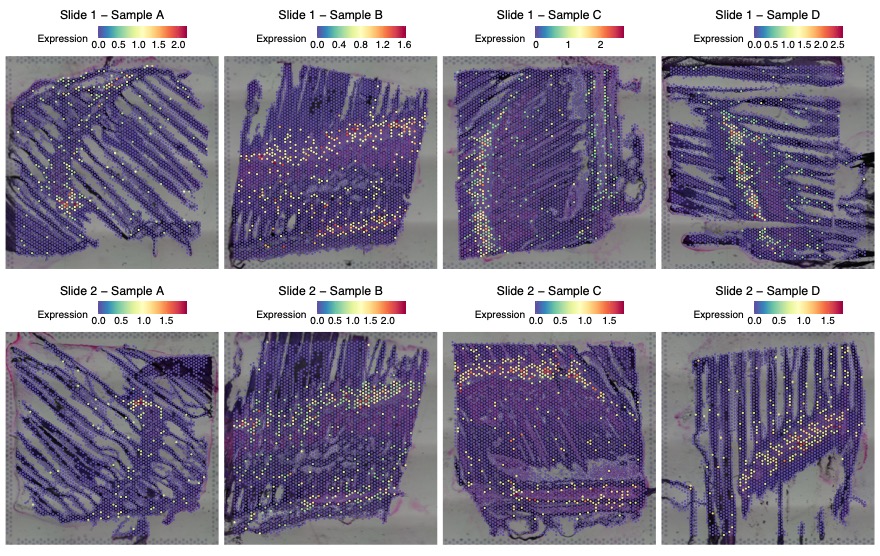

Figure S9.2.4 The spatial distribution of *lmod1b* (leiomodin 1b (smooth muscle); Ensembl ID: ENSSSAG00000064672; Entrez Gene ID: LOC106566054; UniProt: A0A1S3LEU6) expression in eight tissue sections (Slide 1: Samples A to D; Slide 2: Samples A to D). Each panel corresponds to one tissue section, with expression values represented by colour intensity. The colour intensity reflects normalised expression, using SCTransform, where higher values indicate greater relative expression following sequencing depth correction.

***smtna* (ENSSSAG00000050262)**

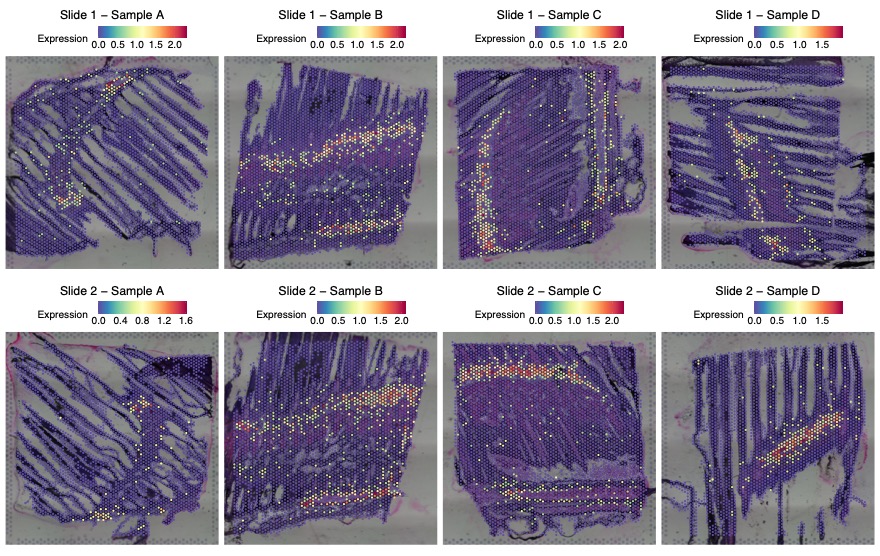

Figure S9.2.5 The spatial distribution smtna (smoothelin a; Ensembl ID: ENSSSAG00000050262; Entrez Gene ID: LOC106563188; UniProt: A0A1S3KZ62) expression in eight tissue sections (Slide 1: Samples A to D; Slide 2: Samples A to D). Each panel corresponds to one tissue section, with expression values represented by colour intensity. The colour intensity reflects normalised expression, using SCTransform, where higher values indicate greater relative expression following sequencing depth correction.

***cnn1b* (ENSSSAG00000122078)**

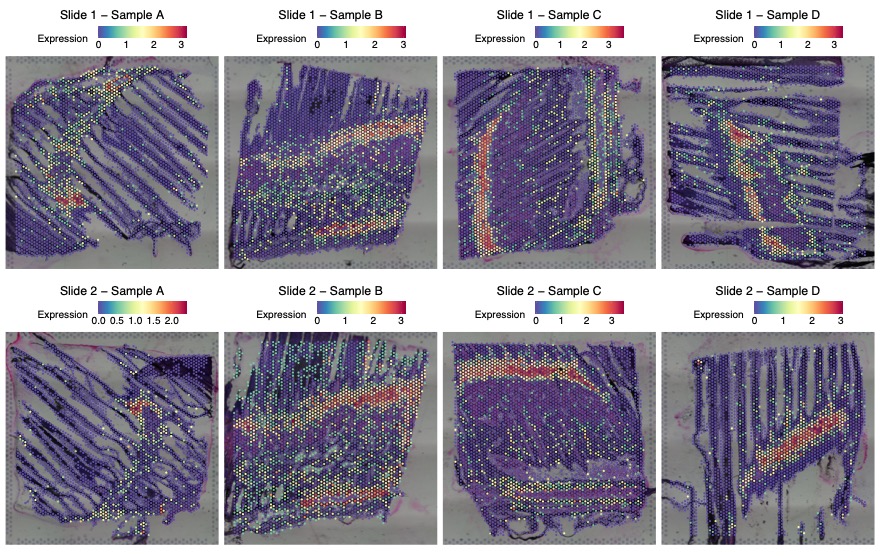

Figure S9.2.6 The spatial distribution of *cnn1b* (calponin 1, basic, smooth muscle, b; Ensembl ID: ENSSSAG00000122078) expression in eight tissue sections (Slide 1: Samples A to D; Slide 2: Samples A to D). Each panel corresponds to one tissue section, with expression values represented by colour intensity. The colour intensity reflects normalised expression, using SCTransform, where higher values indicate greater relative expression following sequencing depth correction.

***actn1*(ENSSSAG00000114975)**

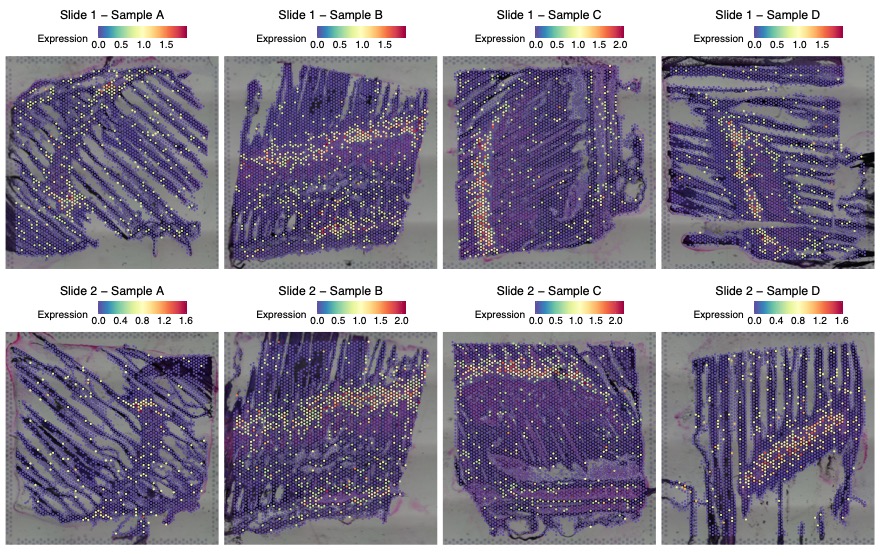

Figure S9.2.7 The spatial distribution of *actn1* (actinin, alpha 1; Ensembl ID: ENSSSAG00000114975) expression in eight tissue sections (Slide 1: Samples A to D; Slide 2: Samples A to D). Each panel corresponds to one tissue section, with expression values represented by colour intensity. The colour intensity reflects normalised expression, using SCTransform, where higher values indicate greater relative expression following sequencing depth correction.

**9.3. Spatial Mapping of Vascular and Lymphatic Function Genes in in 3.2.2 Gill arch clusters**

Gene expression profile indicative of crucial vascular and lymphatic vessel function has been largely exhibited as the top marker genes in cluster 9 with its spots notably scattered within muscular regions (Figure 14).Elevated expression of ***xlkd1*** (Lymphatic vessel endothelial hyaluronic acid receptor 1; Ensembl ID: ENSSSAG00000072759; Cluster 9; Figure S9.3.1) and **ENSSSAG00000098957** (Lymphatic vessel endothelial hyaluronic acid receptor 1-like; Entrez Gene ID: LOC106562111; UniProt: A0A1S3KV59; Cluster 9; Figure S9.3.2), genes commonly associated with lymphatic endothelial cells, may indicate potential involvement in fluid regulation and immune cell trafficking. ***gp182*** (G-protein coupled receptor 182; Ensembl ID: ENSSSAG00000088534 (Figure S9.3.3), ENSSSAG00000077688 (Figure S9.3.4); Cluster 9), linked to angiogenesis and vascular development, was also expressed at high levels, pointing to possible roles in vascular activity. Collectively, the top marker genes observed in Cluster 9 suggest a potential involvement in vascular and lymphatic functions, which may contribute to fluid regulation, immune responses, and tissue remodelling.

***xlkd1* (ENSSSAG00000072759)**

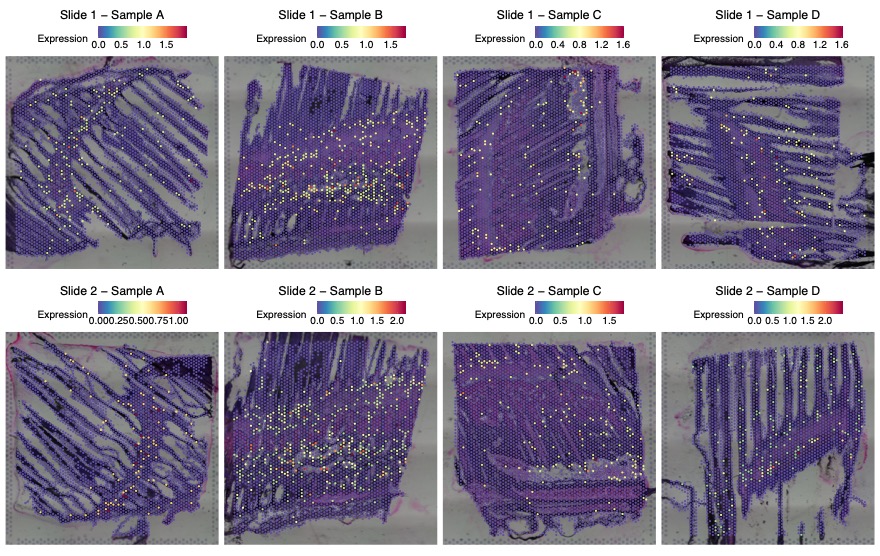

Figure S9.3.1 The spatial distribution of *xlkd1* (Lymphatic vessel endothelial hyaluronic acid receptor 1; Ensembl ID: ENSSSAG00000072759) expression in eight tissue sections (Slide 1: Samples A to D; Slide 2: Samples A to D). Each panel corresponds to one tissue section, with expression values represented by colour intensity. The colour intensity reflects normalised expression, using SCTransform, where higher values indicate greater relative expression following sequencing depth correction.

**ENSSSAG00000098957**

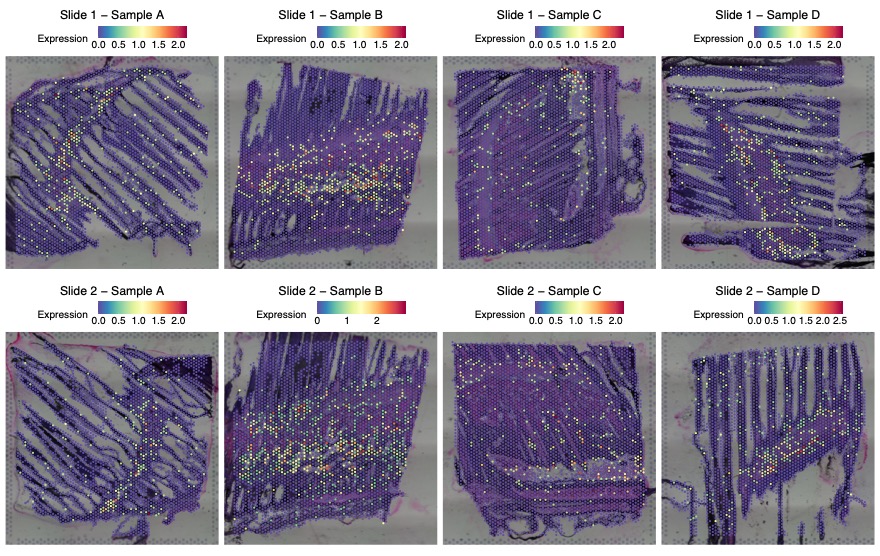

Figure S9.3.2 The spatial distribution of ENSSSAG00000098957 (Lymphatic vessel endothelial hyaluronic acid receptor 1-like; Entrez Gene ID: LOC106562111; UniProt: A0A1S3KV59) expression in eight tissue sections (Slide 1: Samples A to D; Slide 2: Samples A to D). Each panel corresponds to one tissue section, with expression values represented by colour intensity. The colour intensity reflects normalised expression, using SCTransform, where higher values indicate greater relative expression following sequencing depth correction.

***gp182* (ENSSSAG00000088534)**

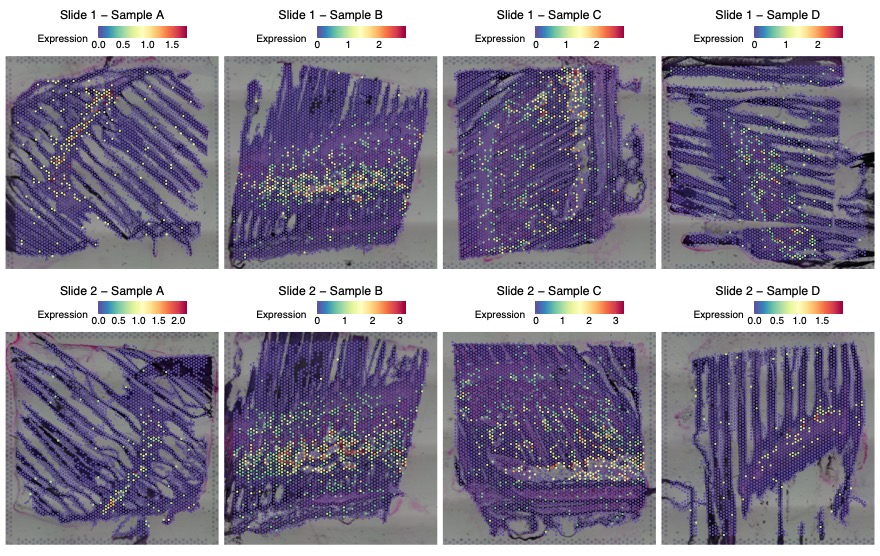

Figure S9.3.3 The spatial distribution of *gp182* (G-protein coupled receptor 182; Ensembl ID: ENSSSAG00000088534) expression in eight tissue sections (Slide 1: Samples A to D; Slide 2: Samples A to D). Each panel corresponds to one tissue section, with expression values represented by colour intensity. The colour intensity reflects normalised expression, using SCTransform, where higher values indicate greater relative expression following sequencing depth correction.

***gp182* (ENSSSAG00000077688)**

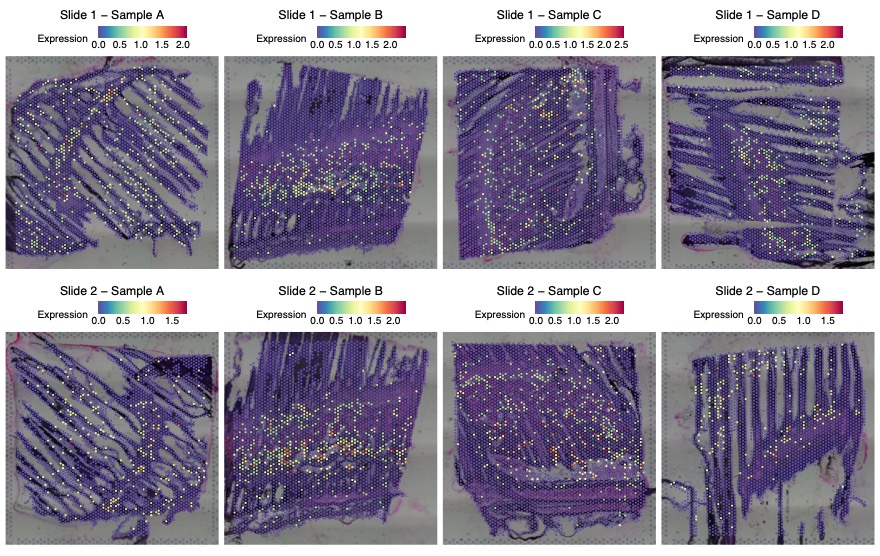

Figure S9.3.4 The spatial distribution of *gp182* (G-protein coupled receptor 182; Ensembl ID: ENSSSAG00000077688) expression in eight tissue sections (Slide 1: Samples A to D; Slide 2: Samples A to D). Each panel corresponds to one tissue section, with expression values represented by colour intensity. The colour intensity reflects normalised expression, using SCTransform, where higher values indicate greater relative expression following sequencing depth correction.
