## Supplementary Note 8 for "Spatial Transcriptomics Reveals Region-Specific Host Responses in Amoebic Gill Disease of Atlantic Salmon"

**Supplementary Note 8.** Spatial Transcriptomic Landscape of Gill Filaments and Lamellae

**8.1. Transcriptional Signatures of Gill Epithelial Integrity and Extracellular Matrix Remodelling in 3.2.1	Filament and lamellae clusters**

An epithelial framework is indicated by elevated expression of **ENSSSAG00000101879** (keratin, type I cytoskeletal 13-like; Entrez Gene ID: LOC106564957; UniProt: A0A1S3L7Q2; Cluster 2; Figure S8.1.1), ***krt15*** (keratin 15; Ensembl ID: ENSSSAG00000045452; Cluster 10; Figure S8.1.2), and **ENSSSAG00000048372** (keratin, type I cytoskeletal 13; Entrez Gene ID: LOC106587847; UniProt: A0A1S3PWC1; Cluster 13; Figure S8.1.3), all of which are essential intermediate filaments likely to be important for mechanical resilience at these exposed surfaces.

**ENSSSAG00000101879**

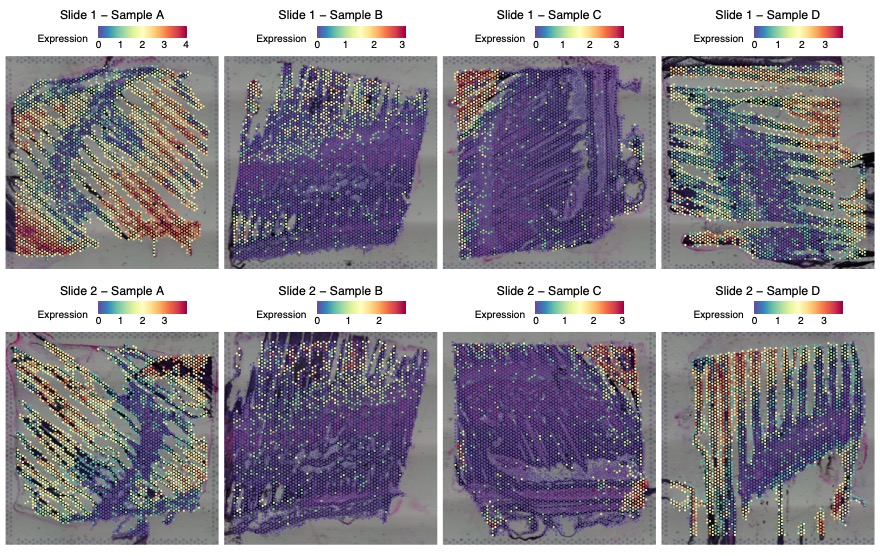

Figure S8.1.1 The spatial distribution of ENSSSAG00000101879 (keratin, type I cytoskeletal 13-like; Entrez Gene ID: LOC106564957; UniProt: A0A1S3L7Q2) expression in eight tissue sections (Slide 1: Samples A to D; Slide 2: Samples A to D). Each panel corresponds to one tissue section, with expression values represented by colour intensity. The colour intensity reflects normalised expression, using SCTransform, where higher values indicate greater relative expression following sequencing depth correction.

***krt15* (ENSSSAG00000045452)**

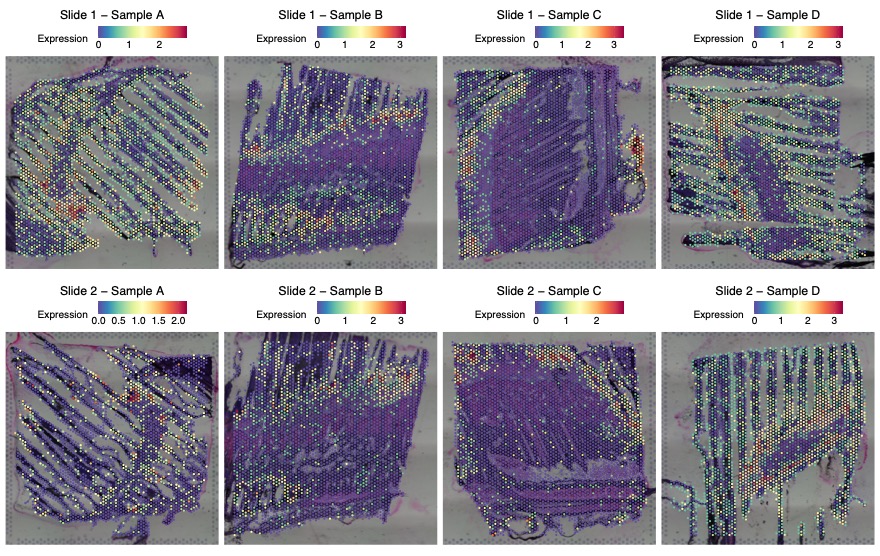

Figure S8.1.2 The spatial distribution of *krt15* (keratin 15; Ensembl ID: ENSSSAG00000045452) expression in eight tissue sections (Slide 1: Samples A to D; Slide 2: Samples A to D). Each panel corresponds to one tissue section, with expression values represented by colour intensity. The colour intensity reflects normalised expression, using SCTransform, where higher values indicate greater relative expression following sequencing depth correction.

**ENSSSAG00000048372**

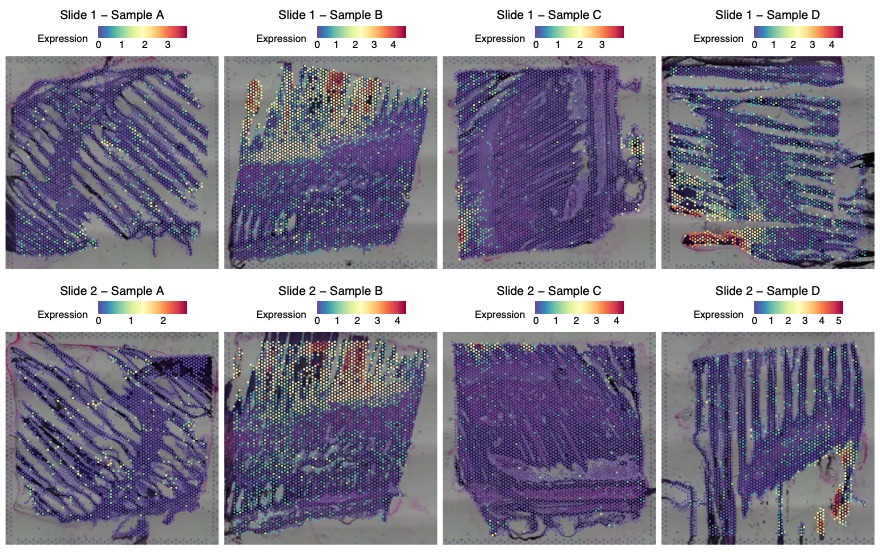

Figure S8.1.3 The spatial distribution of ENSSSAG00000048372 (keratin, type I cytoskeletal 13; Entrez Gene ID: LOC106587847; UniProt: A0A1S3PWC1) expression in eight tissue sections (Slide 1: Samples A to D; Slide 2: Samples A to D). Each panel corresponds to one tissue section, with expression values represented by colour intensity. The colour intensity reflects normalised expression, using SCTransform, where higher values indicate greater relative expression following sequencing depth correction.

Active extracellular matrix (ECM) remodelling is a dynamic process in which cells modify the surrounding ECM by degrading, synthesising, and reorganising its components. This coordinated activity is crucial for proper gill function in fish, supporting development, wound healing, and responses to disease. Active ECM remodelling in Cluster 11 is supported by the expression of several genes. These included ***col10a1b*** (collagen, type X, alpha 1b; Ensembl ID: ENSSSAG00000114656; Cluster 11; Figure S8.1.4), a specialised collagen involved in structural scaffold formation, and, **ENSSSAG00000090328** (collagen alpha-2(VI) chain; Entrez Gene ID: LOC106582581; Cluster 11; Figure S8.1.5), which contributes to ECM elasticity and cellular organisation through microfilament network formation. Furthermore, **ENSSSAG00000079745** (Metalloproteases catalytic domain; InterPro: IPR006026; Cluster 11; Figure S8.1.6) was expressed, indicating controlled ECM degradation as part of the remodelling process. Among the important ECM-associated components, ***spp1*** (Osteopontin; ENSSSAG00000100947; Cluster 11; Figure S8.1.7) was repeatedly identified across different clusters, serving as a matricellular protein mediating cell-matrix interactions and tissue remodelling signals. Collectively, these genes define a spatially organised transcriptional domain likely involved in maintaining the structural integrity of the gill filament and lamellar epithelium, while supporting key physiological functions including osmoregulation and immune defence.

***col10a1b* (ENSSSAG00000114656)**

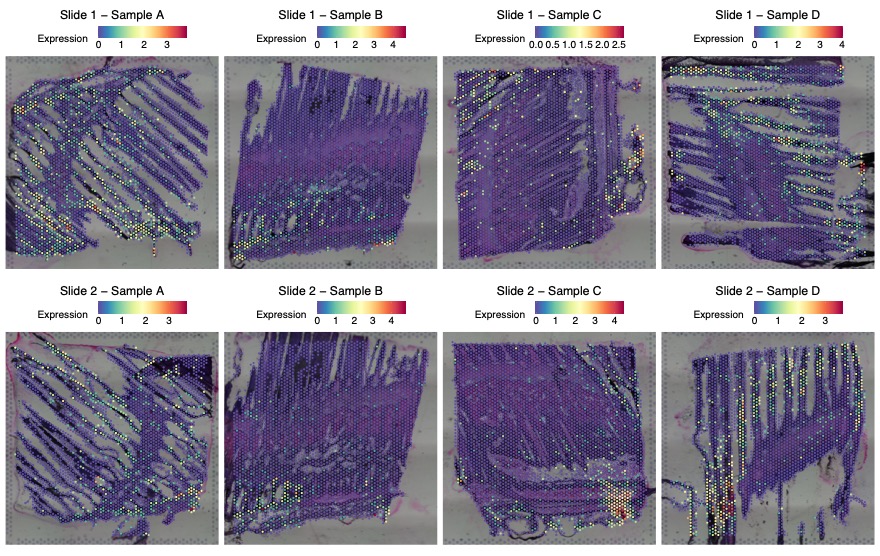

Figure S8.1.4 The spatial distribution of *col10a1b* (collagen, type X, alpha 1b; Ensembl ID: ENSSSAG00000114656) expression in eight tissue sections (Slide 1: Samples A to D; Slide 2: Samples A to D). Each panel corresponds to one tissue section, with expression values represented by colour intensity. The colour intensity reflects normalised expression, using SCTransform, where higher values indicate greater relative expression following sequencing depth correction.

**ENSSSAG00000090328**

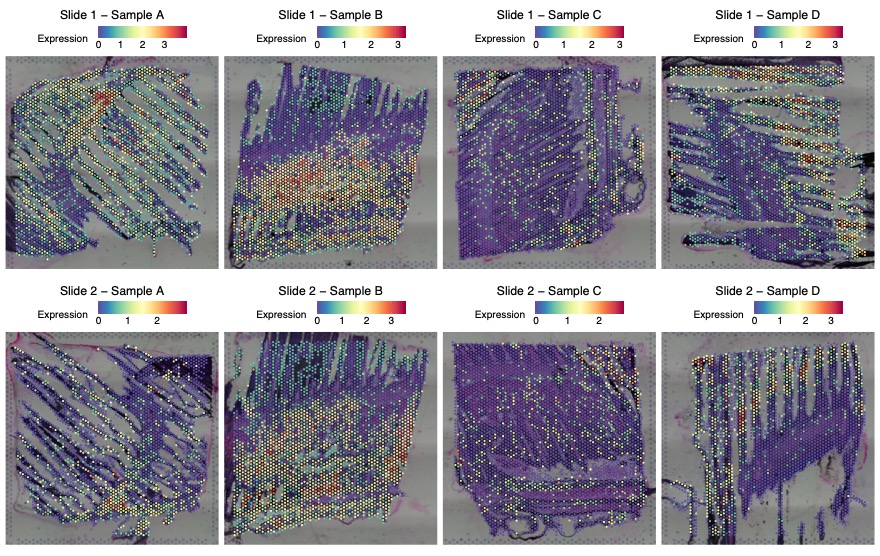

Figure S8.1.5 The spatial distribution of ENSSSAG00000090328 (collagen alpha-2(VI) chain; Entrez Gene ID: LOC106582581) expression in eight tissue sections (Slide 1: Samples A to D; Slide 2: Samples A to D). Each panel corresponds to one tissue section, with expression values represented by colour intensity. The colour intensity reflects normalised expression, using SCTransform, where higher values indicate greater relative expression following sequencing depth correction.

**ENSSSAG00000079745**

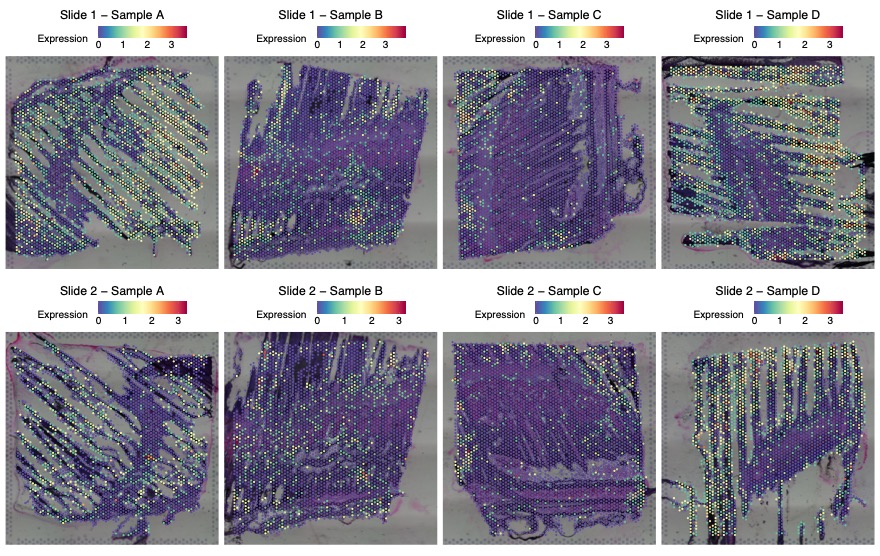

Figure S8.1.6 The spatial distribution of ENSSSAG00000079745 (Metalloproteases catalytic domain; InterPro: IPR006026) expression in eight tissue sections (Slide 1: Samples A to D; Slide 2: Samples A to D). Each panel corresponds to one tissue section, with expression values represented by colour intensity. The colour intensity reflects normalised expression, using SCTransform, where higher values indicate greater relative expression following sequencing depth correction.

***spp1* (ENSSSAG00000100947)**

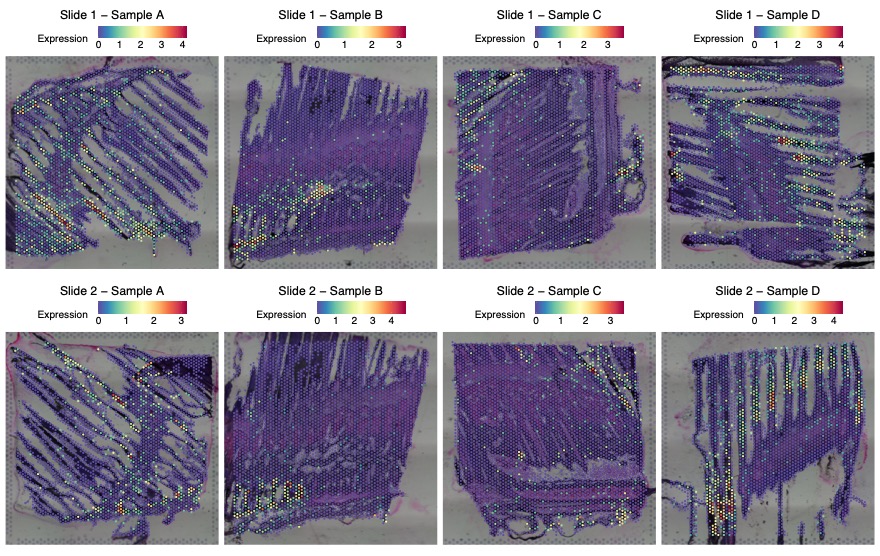

Figure S8.1.7 The spatial distribution of *spp1* (Osteopontin; ENSSSAG00000100947) expression in eight tissue sections (Slide 1: Samples A to D; Slide 2: Samples A to D). Each panel corresponds to one tissue section, with expression values represented by colour intensity. The colour intensity reflects normalised expression, using SCTransform, where higher values indicate greater relative expression following sequencing depth correction.

**8.2. Spatial Mapping of Ion Transporters and Channels Reveals Key Elements of Gill Osmoregulation**

Osmoregulation, the important process of maintaining internal salt and water balance, is a primary function of the fish gill, particularly within its filaments. In the spatial transcriptomics analysis, genes associated with osmoregulation, identified as both DEGs and SVGs, were prominently ranked among the top markers in both Cluster 12 and Cluster 2. These regions showed elevated expression of ATP-dependent ion transporters as well as ion channels that facilitate passive ion movement. Both are essential for maintaining ionic gradients across the gill epithelium and key elements of ionocytes (chloride cells).

Transporter-related genes included **ENSSSAG00000055927** (sodium/hydrogen exchanger 3; Entrez Gene ID: LOC106569766; UniProt: A0A1S3M1V3; Cluster 12; Figure S8.2.1Figure), which contributes to sodium uptake and acid-base balance, and **ENSSSAG00000088702** (P-type Cation transport ATPase; InterPro: IPR004014; Cluster 12; Figure S8.2.2) that establish the electrochemical gradients essential for active ion transport. In addition, **ENSSSAG00000104002** and **ENSSSAG00000056493** (Ion-transport regulator, FXYD family; InterPro: IPR000272; Cluster 12; Figure S8.2.3 & Figure S8.2.4**Error! Reference source not found.**), which modulate Na⁺/K⁺-ATPase activity, were also strongly expressed, supporting the dynamic regulation of transporter function.

**ENSSSAG00000055927**

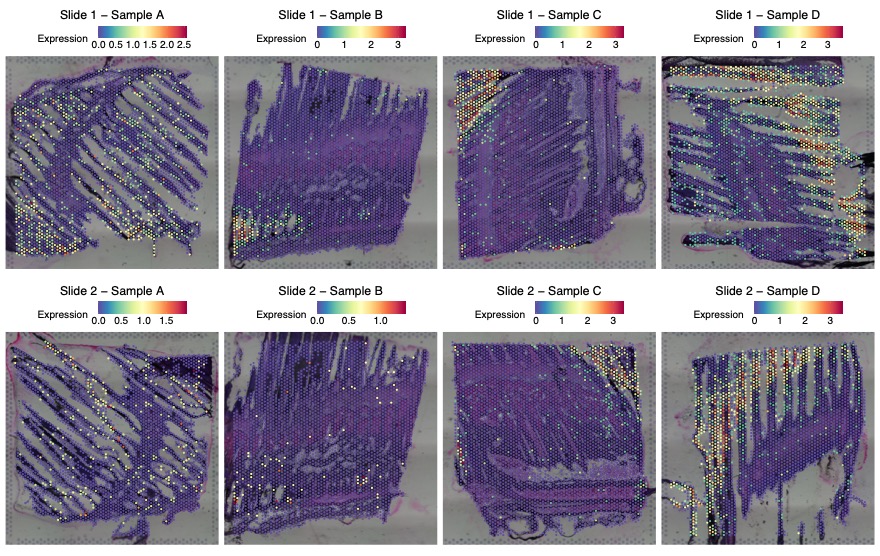

Figure S8.2.1: The spatial distribution of ENSSSAG00000055927 (sodium/hydrogen exchanger 3; Entrez Gene ID: LOC106569766; UniProt: A0A1S3M1V3) expression in eight tissue sections (Slide 1: Samples A to D; Slide 2: Samples A to D). Each panel corresponds to one tissue section, with expression values represented by colour intensity. The colour intensity reflects normalised expression, using SCTransform, where higher values indicate greater relative expression following sequencing depth correction.

**ENSSSAG00000088702**

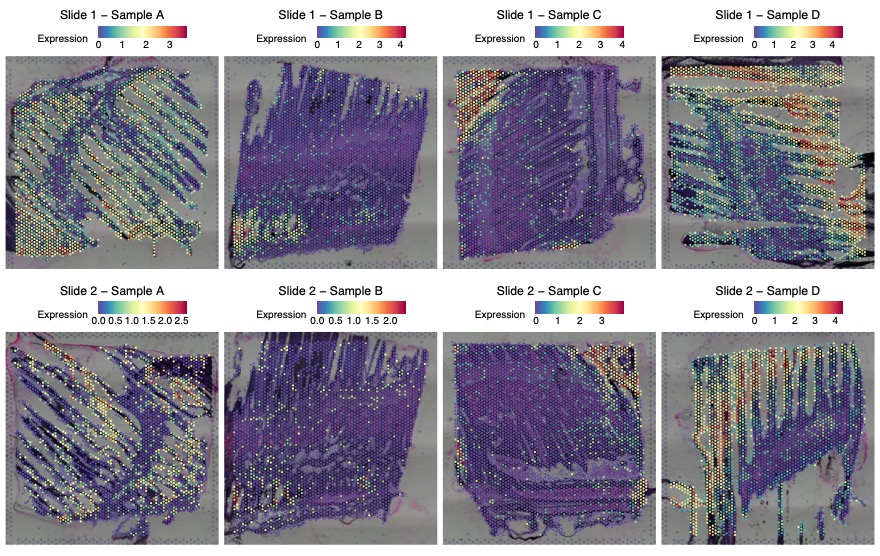

Figure S8.2.2 The spatial distribution of ENSSSAG00000088702 (P-type Cation transport ATPase; InterPro: IPR004014) expression in eight tissue sections (Slide 1: Samples A to D; Slide 2: Samples A to D). Each panel corresponds to one tissue section, with expression values represented by colour intensity. The colour intensity reflects normalised expression, using SCTransform, where higher values indicate greater relative expression following sequencing depth correction.

**ENSSSAG00000104002**

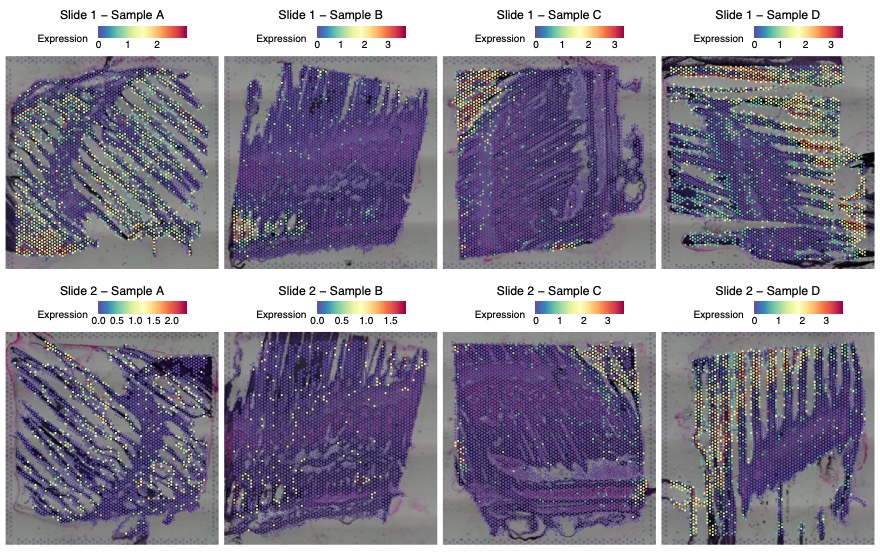

Figure S8.2.3 The spatial distribution of ENSSSAG00000104002 (Ion-transport regulator, FXYD family; InterPro: IPR000272) expression in eight tissue sections (Slide 1: Samples A to D; Slide 2: Samples A to D). Each panel corresponds to one tissue section, with expression values represented by colour intensity. The colour intensity reflects normalised expression, using SCTransform, where higher values indicate greater relative expression following sequencing depth correction.

**ENSSSAG00000056493**

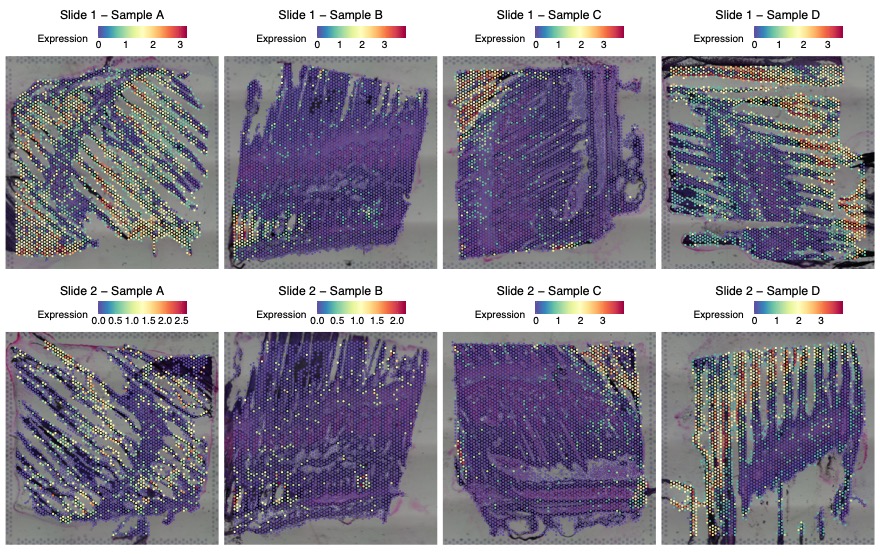

Figure S8.2.4 The spatial distribution of ENSSSAG00000056493 (Ion-transport regulator, FXYD family; InterPro: IPR000272) expression in eight tissue sections (Slide 1: Samples A to D; Slide 2: Samples A to D). Each panel corresponds to one tissue section, with expression values represented by colour intensity. The colour intensity reflects normalised expression, using SCTransform, where higher values indicate greater relative expression following sequencing depth correction.

Channel-related genes included ***clic1*** (chloride intracellular channel protein 1-like; Ensembl ID: ENSSSAG00000112534; Entrez Gene ID: LOC106588213; UniProt: A0A1S3PZ28; Cluster 2; Figure S8.2.5), which encodes chloride intracellular channel protein 1 involved in chloride transport. Additionally, transient receptor potential (TRP) channels, including ***trpv6*** (transient receptor potential cation channel, subfamily V, member 6; Ensembl ID: ENSSSAG00000059186; Entrez Gene ID: LOC106592665; Cluster 12; Figure S8.2.6) and **ENSSSAG00000091602** (Transient receptor potential cation channel M; InterPro: IPR050927; Cluster 12; Figure S8.2.7), were strongly expressed, contributing to calcium flux and osmosensory functions essential for gill ionocyte activity.

***clic1* (ENSSSAG00000112534)**

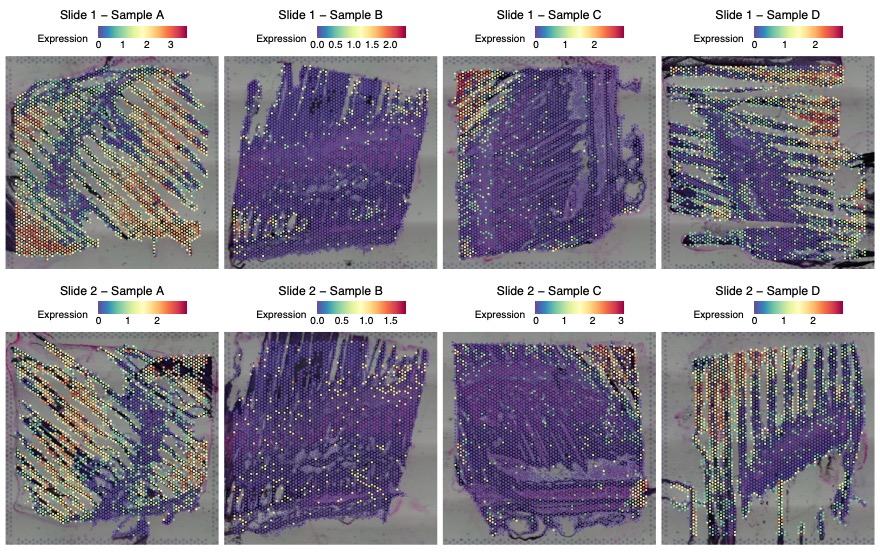
Figure S8.2.5 The spatial distribution of *clic1* (chloride intracellular channel protein 1-like; Ensembl ID: ENSSSAG00000112534; Entrez Gene ID: LOC106588213; UniProt: A0A1S3PZ28) expression in eight tissue sections (Slide 1: Samples A to D; Slide 2: Samples A to D). Each panel corresponds to one tissue section, with expression values represented by colour intensity. The colour intensity reflects normalised expression, using SCTransform, where higher values indicate greater relative expression following sequencing depth correction.

***trpv6* (ENSSSAG00000059186)**

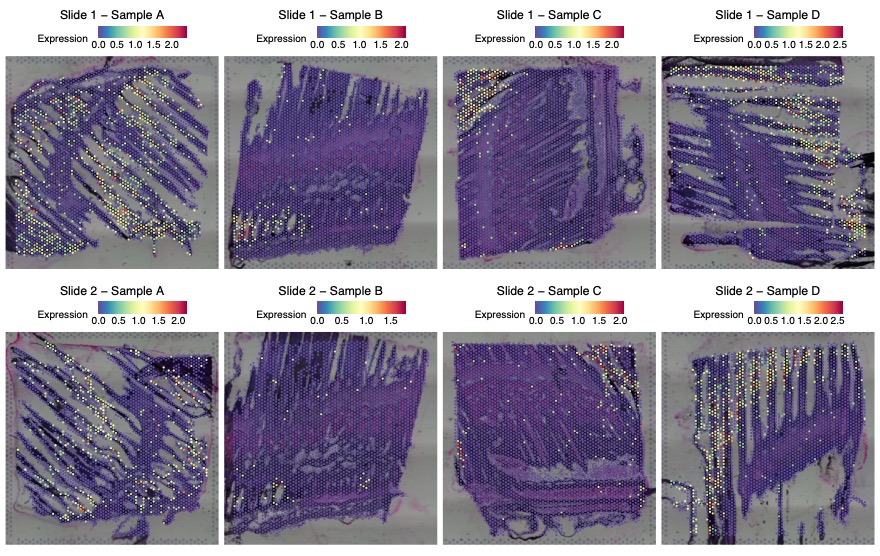

Figure S8.2.6 The spatial distribution of *trpv6* (transient receptor potential cation channel, subfamily V, member 6; Ensembl ID: ENSSSAG00000059186; Entrez Gene ID: LOC106592665) expression in eight tissue sections (Slide 1: Samples A to D; Slide 2: Samples A to D). Each panel corresponds to one tissue section, with expression values represented by colour intensity. The colour intensity reflects normalised expression, using SCTransform, where higher values indicate greater relative expression following sequencing depth correction.

**ENSSSAG00000091602**

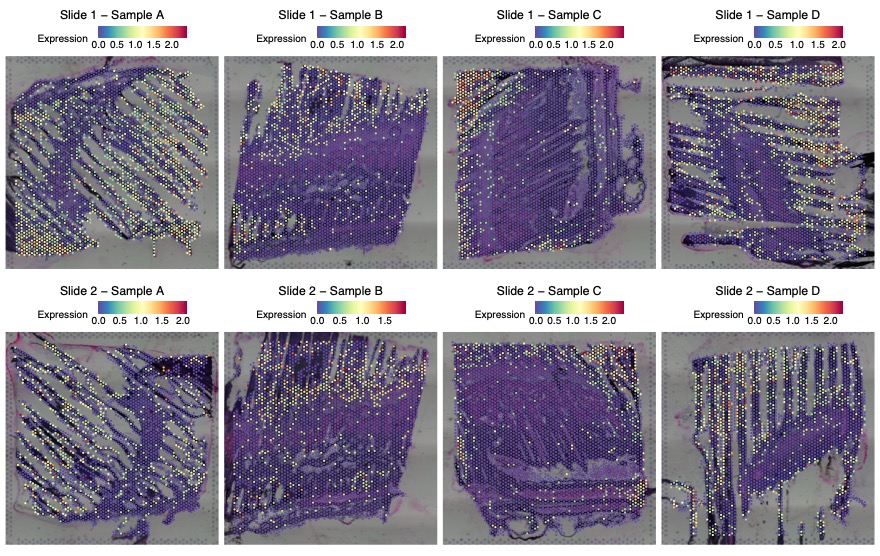

Figure S8.2.7 The spatial distribution of ENSSSAG00000091602 (Transient receptor potential cation channel M; InterPro: IPR050927) expression in eight tissue sections (Slide 1: Samples A to D; Slide 2: Samples A to D). Each panel corresponds to one tissue section, with expression values represented by colour intensity. The colour intensity reflects normalised expression, using SCTransform, where higher values indicate greater relative expression following sequencing depth correction.

Additional top marker genes in the cluster 12 included ***igfbp5a*** (insulin-like growth factor binding protein 5a; Ensembl ID: ENSSSAG00000088363, ENSSSAG00000039751; Entrez Gene ID: LOC106575566; Cluster 12; Figure S8.2.8 & Figure S8.2.9), a component of the Insulin-like Growth Factor (IGF) system. ***igfbp5a*** regulates IGF bioavailability and activity, suggesting a possible role in hormonal modulation of osmoregulatory functions within these cells.

***igfbp5a* (ENSSSAG00000088363)**

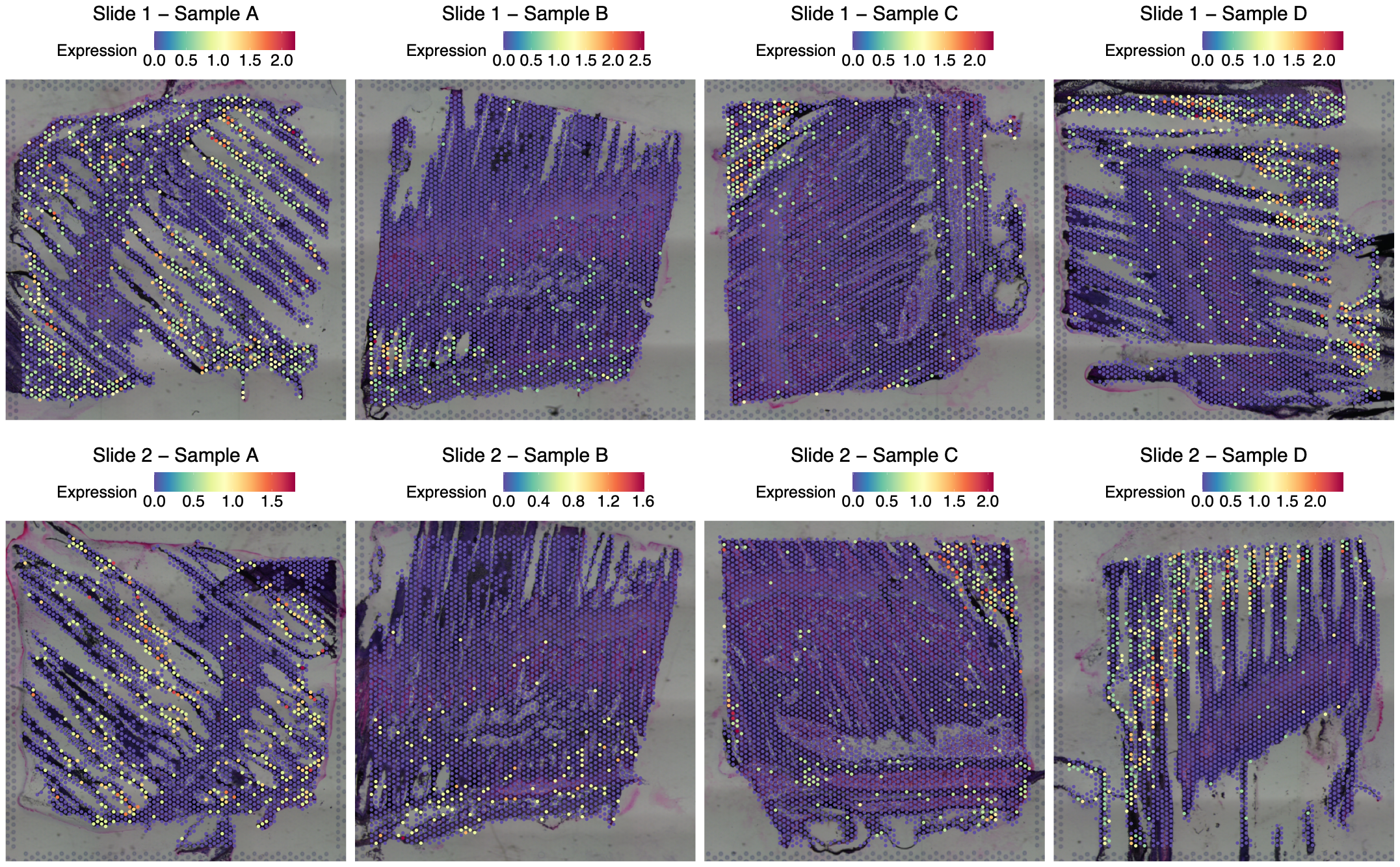

Figure S8.2.8 The spatial distribution of *igfbp5a* (insulin-like growth factor binding protein 5a; Ensembl ID: ENSSSAG00000088363; Entrez Gene ID: LOC106575566) expression in eight tissue sections (Slide 1: Samples A to D; Slide 2: Samples A to D). Each panel corresponds to one tissue section, with expression values represented by colour intensity. The colour intensity reflects normalised expression, using SCTransform, where higher values indicate greater relative expression following sequencing depth correction.

***igfbp5a* (ENSSSAG00000039751)**

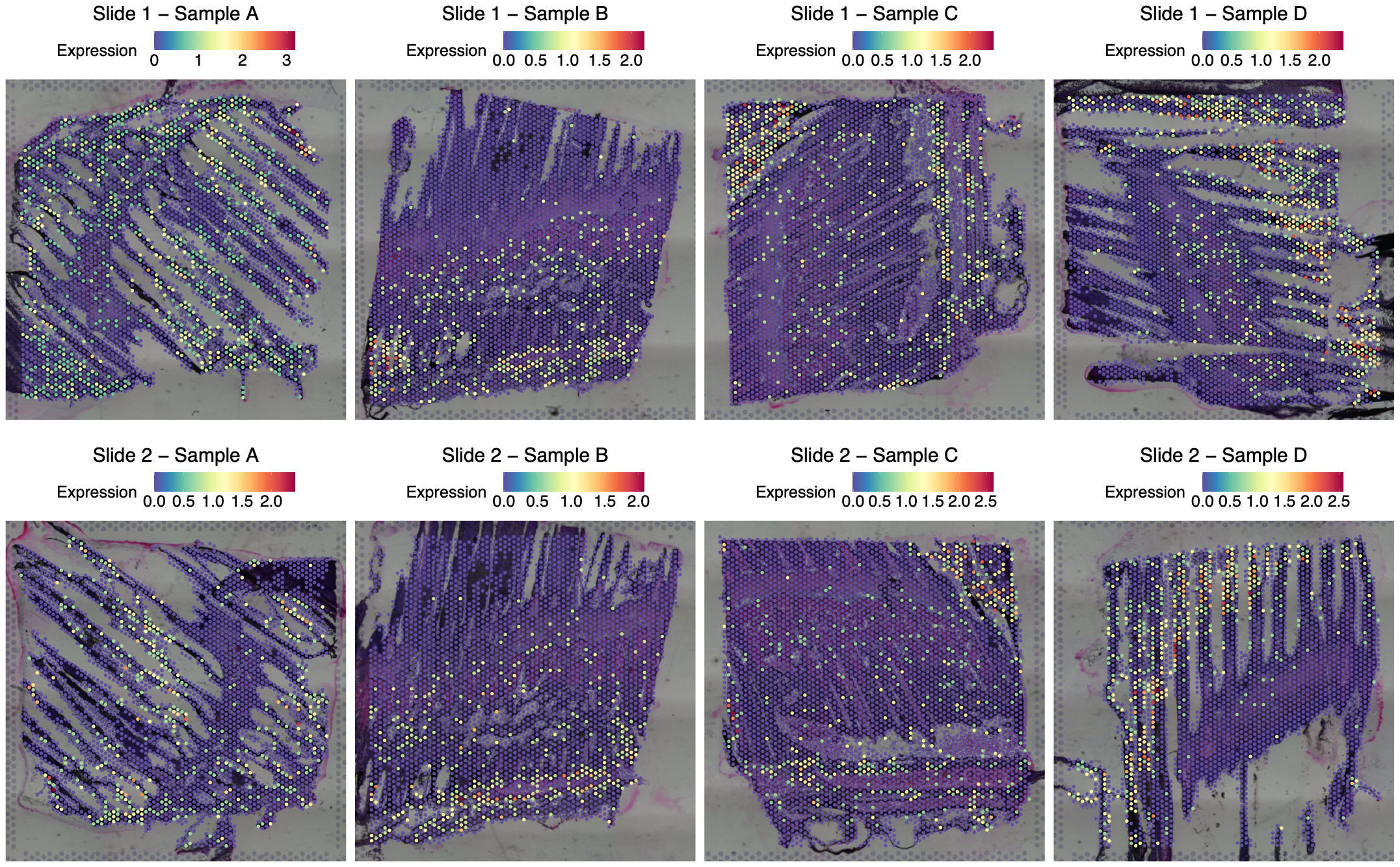

Figure S8.2.9 The spatial distribution of *igfbp5a* (insulin-like growth factor binding protein 5a; Ensembl ID: ENSSSAG00000039751; Entrez Gene ID: LOC106575566) expression in eight tissue sections (Slide 1: Samples A to D; Slide 2: Samples A to D). Each panel corresponds to one tissue section, with expression values represented by colour intensity. The colour intensity reflects normalised expression, using SCTransform, where higher values indicate greater relative expression following sequencing depth correction.
