## Supplementary Note 7 for "Spatial Transcriptomics Reveals Region-Specific Host Responses in Amoebic Gill Disease of Atlantic Salmon"

**Supplementary Note 7. Top 10 marker genes for each cluster**

The gene expression level of all genes within each cluster were compared to all the genes to the other clusters. The tables below showed the top 10 marker genes identified for each cluster. Highlighted are those marker genes that has been mentioned in the result section.

Table S7.1: Genes identified as differentially expressed by FindAllMarkers and retained as spatially variable using FindSpatiallyVariableFeatures. Shown here are results for cluster 0, gill epithelial tissue, including average log2 fold-change (avg_log2FC), adjusted p-values (p_val_adj), Ensembl gene identifiers, and predicted or known functions based on Ensembl protein domain annotations.

| **avg_log2FC** | **p_val_adj** | **cluster** | **gene** | **Potential Function** |
| --- | --- | --- | --- | --- |
| 2.70805941228447 | 0 | 0 | ENSSSAG00000065916 | *ccl28*; C-C motif chemokine 28 |
| 2.65788695258465 | 3.65059377863021E-80 | 0 | ENSSSAG00000115746 | lncRNA |
| 2.33902818037997 | 4.07991893387337E-107 | 0 | ENSSSAG00000005895 | Ortho: calcium binding protein |
| 2.08211174898902 | 0 | 0 | ENSSSAG00000108673 | Immunoglobulin-like domain superfamily |
| 2.05712206836971 | 1.20604251923419E-254 | 0 | ENSSSAG00000100297 | Immunoglobulin-like domain superfamily |
| 1.95784493713183 | 5.73809144997985E-120 | 0 | ENSSSAG00000000779 | Calycin or Lipocalin |
| 1.95784493713183 | 5.73809144997985E-120 | 0 | ENSSSAG00000004300 | Calycin or Lipocalin |
| 1.9312992898765 | 1.62073975432573E-287 | 0 | ENSSSAG00000005097 | ***sal***; Rhamnose-binding lectin |
| 1.93095995198457 | 1.03822349602324E-50 | 0 | ENSSSAG00000091532 | spp1; Osteopontin |
| 1.93095995198457 | 1.03822349602324E-50 | 0 | ENSSSAG00000095024 | Spp1; Osteopontin |

Table S7.2: Genes identified as differentially expressed by FindAllMarkers and retained as spatially variable using FindSpatiallyVariableFeatures. Shown here are results for cluster 1, hemoglobulin-related domain, including average log2 fold-change (avg_log2FC), adjusted p-values (p_val_adj), Ensembl gene identifiers, and predicted or known functions based on Ensembl protein domain annotations.

| **avg_log2FC** | **p_val_adj** | **cluster** | **gene** | **May encode for** |
| --- | --- | --- | --- | --- |
| 4.21021517175193 | 0 | 1 | ENSSSAG00000093364 | HLH, helix-loop-helix DNA-binding domain |
| 4.10828721340175 | 0 | 1 | ENSSSAG00000065229 | ***hbaa2***; hemoglobin, alpha adult 2 |
| 4.10767621885902 | 0 | 1 | ENSSSAG00000084006 | ***gp1bb;*** glycoprotein Ib platelet subunit beta |
| 4.09575331147672 | 0 | 1 | ENSSSAG00000109306 | Likely NADH-ubiquinone reductase comp. |
| 3.99397251545919 | 0 | 1 | ENSSSAG00000065233 | ***hbe1;*** Hemoglobin embryonic-1; as described in Andresen et al. (2024)Andresen et al. (2024) |
| 3.96558895823414 | 0 | 1 | ENSSSAG00000082132 | si:ch211-103n10.5. Predicted to enable DNA binding activity. |
| 3.93612874127522 | 0 | 1 | ENSSSAG00000068428 | Tetrapyrrole biosynthesis, 5-aminolevulinic acid synthase |
| 3.91220846796451 | 0 | 1 | ENSSSAG00000089308 | lncRNA |
| 3.87079436804964 | 0 | 1 | ENSSSAG00000046009 | ***hbae4***; hemoglobin alpha embryonic-4 |
| 3.86308036049317 | 0 | 1 | ENSSSAG00000007203 | ***Septin3***; septin 3 |

Table S7.3: Genes identified as differentially expressed by FindAllMarkers and retained as spatially variable using FindSpatiallyVariableFeatures. Shown here are results for cluster 2, gill epithelial tissue, including average log2 fold-change (avg_log2FC), adjusted p-values (p_val_adj), Ensembl gene identifiers, and predicted or known functions based on Ensembl protein domain annotations.

| **avg_log2FC** | **p_val_adj** | **cluster** | **gene** | **Potential function** |
| --- | --- | --- | --- | --- |
| 3.95727217553998 | 0 | 2 | ENSSSAG00000079841 | *LIPO*; lipocalin |
| 3.89014699854109 | 0 | 2 | ENSSSAG00000001874 | Serine protease 27-like |
| 3.88289679438989 | 0 | 2 | ENSSSAG00000071998 | Cytochrome P450 superfamily |
| 3.58602511662115 | 0 | 2 | ENSSSAG00000077906 | Apod; apolipoprotein D |
| 3.44321510848059 | 0 | 2 | ENSSSAG00000112110 | LncRNA |
| 3.4303493647524 | 0 | 2 | ENSSSAG00000112534 | *CLIC1*; chloride intracellular channel protein 1-like |
| 3.42733109446897 | 0 | 2 | ENSSSAG00000101879 | keratin, type I cytoskeletal 13-like |
| 3.31734543538883 | 0 | 2 | ENSSSAG00000001998 | Lysozyme-like domain containing |
| 3.23274637589781 | 0 | 2 | ENSSSAG00000087606 | AIG1-type guanine nucleotide-binding (G) domain |
| 3.22189661660732 | 0 | 2 | ENSSSAG00000041180 | Ubiquitin-conjugating enzyme/RWD-like |

Table S7.4: Genes identified as differentially expressed by FindAllMarkers and retained as spatially variable using FindSpatiallyVariableFeatures. Shown here are results for cluster 3, skeletal muscle tissue, including average log2 fold-change (avg_log2FC), adjusted p-values (p_val_adj), Ensembl gene identifiers, and predicted or known functions based on Ensembl protein domain annotations.

| **avg_log2FC** | **p_val_adj** | **cluster** | **gene** | **Potential Function** |
| --- | --- | --- | --- | --- |
| 5.34722845107993 | 0 | 3 | ENSSSAG00000099398 | Glyceraldehyde-3-phosphate dehydrogenase |
| 5.25822939429296 | 0 | 3 | ENSSSAG00000067521 | Glyceraldehyde-3-phosphate dehydrogenase |
| 5.02583725157844 | 0 | 3 | ENSSSAG00000066537 | *tnnc2.2*; troponin C2, fast skeletal type, tandem duplicate 2 |
| 4.92292675282226 | 0 | 3 | ENSSSAG00000062577 | *mlrb*; myosin regulatory light chain 2B, cardiac muscle isoform |
| 4.906293972135 | 0 | 3 | ENSSSAG00000007690 | *acta1-like*; actin alpha 1, skeletal muscle |
| 4.8775642877016 | 0 | 3 | ENSSSAG00000063443 | *fhl1a*; four and a half LIM domains 1a |
| 4.80075314033006 | 0 | 3 | ENSSSAG00000073229 | *actc1a*; actin alpha cardiac muscle 1a |
| 4.51989934419116 | 0 | 3 | ENSSSAG00000073639 | Glyceraldehyde-3-phosphate dehydrogenase |
| 4.34338561400725 | 0 | 3 | ENSSSAG00000041369 | CX6A1; cytochrome c oxidase subunit 6A |
| 4.34214680616668 | 0 | 3 | ENSSSAG00000063792 | cox4i2; cytochrome c oxidase subunit 4I2 |

Table S7.5: Genes identified as differentially expressed by FindAllMarkers and retained as spatially variable using FindSpatiallyVariableFeatures. Shown here are results for cluster 4, gill arch tissue, including average log2 fold-change (avg_log2FC), adjusted p-values (p_val_adj), Ensembl gene identifiers, and predicted or known functions based on Ensembl protein domain annotations.

| **avg_log2FC** | **p_val_adj** | **cluster** | **gene** |  |
| --- | --- | --- | --- | --- |
| 3.71772537392441 | 0 | 4 | ENSSSAG00000120863 | lncRNA |
| 3.53862031571618 | 0 | 4 | ENSSSAG00000084734 | lncRNA |
| 3.47655421387085 | 0 | 4 | ENSSSAG00000102051 | lncRNA |
| 3.44911265056986 | 0 | 4 | ENSSSAG00000055826 | C-type lectin like domain containing |
| 3.10026484667687 | 0 | 4 | ENSSSAG00000003409 | C-type lectin like domain containing |
| 3.01705446145491 | 5.1341932340152E-162 | 4 | ENSSSAG00000042007 | *cd59a*; CD59a |
| 2.91051776269042 | 0 | 4 | ENSSSAG00000111491 | lncRNA |
| 2.34847004806474 | 0 | 4 | ENSSSAG00000054088 | fatty acid binding protein 3, muscle and heart |
| 2.34100954936397 | 0 | 4 | ENSSSAG00000092624 | GPCR family 3, GABA-B receptor |
| 2.27337760708924 | 1.04238685628424E-98 | 4 | ENSSSAG00000113740 | Immunoglobulin-like domain superfamily |

Table S7.6: Genes identified as differentially expressed by FindAllMarkers and retained as spatially variable using FindSpatiallyVariableFeatures. Shown here are results for cluster 5, gill epithelial tissue, including average log2 fold-change (avg_log2FC), adjusted p-values (p_val_adj), Ensembl gene identifiers, and predicted or known functions based on Ensembl protein domain annotations.

| **avg_log2FC** | **p_val_adj** | **cluster** | **gene** | **Potential Function** |
| --- | --- | --- | --- | --- |
| 2.8506398967935 | 0 | 5 | ENSSSAG00000091158 | Uncharacterised protein |
| 2.78039535744832 | 0 | 5 | ENSSSAG00000114104 | Uncharacterised protein |
| 2.76041987593788 | 0 | 5 | ENSSSAG00000107315 | Uncharacterised protein |
| 2.75581713964422 | 0 | 5 | ENSSSAG00000094189 | Uncharacterised protein |
| 2.74419201046151 | 6.68554E-318 | 5 | ENSSSAG00000115803 | Orthologs: SMIM8 |
| 2.74049878317935 | 0 | 5 | ENSSSAG00000103001 | Orthologs: SMIM8(small integral membrane protein 8) |
| 2.72133516596939 | 0 | 5 | ENSSSAG00000091954 | Uncharacterised protein |
| 2.71146673246368 | 0 | 5 | ENSSSAG00000084152 | Orthologs: paox |
| 2.70447441048869 | 0 | 5 | ENSSSAG00000090097 | Uncharacterised protein |
| 2.69251582305642 | 0 | 5 | ENSSSAG00000098125 | Orthologs: SMIM8 |

Table S7.7: Genes identified as differentially expressed by FindAllMarkers and retained as spatially variable using FindSpatiallyVariableFeatures. Shown here are results for cluster 6, gill epithelial tissue, including average log2 fold-change (avg_log2FC), adjusted p-values (p_val_adj), Ensembl gene identifiers, and predicted or known functions based on Ensembl protein domain annotations.

| **avg_log2FC** | **p_val_adj** | **cluster** | **gene** |  |
| --- | --- | --- | --- | --- |
| 3.92736062663749 | 0 | 6 | ENSSSAG00000074950 | ***CATH-1A***; cathelicidin antimicrobial peptide |
| 3.73667166897796 | 0 | 6 | ENSSSAG00000121747 | Winged helix-like DNA-binding domain superfamily |
| 3.6042744147229 | 0 | 6 | ENSSSAG00000003986 | Lipoxygenase |
| 3.25490789748642 | 0 | 6 | ENSSSAG00000057034 | glutamate decarboxylase like 1 |
| 3.18323841288607 | 0 | 6 | ENSSSAG00000005307 | Mannose-specific lectin |
| 3.17661490286057 | 0 | 6 | ENSSSAG00000052761 | Epidermis-type lipoxygenase 3: loxe3 |
| 3.00800649382198 | 0 | 6 | ENSSSAG00000000759 | cystatin 14b, tandem duplicate 1 |
| 3.00662929700564 | 0 | 6 | ENSSSAG00000073649 | anterior gradient 2, protein disulphide isomerase family member |
| 3.0004188865359 | 0 | 6 | ENSSSAG00000049361 | transglutaminase 1 like 1 |
| 2.97923244382059 | 0 | 6 | ENSSSAG00000048372 | keratin, type I cytoskeletal 13 |

Table S7.8: Genes identified as differentially expressed by FindAllMarkers and retained as spatially variable using FindSpatiallyVariableFeatures. Shown here are results for cluster 7, gill arch tissue, including average log2 fold-change (avg_log2FC), adjusted p-values (p_val_adj), Ensembl gene identifiers, and predicted or known functions based on Ensembl protein domain annotations.

| **avg_log2FC** | **p_val_adj** | | **cluster** | | **gene** |  |
| --- | --- | --- | --- | --- | --- | --- |
| 4.89103926526838 | 2.96185443248131E-193 | 7 | | ENSSSAG00000071963 | | *thbs4a*; thrombospondin 4a |
| 4.20632687685175 | 3.99784996199243E-194 | 7 | | ENSSSAG00000082037 | | myocilin |
| 4.03464337668905 | 5.176349062916E-285 | 7 | | ENSSSAG00000073256 | | *TIMP3*; TIMP metallopeptidase inhibitor 3 |
| 3.76042098150709 | 0 | 7 | | ENSSSAG00000083363 | | lncRNA |
| 3.59002137393249 | 0 | 7 | | ENSSSAG00000093421 | | hyaluronan and proteoglycan link protein 1-like |
| 3.57726026865179 | 0 | 7 | | ENSSSAG00000020979 | | *lum*; lumican |
| 3.45201429503001 | 0 | 7 | | ENSSSAG00000085249 | | Tropoelastin |
| 3.24224354031441 | 0 | 7 | | ENSSSAG00000042764 | | follistatin-like 1a |
| 3.17360614363773 | 0 | 7 | | ENSSSAG00000116025 | | lncRNA |
| 3.15681387351194 | 0 | 7 | | ENSSSAG00000005527 | | collagen alpha-1(I) chain |

Table S7.9: Genes identified as differentially expressed by FindAllMarkers and retained as spatially variable using FindSpatiallyVariableFeatures. Shown here are results for cluster 8, gill arch tissue, including average log2 fold-change (avg_log2FC), adjusted p-values (p_val_adj), Ensembl gene identifiers, and predicted or known functions based on Ensembl protein domain annotations.

| **avg_log2FC** | **p_val_adj** | **cluster** | **gene** |  |
| --- | --- | --- | --- | --- |
| 4.43340009075944 | 0 | 8 | ENSSSAG00000100045 | cald1b; caldesmon 1b, non-muscle caldesmon-like isoform X1 |
| 4.32043479079649 | 0 | 8 | ENSSSAG00000041742 | Transmembrane protein 56-B-like |
| 3.79392476283135 | 0 | 8 | ENSSSAG00000044179 | ***acta2***; actin alpha 2, smooth muscle |
| 3.71936615278355 | 0 | 8 | ENSSSAG00000047730 | Desmin |
| 3.67176313513446 | 0 | 8 | ENSSSAG00000113814 | Tensin Phosphatase |
| 3.6524917364821 | 0 | 8 | ENSSSAG00000064672 | *lmod1b*; leiomodin 1b (smooth muscle) |
| 3.63576625539922 | 0 | 8 | ENSSSAG00000050262 | ***Smtna***; smoothelin a |
| 3.58024673490089 | 0 | 8 | ENSSSAG00000122078 | cnn1b; calponin 1, basic, smooth muscle, b |
| 3.44258107298881 | 0 | 8 | ENSSSAG00000023533 | regulator of G protein signaling 5 |
| 3.38873016598851 | 0 | 8 | ENSSSAG00000114975 | actn1; actinin, alpha 1 |

Table S7.10: Genes identified as differentially expressed by FindAllMarkers and retained as spatially variable using FindSpatiallyVariableFeatures. Shown here are results for cluster 9, gill arch tissue, including average log2 fold-change (avg_log2FC), adjusted p-values (p_val_adj), Ensembl gene identifiers, and predicted or known functions based on Ensembl protein domain annotations.

| **avg_log2FC** | **p_val_adj** | **cluster** | **gene** | Encoded for |
| --- | --- | --- | --- | --- |
| 5.04485855227398 | 0 | 9 | ENSSSAG00000088534 | GP182; G-protein coupled receptor 182 |
| 4.97919580397469 | 0 | 9 | ENSSSAG00000069027 | Tissue factor pathway inhibitor 2 |
| 4.79701747045672 | 0 | 9 | ENSSSAG00000073565 | ***Mrc1*** (mannose receptor C-type 1) |
| 4.74961627861206 | 0 | 9 | ENSSSAG00000005576 | SPARC-like protein 1 |
| 4.27611706441302 | 0 | 9 | ENSSSAG00000001017 | cfd; complement factor D (adipsin) |
| 4.04342957913931 | 0 | 9 | ENSSSAG00000047898 | Carboxypeptidase E |
| 3.91507029951151 | 0 | 9 | ENSSSAG00000077688 | GP182; G-protein coupled receptor 182 |
| 3.88758324318333 | 0 | 9 | ENSSSAG00000098957 | Lymphatic vessel endothelial hyaluronic acid receptor 1-like |
| 3.79667693922593 | 0 | 9 | ENSSSAG00000117627 | IGF binding protein 6 precursor, Insulin-like growth factor binding protein 6 paralog B1, Insulin-like growth factor-binding protein 6 |
| 3.71152923189619 | 0 | 9 | ENSSSAG00000072759 | xlkd1; Lymphatic vessel endothelial hyaluronic acid receptor 1 |

Table S7.11: Genes identified as differentially expressed by FindAllMarkers and retained as spatially variable using FindSpatiallyVariableFeatures. Shown here are results for cluster 10, gill epithelial tissue, including average log2 fold-change (avg_log2FC), adjusted p-values (p_val_adj), Ensembl gene identifiers, and predicted or known functions based on Ensembl protein domain annotations.

| **avg_log2FC** | **p_val_adj** | **cluster** | **gene** | **Potential Function** |
| --- | --- | --- | --- | --- |
| 3.03308570123039 | 0 | 10 | ENSSSAG00000051243 | platelet-activating factor acetylhydrolase, isoform Ib, gamma subunit |
| 2.94962057296078 | 6.9243994822798E-90 | 10 | ENSSSAG00000089279 | Phospholipase A2 inhibitor 31 kDa subunit-like |
| 2.60471232284828 | 3.62507245926096E-291 | 10 | ENSSSAG00000116244 | C-type lectin like domain containing |
| 2.55223012272194 | 0 | 10 | ENSSSAG00000033609 | C-C motif chemokine 19-like |
| 2.47055671731106 | 6.62351846361557E-207 | 10 | ENSSSAG00000118957 | lncRNA |
| 2.39650273975041 | 4.2488698730403E-65 | 10 | ENSSSAG00000121747 | Winged helix-like DNA-binding domain superfamily |
| 2.36816927290875 | 5.05761315468074E-143 | 10 | ENSSSAG00000110347 | DEDDh 3'-5' exonuclease domain family |
| 2.34191323164849 | 2.02177755195019E-190 | 10 | ENSSSAG00000122034 | lncRNA |
| 2.26383714179497 | 8.68902769995009E-200 | 10 | ENSSSAG00000045259 | AIG1-type guanine nucleotide-binding (G) domain |
| 2.22432651675293 | 3.07656092136611E-223 | 10 | ENSSSAG00000045452 | *krt15*; keratin 15 |

Table S7.12: Genes identified as differentially expressed by FindAllMarkers and retained as spatially variable using FindSpatiallyVariableFeatures. Shown here are results for cluster 11, gill epithelial tissue, including average log2 fold-change (avg_log2FC), adjusted p-values (p_val_adj), Ensembl gene identifiers, and predicted or known functions based on Ensembl protein domain annotations.

| **avg_log2FC** | **p_val_adj** | **cluster** | **gene** | **Potential Function** |
| --- | --- | --- | --- | --- |
| 2.42409092221503 | 9.15859347239635E-197 | 11 | ENSSSAG00000090328 | collagen alpha-2(VI) chain |
| 2.39100145581574 | 1.13753929292138E-176 | 11 | ENSSSAG00000114656 | *col10a1b*; collagen, type X, alpha 1b |
| 2.18030113829611 | 6.59728491910489E-243 | 11 | ENSSSAG00000084734 | lncRNA |
| 2.14204313571724 | 2.33231987377662E-73 | 11 | ENSSSAG00000111491 | lncRNA |
| 2.08840595343462 | 2.82866505408071E-82 | 11 | ENSSSAG00000100947 | spp1; Osteopontin |
| 2.01302279237108 | 5.9691352304981E-242 | 11 | ENSSSAG00000055826 | C-type lectin like domain containing |
| 1.91712550959101 | 3.38106281384662E-27 | 11 | ENSSSAG00000076592 | tyrosinase related protein 1 |
| 1.8992945631045 | 1.29907237925882E-65 | 11 | ENSSSAG00000115746 | lncRNA |
| 1.88479627877617 | 3.38575357047408E-66 | 11 | ENSSSAG00000000157 | ***mpx***; Myeloid-specific peroxidase |
| 1.79417818709674 | 3.06266958219724E-76 | 11 | ENSSSAG00000079745 | Metallopeptidase, catalytic domain superfamily |

Table S7.13: Genes identified as differentially expressed by FindAllMarkers and retained as spatially variable using FindSpatiallyVariableFeatures. Shown here are results for cluster 12, gill epithelial tissue, including average log2 fold-change (avg_log2FC), adjusted p-values (p_val_adj), Ensembl gene identifiers, and predicted or known functions based on Ensembl protein domain annotations.

| **avg_log2FC** | **p_val_adj** | **cluster** | **gene** | **Potential Function** |
| --- | --- | --- | --- | --- |
| 3.51222070337516 | 0 | 12 | ENSSSAG00000055927 | sodium/hydrogen exchanger 3 |
| 3.29256762938362 | 0 | 12 | ENSSSAG00000084204 | Synaptobrevin/Vesicle-associated membrane protein |
| 3.16046744040734 | 0 | 12 | ENSSSAG00000104002 | Ion-transport regulator, FXYD family |
| 3.13077859147755 | 4.501038606E-315 | 12 | ENSSSAG00000056493 | Ion-transport regulator, FXYD family |
| 2.93763660683861 | 1.03724809507778E-305 | 12 | ENSSSAG00000088702 | P-type Cation transport ATPase |
| 2.8879208182184 | 8.42555842787236E-224 | 12 | ENSSSAG00000088363 | igfbp5a; insulin-like growth factor binding protein 5a |
| 2.83499651248165 | 2.81252089905946E-284 | 12 | ENSSSAG00000103110 | lncRNA |
| 2.72003709523641 | 8.32816630946042E-215 | 12 | ENSSSAG00000059186 | *trpv6*; transient receptor potential cation channel, subfamily V, member 6 |
| 2.60177074361169 | 1.00726260340131E-159 | 12 | ENSSSAG00000039751 | igfbp5a; insulin-like growth factor binding protein 5a |
| 2.53552044187731 | 1.96269858391604E-198 | 12 | ENSSSAG00000091602 | Transient receptor potential cation channel M |

Table S7.14: Genes identified as differentially expressed by FindAllMarkers and retained as spatially variable using FindSpatiallyVariableFeatures. Shown here are results for cluster 13, gill epithelial tissue, including average log2 fold-change (avg_log2FC), adjusted p-values (p_val_adj), Ensembl gene identifiers, and predicted or known functions based on Ensembl protein domain annotations.

| **avg_log2FC** | **p_val_adj** | **cluster** | **gene** | **Potential function** |
| --- | --- | --- | --- | --- |
| 5.42383869955706 | 1.61320198008358E-259 | 13 | ENSSSAG00000048372 | Keratin, type I cytoskeletal 13-like |
| 3.72915442802763 | 8.56876123675411E-156 | 13 | ENSSSAG00000057034 | glutamate decarboxylase like 1 |
| 3.70652660842643 | 2.68373044930715E-158 | 13 | ENSSSAG00000074950 | CATH-1A; cathelicidin antimicrobial peptide |
| 3.61740314675849 | 7.75764182987351E-142 | 13 | ENSSSAG00000089279 | Phospholipase A2 inhibitor 31 kDa subunit-like |
| 3.29827042980839 | 1.55318023617572E-56 | 13 | ENSSSAG00000045448 | Interleukin-1 |
| 3.29520624371937 | 1.45328303427731E-155 | 13 | ENSSSAG00000110616 | Uncharacterized protein |
| 3.22273688765334 | 2.90886271868259E-77 | 13 | ENSSSAG00000077262 | E3 ubiquitin-protein ligase LINCR |
| 3.21995000258679 | 1.88968446896666E-25 | 13 | ENSSSAG00000061390 | cxl10; C-X-C motif chemokine 10 |
| 3.03236386667724 | 1.94564379308293E-81 | 13 | ENSSSAG00000068317 | MARVEL domain-containing and chemokine-like factor proteins. |
| 2.9765549618734 | 3.22328175186707E-63 | 13 | ENSSSAG00000110347 | Ribonuclease H superfamily |
