## Supplementary Note 11 for "Spatial Transcriptomics Reveals Region-Specific Host Responses in Amoebic Gill Disease of Atlantic Salmon"

**Supplementary Note 11.** **Transcriptional Identification of Erythroid and Platelet-Associated Signatures in 3.2.4 Haemoglobulin cluster (Cluster 1)**

Top marker genes in Cluster 1 suggest transcriptional features linked to both vascular and erythroid functions, indicating a potential role in oxygen transport and vascular integrity. Spatially, this cluster is primarily localised within blood vessels of the gill arch and filaments, consistent with its association with the broader circulatory network (Figure 33).

Elevated expression of red blood cell associated genes, and their metabolic components further supports a role in oxygen binding and transport. Notably, high expression of haemoglobin genes, including ***hbaa2*** (hemoglobin, alpha adult 2; Ensembl ID: ENSSSAG00000065229; Cluster 1; Figure S11.1), ***hbae4*** (hemoglobin alpha embryonic-4; Ensembl ID: ENSSSAG00000046009; Cluster 1; Figure S11.2), and ***hbe1*** (Hemoglobin embryonic-1; Ensembl ID: ENSSSAG00000065233; Cluster 1; Figure S11.3) highlights a erythroid signature corresponding the Andresen et al. (2024) study. Additionally, increased levels of **ENSSSAG00000068428** (Tetrapyrrole biosynthesis, 5-aminolevulinic acid synthase; InterPro: IPR010961; Cluster 1; ; Figure S11.4) may reflect active haem biosynthesis which is important for haemoglobin production.

***hbaa2* (ENSSSAG00000065229)**

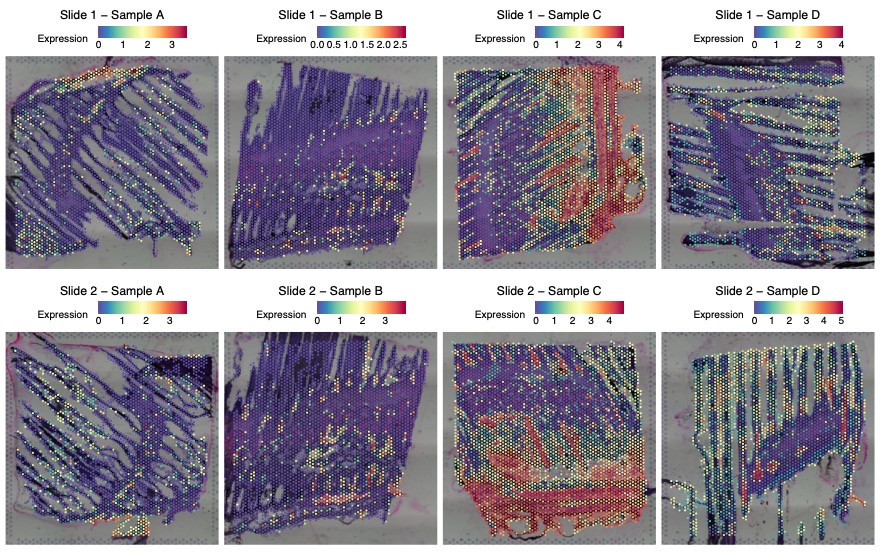

Figure S11.1 The spatial distribution of *hbaa2* (hemoglobin, alpha adult 2; Ensembl ID: ENSSSAG00000065229) expression in eight tissue sections (Slide 1: Samples A to D; Slide 2: Samples A to D). Each panel corresponds to one tissue section, with expression values represented by colour intensity. The colour intensity reflects normalised expression, using SCTransform, where higher values indicate greater relative expression following sequencing depth correction.

***hbae4* (ENSSSAG00000046009)**

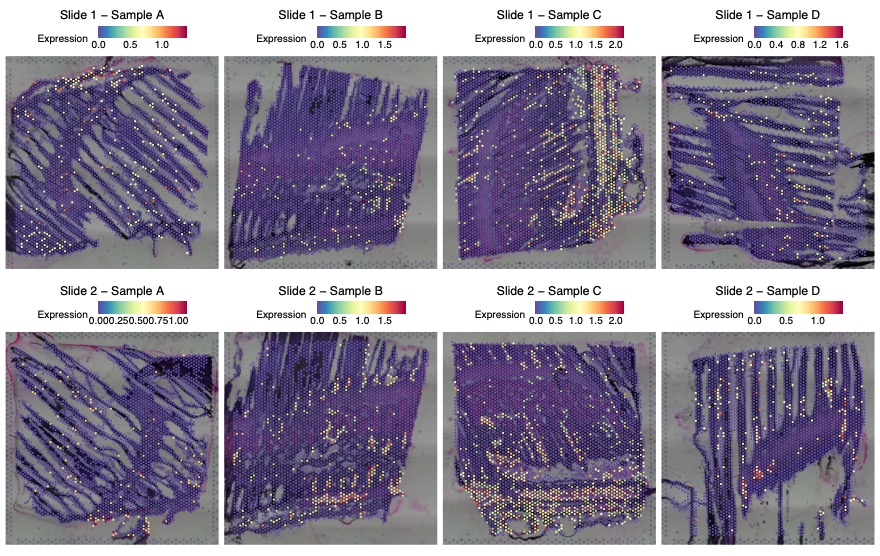

Figure S11.2 The spatial distribution of *hbae4* (hemoglobin alpha embryonic-4; Ensembl ID: ENSSSAG00000046009) expression in eight tissue sections (Slide 1: Samples A to D; Slide 2: Samples A to D). Each panel corresponds to one tissue section, with expression values represented by colour intensity. The colour intensity reflects normalised expression, using SCTransform, where higher values indicate greater relative expression following sequencing depth correction.

***hbe1* (ENSSSAG00000065233)**

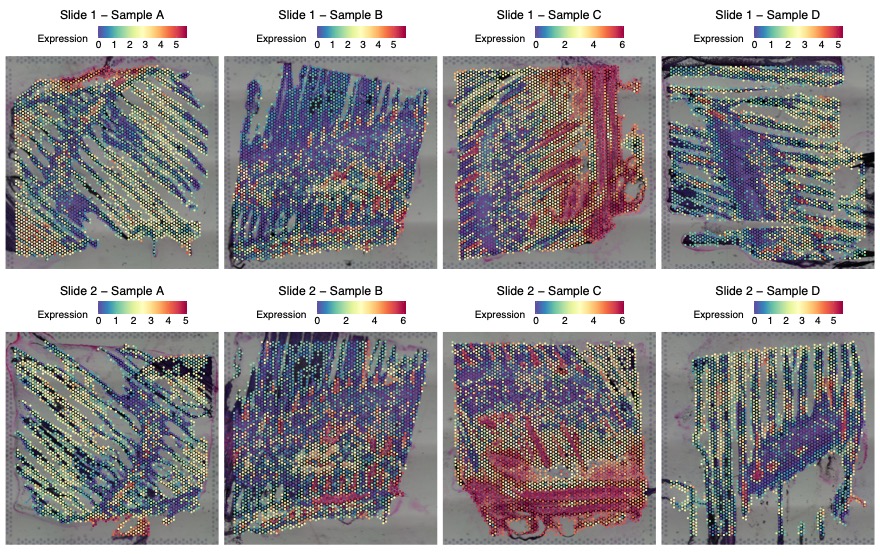

Figure S11.3 The spatial distribution of hbe1 (Hemoglobin embryonic-1; Ensembl ID: ENSSSAG00000065233) expression in eight tissue sections (Slide 1: Samples A to D; Slide 2: Samples A to D). Each panel corresponds to one tissue section, with expression values represented by colour intensity. The colour intensity reflects normalised expression, using SCTransform, where higher values indicate greater relative expression following sequencing depth correction.

**ENSSSAG00000068428**

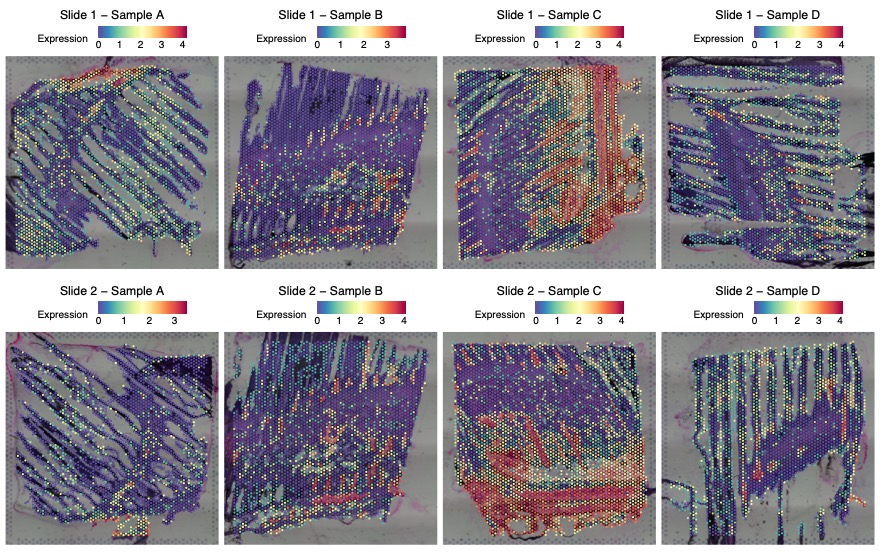

Figure S11.4 The spatial distribution of ENSSSAG00000068428 (Tetrapyrrole biosynthesis, 5-aminolevulinic acid synthase; InterPro: IPR010961) expression in eight tissue sections (Slide 1: Samples A to D; Slide 2: Samples A to D). Each panel corresponds to one tissue section, with expression values represented by colour intensity. The colour intensity reflects normalised expression, using SCTransform, where higher values indicate greater relative expression following sequencing depth correction.

Apart from erythroid markers, genes indicative of vascular and structural support are among the top marker genes in Cluster 1. ***gp1bb*** (glycoprotein Ib platelet subunit beta; Ensembl ID: ENSSSAG00000084006; Cluster 1; Figure S11.5), one of the top marker genes in Cluster 1, may reflect platelet involvement in maintaining vascular integrity and facilitating repair. Similarly, ***septin3*** (septin 3; Ensembl ID: ENSSSAG00000007203; Cluster 1; Figure S11.6), another top marker, belongs to the septin family of cytoskeletal proteins, several of which are well established in platelet structure and function(Kim *et al.*, 2022).

***gp1bb* (ENSSSAG00000084006)**

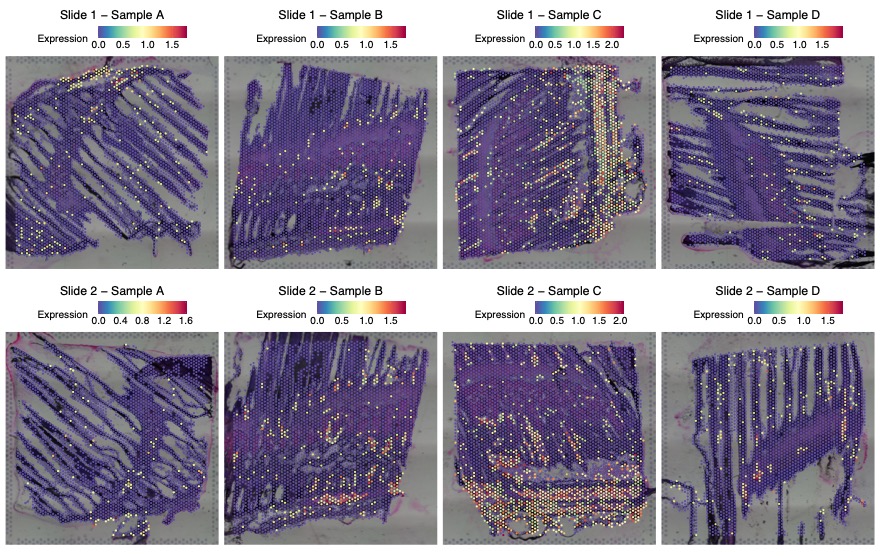

Figure S11.5 The spatial distribution of *gp1bb* (glycoprotein Ib platelet subunit beta; Ensembl ID: ENSSSAG00000084006) expression in eight tissue sections (Slide 1: Samples A to D; Slide 2: Samples A to D). Each panel corresponds to one tissue section, with expression values represented by colour intensity. The colour intensity reflects normalised expression, using SCTransform, where higher values indicate greater relative expression following sequencing depth correction.

***septin3* (ENSSSAG00000007203)**

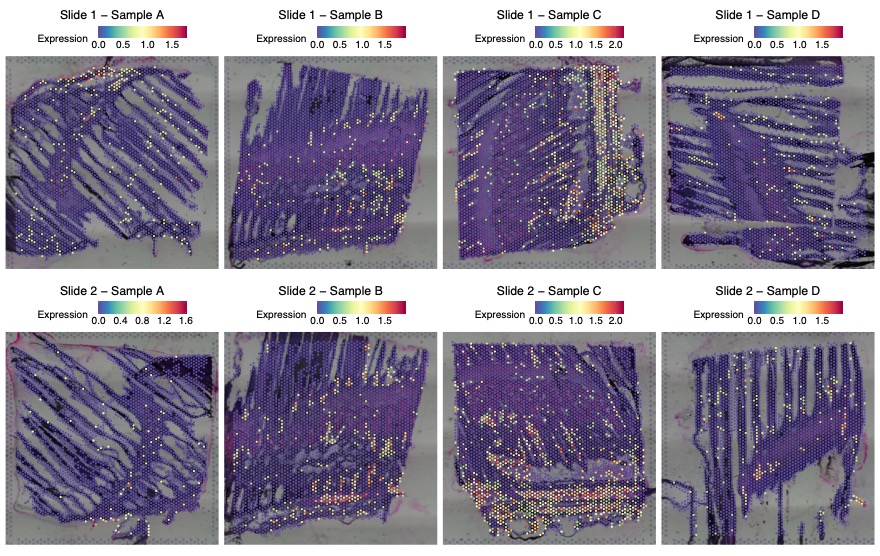

Figure S11.6 The spatial distribution of *septin3* (septin 3; Ensembl ID: ENSSSAG00000007203) expression in eight tissue sections (Slide 1: Samples A to D; Slide 2: Samples A to D). Each panel corresponds to one tissue section, with expression values represented by colour intensity. The colour intensity reflects normalised expression, using SCTransform, where higher values indicate greater relative expression following sequencing depth correction.
