## Supplementary Note 12 for "Spatial Transcriptomics Reveals Region-Specific Host Responses in Amoebic Gill Disease of Atlantic Salmon"

**Supplementary Note 1****2.** Integrating Spatial and Single Nuclei Transcriptomics in Gills

To address cellular heterogeneity within the gill-associated lymphoid tissue (GIALT) and to gain a higher-resolution understanding of the spatial arrangement specific immune and structural components that support mucosal defence, our spatial transcriptomics dataset was integrated with single-nucleus RNA sequencing data from West et al. (2021). GIALT encompasses both the immune cell populations directly engaged in pathogen recognition and response, as well as the surrounding epithelial and structural framework that maintains the mucosal barrier and facilitates immune cell trafficking. Briefly, West et al. (2021) identified 20 distinct cell clusters in their sn-RNA experiment, including pavement cells, erythrocytes, four subtypes of vascular cells, pillar cells, mitochondrion-rich cells (MRCs) from both freshwater (FW) and seawater (SW) conditions, T cells, two subtypes of epithelial cells, goblet cells, non-differentiated cells, neuroepithelial cells, monocytes, lymphatic cells, dendritic cells, accessory cells, and fibrocytes. The integration was performed using RCTD in doublet mode, which constrains each spatial spot to contain at most two cell types and assumes that spots may represent transcriptomes from up to two distinct cells. Across the eight samples, an average of 51.2% of spatial spots were classified as singlets, representing areas dominated by a single cell type. 31.2% were categorised as doublets, while 17.6% were rejected due to unassigned cell type profiles (Table 2). Doublet spots reflect the close spatial proximity of two cell types, which may suggest for potential cell–cell interactions. These interactions are especially important in the gill, where the cooperation between structural and immune cells is essential for preserving barrier integrity and coordinating mucosal defence. The distribution of these cell types, as assigned by RCTD based on the West et al. (2021) reference data, is shown across the gills from Figure 20 to Figure 23. All 20 cell types were recovered in the integrated dataset, five cell types with immunological relevance includes S12.1 T cells, S12.2 dendritic cells, S12.3 monocytes, S12.4 lymphatic cells, and S12.5 goblet cells. The following sections examined their spatial associations with other cell types as revealed through doublet classification.

**12.1 T cells**

T cells are a type of lymphocyte essential for adaptive immunity that identify specific antigens and coordinate the immune response. Across all samples, T cell-containing spots were predominantly classified as doublets ranging from 69.1 to 100%, with a minority of spots classified as singlets (ranging from 0.0 to 30.9%) (Figure, inner rings). Spatial mapping revealed that these T cell-containing spots were consistently localised to the filaments and lamellae of the gill, rather than the arch region (Figure 20 to Figure 23). This spatial arrangement suggests that T cells are integrated within the peripheral epithelial-vascular region, which constitutes the main mucosal interface with the external environment.

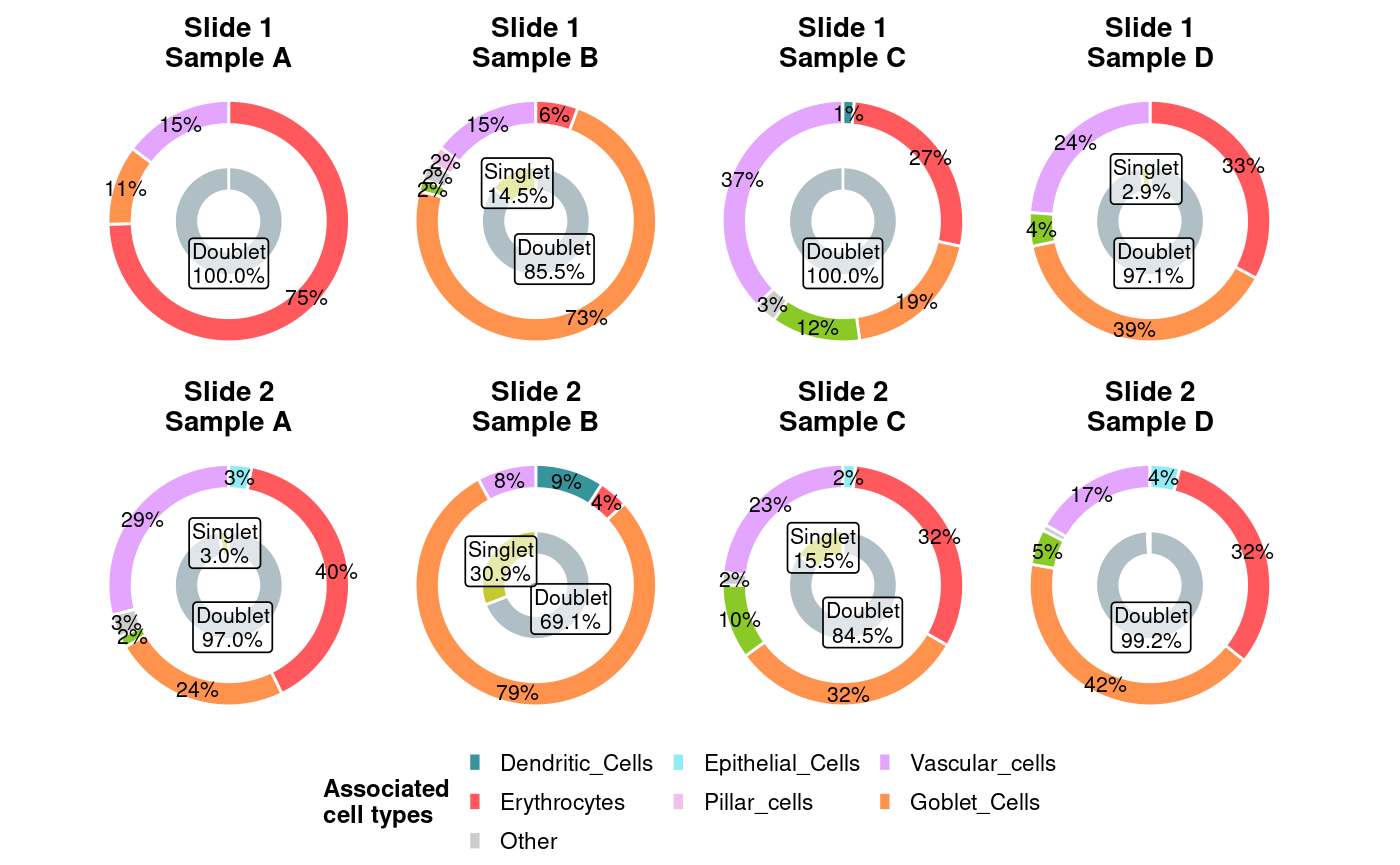

Figure S12.1.1: Distribution of T cell singlets and doublet associations across eight gill samples. The inner ring of each doughnut chart represents the proportion of T cell containing spots classified as singlets (dominated by T cells alone) versus doublets (T cells co-localising with another cell type). The outer ring shows the distribution of associated partner cell types within the doublet fraction. Only the top five most abundant associated cell types are displayed per sample; all other associations are collapsed into the “Other” category. Colours for each cell type correspond to the guide legend. Percentages are calculated relative to all T cells containing spots in each sample.

Across T cells-containing doublet spots, T cells were found frequently associated with goblet cells, erythrocytes, and vascular cells, with additional associations involving lymphatic and dendritic cells with variation across samples (**Figure** **S12.1.1**, outer rings). In Sample A, the most frequent pairings were with erythrocytes (40–75%), followed by vascular cells (15–29%) and goblet cells (11–24%), with smaller contributions from epithelial (3%) and lymphatic cells (2%). In Sample B, associations were dominated by goblet cells (73–79%), with additional contributions from vascular cells (8–15%), dendritic cells (9%), erythrocytes (4–6%), and occasional pillar cells (2%). Sample C displayed a mixed profile, with vascular cells (23–37%), erythrocytes (27–32%), and goblet cells (19–32%) as the dominant associated cell types, followed by smaller contributions from lymphatic (10–12%), dendritic (1%), and epithelial (2%) cells. In Sample D, goblet cells were again the dominant associated cell types (39–42%), accompanied by erythrocytes (32–33%) and vascular cells (17–24%), with smaller contributions from lymphatic (4–5%) and epithelial (4%) cells.

**11. 2 Dendritic cells**

Dendritic cells, a type of antigen-presenting cell, play an important role to capture, process, and present antigens to lymphocytes, thereby bridging innate and adaptive immunity. Dendritic cells were classified by RCTD as both singlets (26.7-62.4%) and doublets (37.6-73.3%), with their proportions varying across samples (Figure, inner ring). These cells were primarily clustered at the gill arch, though they were also distributed along the filaments, placing them at critical sites for both antigen entry and vascular exchange (**Error! Reference source not found.** - **Error! Reference source not found.**). Such localisation aligns with the sentinel role of dendritic cells as antigen-presenting cells, positioning them to bridge innate and adaptive immune responses within the GIALT.

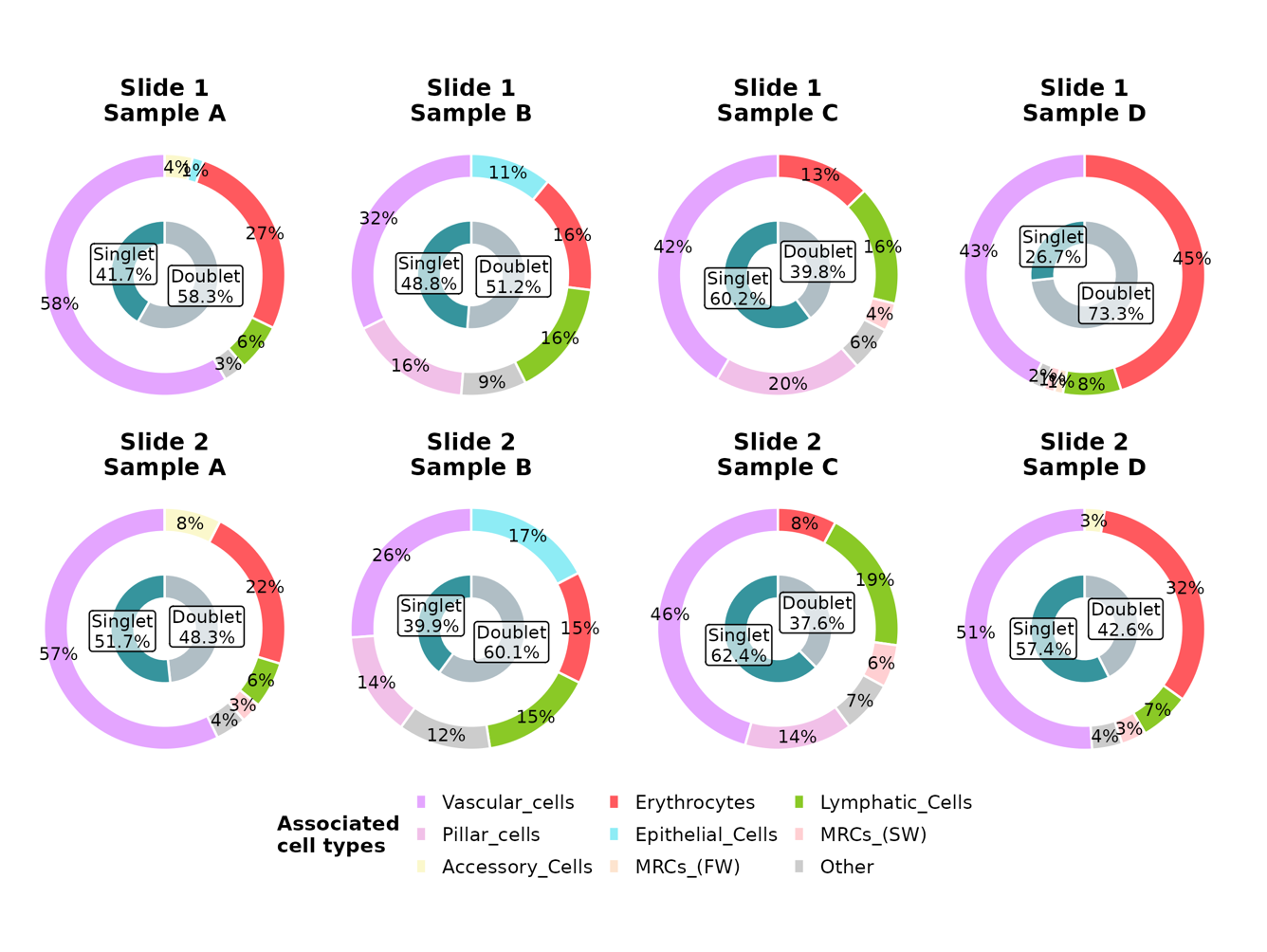

Figure S12.2.1: Distribution of dendritic cell singlets and doublet associations across eight gill samples. The inner ring of each doughnut chart represents the proportion of dendritic cell containing spots classified as singlets (dominated by dendritic cells alone) versus doublets (dendritic cells co-localising with another cell type). The outer ring shows the distribution of associated partner cell types within the doublet fraction. Only the top five most abundant associated cell types are displayed per sample; all other associations are collapsed into the “Other” category. Colours for each cell type correspond to the guide legend. Percentages are calculated relative to all dendritic cell-containing spots in each sample.

Among doublet assignments, dendritic cells are most frequently found co-localising with vascular cells and erythrocytes, highlighting their close association with the water-blood interface (Figure, outer ring). In Sample A, dendritic cells were predominantly paired with vascular cells (57 - 58%) and erythrocytes (22 - 27%), alongside accessory (4 - 8%) and lymphatic cells (6%), with smaller contributions from epithelial cells and mitochondrion-rich cells (MRCs). Sample B showed more diverse co-localisation with vascular cells (26 - 32%), pillar cells (14 - 16%), erythrocytes (15 - 16%), lymphatic cells (15 - 16%), and epithelial cells (11 - 17%) all appearing as frequent associated cell types. In Sample C, vascular cells again dominated (42 - 46%), followed by pillar cells (14 - 20%), lymphatic cells (16 - 19%), erythrocytes (8 - 13%), and MRCs (4 - 6%). In Sample D, both erythrocytes (32 - 45%) and vascular cells (43 - 51%) were the dominant associated cell type, followed by smaller proportions of lymphatic cells (7 - 8%), MRCs (1 - 3%), and accessory cells (3%). Across samples, frequent DC co-localisation ranges were vascular 26–58%, erythrocyte 8–45%, pillar 14–20%, lymphatic 6–19%, epithelial 11–17%, goblet 4–9% and MRCs 1–6% (Figure 1.90).

**11.3 Monocytes**

Teleost monocytes are phagocytic leukocytes circulating in the blood, serving as precursors to macrophages and playing a central role in both innate and adaptive immunity (Lu and Chen, 2019; Zhu and Su, 2022). Their primary functions include phagocytosis, antigen presentation, cytokine production, and orchestration of antimicrobial responses. Monocytes were found scattered throughout the gill, with no clear restriction to the arch, filament, or lamellae (**Error! Reference source not found.** - **Error! Reference source not found.**). All monocytes were classified in doublets (100%), suggesting they always appeared together with other immune or structural cell types in a close proximity rather than existing as isolated singlets (Figure, inner ring).

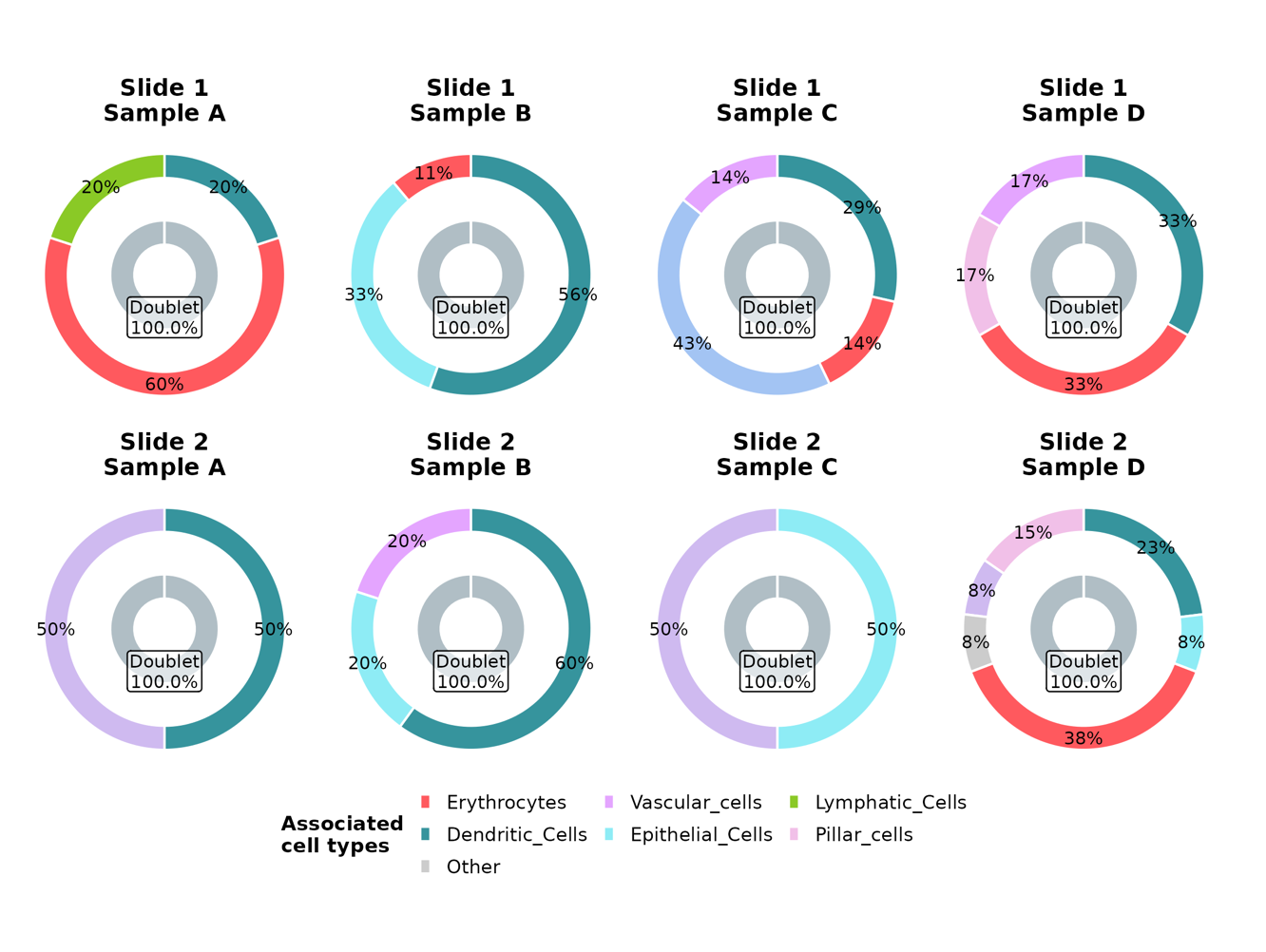

Figure S12.3.1 Distribution of monocyte singlets and doublet associations across eight gill samples. The inner ring of each doughnut chart represents the proportion of monocyte-containing spots classified as singlets (dominated by monocytes alone) versus doublets (monocyte co-localising with another cell type). The outer ring shows the distribution of associated partner cell types within the doublet fraction. Only the top five most abundant associated cell types are displayed per sample; all other associations are collapsed into the “Other” category. Colours for each cell type correspond to the guide legend. Percentages are calculated relative to all monocyte-containing spots in each sample.

Within monocytes-containing doublet spots, monocytes consistently co-localising with erythrocytes and dendritic cells, though their spatial association varied across samples (Figure, outer ring). This reflects the flexible integration of monocytes into vascular, epithelial, and immune niches. In Sample A, monocytes most frequently paired with erythrocytes (60%), followed by dendritic cells (20%) and lymphatic cells (20%) in Slide 1, while Slide 2 showed equal associations with dendritic cells (50%) and pavement cells (50%). In Sample B, dendritic cells were the dominant neighbours (56 - 60%), with epithelial cells (20 - 33%) and erythrocytes (11 - 15%) also represented; vascular cells contributed 20% in Slide 2. In Sample C, monocytes are found spatially associated with non-differentiated cells (43%), dendritic cells (29%), erythrocytes (14%), and vascular cells (14%) in Slide 1; whereas in Slide 2, epithelial and pavement cells were equally represented (50% respectively). In Sample D, monocytes were found colocalising with dendritic cells (33%), erythrocytes (33%), pillar cells (17%), and vascular cells (17%) in Slide 1; while Slide 2 highlighted erythrocytes (38%) and dendritic cells (23%) as frequent neighbours, with additional contributions from pillar (15%), epithelial (8%), and pavement cells (8%).

**11.4 Lymphatic cells**

In this study, the lymphatic cell population was defined following West *et al.* (2021), who characterised these cells by their typical *Prox1* expression patterns in single-nuclei transcriptomics. These lymphatic cells were abundant across all samples, with widespread localisation throughout both the gill arch and filaments (**Error! Reference source not found.** - **Error! Reference source not found.**). Most lymphatic cell containing spots were classified as doublets, ranging from 96 to 100% across slides, reflecting the strong tendency of lymphatic cells to associate with other cell types rather than occur as isolated singlets (Figure , inner ring).

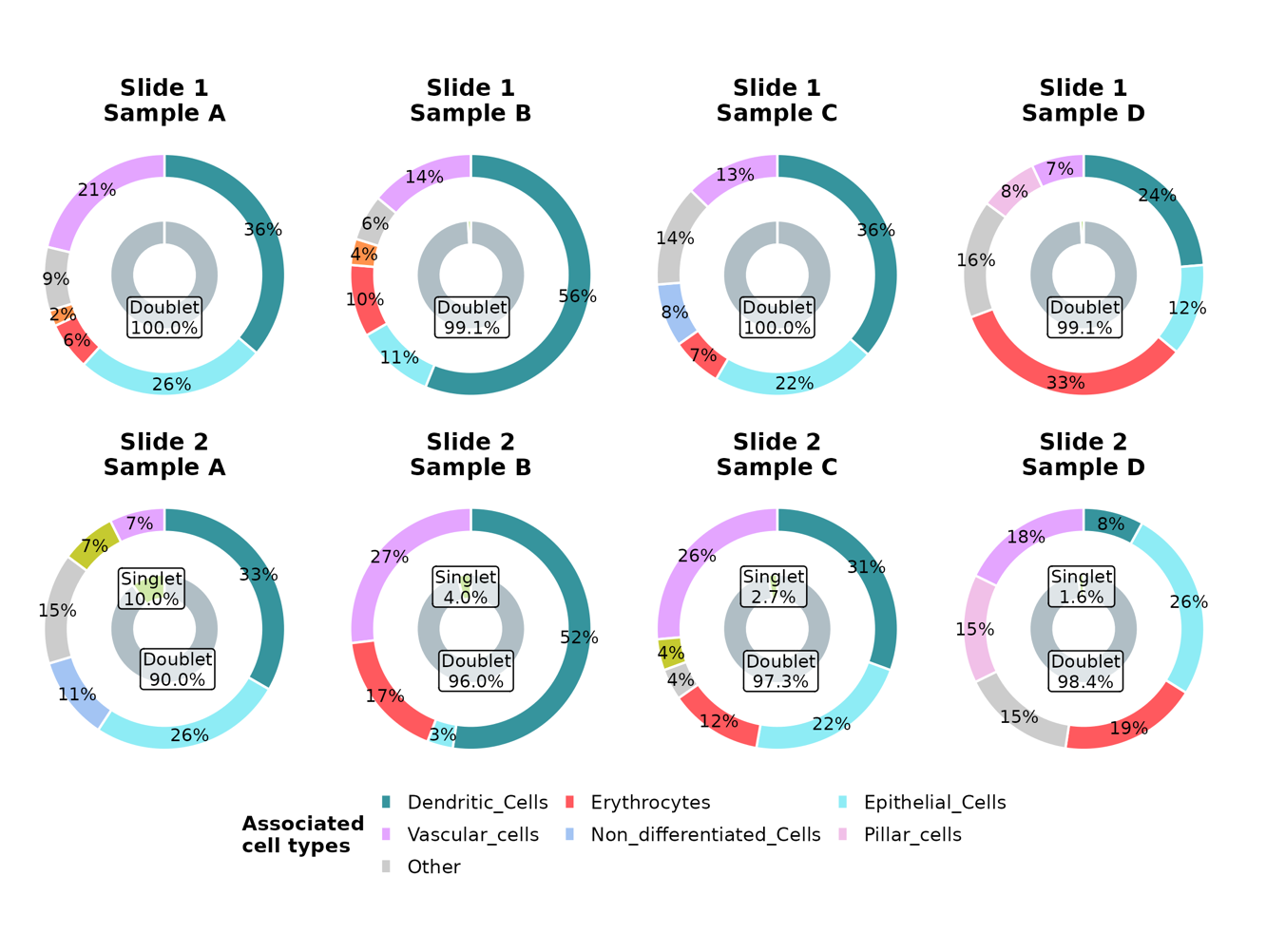

Figure S12.4.1 Distribution of lymphatic cell singlets and doublet associations across eight gill samples. The inner ring of each doughnut chart represents the proportion of lymphatic cell-containing spots classified as singlets (dominated by lymphatic cells alone) versus doublets (lymphatic cells co-localising with another cell type). The outer ring shows the distribution of associated partner cell types within the doublet fraction. Only the top five most abundant associated cell types are displayed per sample; all other associations are collapsed into the “Other” category. Colours for each cell type correspond to the guide legend. Percentages are calculated relative to all lymphatic cell-containing spots in each sample.

Lymphatic cell-containing doublet spots were consistently associated with dendritic, epithelial, vascular, and erythrocyte populations, with additional but less frequent associations involving goblet, pillar, and T cells depending on the sample (Figure ). In Sample A, lymphatic cells were most commonly paired with dendritic cells (33 - 36%) and epithelial cells (26%), followed by vascular cells (7 - 21%) and erythrocytes (6%), with smaller contributions from goblet cells (2%) and T cells (7%). In Sample B, lymphatic cells were frequently found in spatial association with dendritic cells (52 - 56%), followed by vascular cells (14 - 27%) and erythrocytes (10 - 17%), with additional associations involving epithelial cells (3 - 11%). In Sample C, lymphatic cells consistently showed spatial associations with dendritic cells (31 - 36%) and epithelial cells (22%), while also co-localising with vascular cells (13 - 26%), erythrocytes (7 - 12%), T cells (4%) and non-differentiated cells (8%). In Sample D, lymphatic cell associations were more evenly distributed, involving erythrocytes (19 - 33%), dendritic cells (8 - 24%), epithelial cells (12 - 26%), vascular cells (7 - 18%), and pillar cells (8 - 15%).

**11.5 Goblet cells**

In teleosts, goblet cells are mucus-secreting epithelial cells that contribute to mucosal immunity by producing and releasing mucins rich in antimicrobial molecules (such as lysozymes and lectins), which trap and neutralise pathogens while supporting the physical and chemical barrier of the skin, gills, and gut. Goblet cells were abundant across all samples, with predominant localisation in the filament and lamellar regions of the gill (**Error! Reference source not found.** - **Error! Reference source not found.**). Most goblet cell containing spots are doublets, ranging from 95 to 100% across slides, suggesting frequent co-localisation with other cell types rather than singlet distributions (Figure, inner ring).

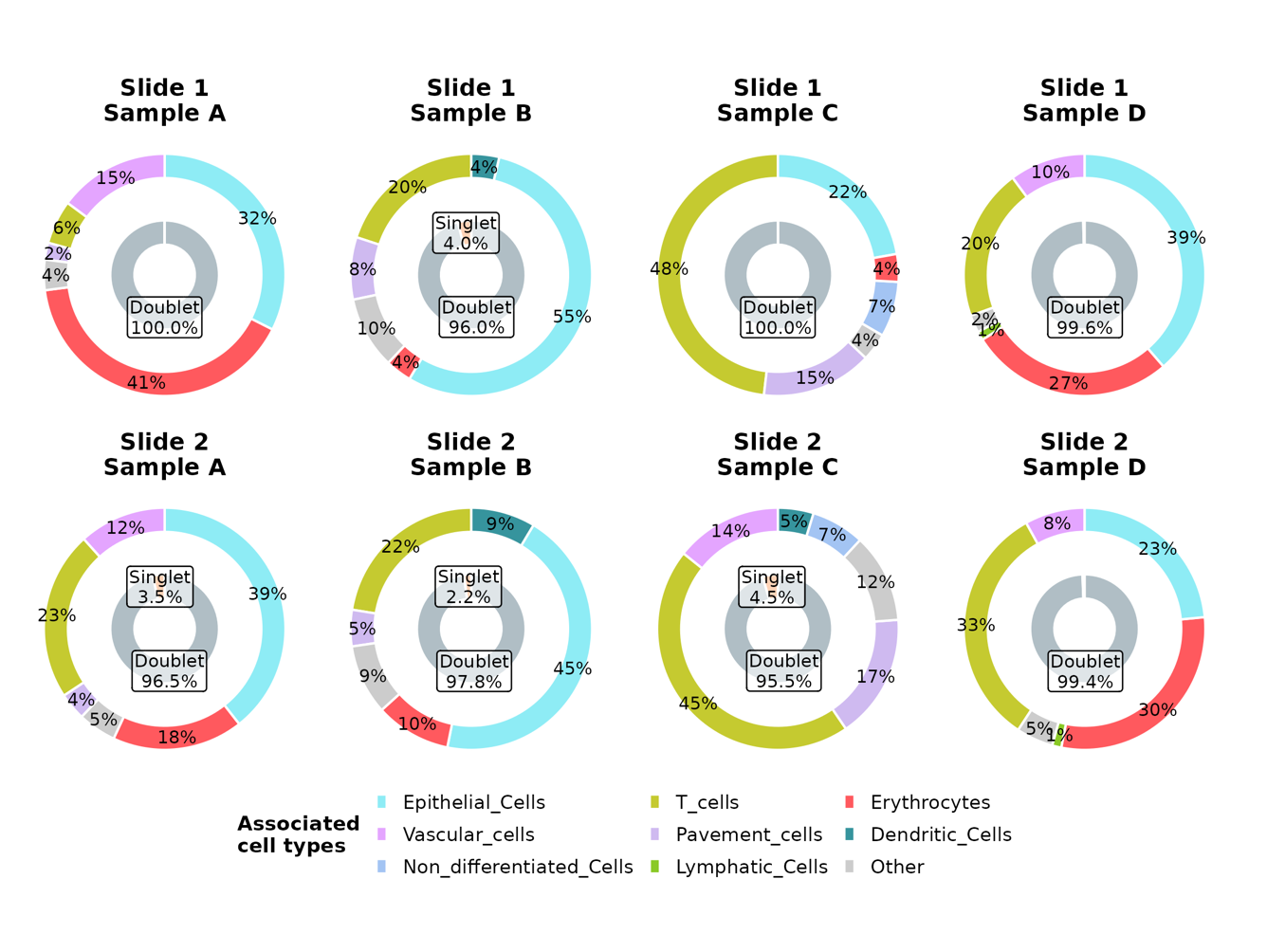

Figure S12.5.1 Distribution of goblet cell singlets and doublet associations across eight gill samples. The inner ring of each doughnut chart represents the proportion of goblet cell-containing spots classified as singlets (dominated by goblet cells alone) versus doublets (goblet cells co-localising with another cell type). The outer ring shows the distribution of associated partner cell types within the doublet fraction. Only the top five most abundant associated cell types are displayed per sample; all other associations are collapsed into the “Other” category. Colours for each cell type correspond to the guide legend. Percentages are calculated relative to all goblet cell-containing spots in each sample.

Across doublet associations, goblet cells most often paired with epithelial cells, T cells, erythrocytes, and vascular cells, with smaller contributions from pavement and dendritic cells depending on the sample (Figure, outer ring). In Sample A, goblet cells were most frequently associated with erythrocytes (18 - 41%) and epithelial cells (32 - 39%), followed by vascular cells (12 - 15%), T cells (6 - 23%) and pavement cells (2 - 4%). In Sample B, pairings were dominated by epithelial cells (45 - 55%) and T cells (20 - 22%), with additional contributions from pavement (5 - 8%), dendritic (4 - 9%), and erythrocytes (4 - 10%). In Sample C, goblet cells most frequently partnered with T cells (45 - 48%), alongside epithelial cells (22%), pavement cells (15 - 17%), vascular cells (14%), and smaller proportions of non-differentiated (7%) and dendritic cells (5%). In Sample D, associations were more evenly distributed between T cells (20 - 33%), erythrocytes (27 – 30%), epithelial cells (23 – 39%), and vascular cells (8 - 10%), with minor contributions from lymphatic cells (1%).

Across samples, observed goblet-cell partner ranges were: epithelial 22–55%, T cells 6–48%, erythrocytes 4–41%, vascular 8–15%, pavement 2–17%, dendritic 4–9%, lymphatic up to 1% and non-differentiated 7% (Figure 1.93). See Figures 1.84–1.85 for spatial context.
